# Two Y chromosome-encoded genes determine sex in kiwifruit

**DOI:** 10.1101/615666

**Authors:** Takashi Akagi, Sarah M. Pilkington, Erika Varkonyi-Gasic, Isabelle M. Henry, Shigeo S. Sugano, Minori Sonoda, Alana Firl, Mark A. McNeilage, Mikaela J. Douglas, Tianchi Wang, Ria Rebstock, Charlotte Voogd, Paul Datson, Andrew C. Allan, Kenji Beppu, Ikuo Kataoka, Ryutaro Tao

**Affiliations:** Graduate School of Agriculture, Kyoto University, Kyoto 606-8502, Japan; JST, PRESTO, Kawaguchi-shi, Saitama 332-0012, Japan; The New Zealand Institute for Plant and Food Research Limited (PFR), Private Bag 92169, Auckland 1142, New Zealand; Department of Plant Biology and Genome Center, University of California Davis, CA, USA; R-GIRO, Ritsumeikan University, Shiga 525-8577, Japan; School of Biological Sciences, University of Auckland, Private Bag 92019, Auckland 1142, New Zealand; Faculty of Agriculture, Kagawa University, Miki, Kagawa 761-0795, Japan

**Author notes:** Graduate School of Environmental and Life Science, Okayama University, 700-8530, Okayama, Japan (Takashi Akagi).

## Abstract

Dioecy, the presence of male and female individuals, has evolved independently in multiple flowering plant lineages. Although theoretical models for the evolution of dioecy, such as the “two-mutation” model, are well established, little is known about the specific genes determining sex and their evolutionary history. Kiwifruit, a major tree crop consumed worldwide, is a dioecious species. In kiwifruit, we had previously identified a Y-encoded sex-determinant candidate gene acting as the suppressor of feminization (SuF), named *Shy Girl* (*SyGI*). Here, we identified a second Y-encoded sex-determinant that we named *Friendly boy* (*FrBy*), which exhibits strong expression in tapetal cells. Gene-editing and complementation analyses in *Arabidopsis thaliana* and *Nicotiana tabacum* indicated that *FrBy* acts for the maintenance of male (M) functions, independently of *SyGI*, and that these functions are conserved across angiosperm species. We further characterized the genomic architecture of the small (< 1 Mb) male specific region of the Y-chromosome (MSY), which harbors only two genes significantly expressed in developing gynoecia and androecia, respectively: *SyGI* and *FrBy*. Resequencing of the genome of a natural hermaphrodite kiwifruit revealed that this individual is genetically male but carries deletion(s) of parts of the Y-chromosome, including *SyGI*. Additionally, expression of *FrBy* in female kiwifruit resulted in hermaphrodite plants. These results clearly indicate that Y-encoded *SyGI* and *FrBy* act independently as the SuF and M factors in kiwifruit, respectively, and provide insight into the evolutionary path leading to a two-factor sex determination system but also a new breeding approach for dioecious species.

## MAIN TEXT

In flowering plants, hermaphroditism is ancestral and most common, but a minority of plant species have evolved separate sexes (dioecy), in a lineage-specific manner (1–3). Similar to mammals, sexuality in plants is often determined by a heterogametic male system with XY chromosomes, where the Y chromosome is thought to carry one or two male-determining factors (1, 3, 4, 5). Previous analyses of Y chromosome evolution in plants, initially in *Silene latifolia* and *Carica papaya*, have revealed long non-recombining male-specific regions, which encompass many genes (6–8), although the sex-determining genes have not been fully characterized. Recently, Y chromosome-encoded sex determinants have been identified in persimmons (*Diospyros* spp.) and garden asparagus (*Asparagus officinalis*) (9, 10). In persimmons, the Y-encoded pseudogene *OGI* encodes a small-RNA targeting its autosomal counterpart gene, *MeGI*. To the best of our understanding, *OGI* is sufficient for expression of maleness and repression of female development (3, 9, 11). In garden asparagus, two Y-encoded factors, named *SOFF* and *aspTDF*, act independently to suppress gynoecium and promote androecium development, respectively (10). The asparagus observation is consistent with a previously proposed theoretical framework, called the “two-mutation model” (12, 13), while the persimmon case is not. This model proposes that evolution from an ancestral hermaphrodite could occur if females carried a mutated (non-functional) version of a male promoting factor (M) on the proto-X chromosome, resulting in establishment of gynodioecy. A second mutation, a gain-of function suppressor of feminization (SuF) on the proto-Y chromosome would then establish males. Together, these sex-determining mutations may, if closely linked, define a genome region resembling an XY chromosome pair or sex-linked genome region (13). Still, the evolutionary pathways governing the transitions into dioecy are poorly understood because only a few examples have been characterized to date.

Kiwifruit, a major fruit crop consumed worldwide, belongs to the genus *Actinidia*, in which most species are dioecious (14). The sexuality in kiwifruit is genetically controlled by a heterogametic male system (i.e., XY system). A Y-encoded cytokinin response regulator, named *Shy Girl* (*SyGI*), acts as one of the two putative sex determinants, the suppressor of female development (Su^F^) (15). The other sex determinant, the putative male promoting factor (M) has not been identified. Although the establishment of SuF in *Actinidia* is estimated to have occurred approximately 20-mya (15) and predated the divergence of the *Actinidia* species, kiwifruit still carries incipient and homomorphic sex chromosomes, including a small sex-determining region (15–17). Recent breeding has derived some hermaphroditic and neuter individuals in *A. deliciosa*, which constitute an additional resource for the identification of a second sex determinant in kiwifruit. Here, we attempted to identify the male promoting sex determinant (M factor), by fine assessment of the genes located on the male-specific region of the Y-chromosome (MSY), and identification of genes differentially expressed between male and female at an early stage of tapetum differentiation. The function of the M factor was validated in model plants and in kiwifruit, resulting in the first development of an artificial hermaphrodite crop from a dioecious individual. Finally, we further assessed the evolution of the two sex determinants on the Y-chromosome, unveiling transitions into and out of dioecy in kiwifruit.

Previously, genomic sequencing reads from F1 sibling trees derived from an interspecific cross, *A. rufa* sel. Fuchu × *A. chinensis* sel. FCM1 were used to identify and assemble the potential male specific region of the Y chromosome (MSY) of *A. chinensis* (15). The 249 resulting contigs, totaling approximately 0.5Mb in length, contained Y-specific sequences with perfect co-segregation with the plants’ sex, and included 61 hypothetical genes (15). In kiwifruit, androecia differentiation between males and females is observed during tapetum degeneration (stage 3-4 in Fig. S1) (15, 18, 19). To identify candidate male promoting (M) factors within the MSY, we conducted mRNA-Seq analyses on developing anthers (5 males and 5 females) before tapetum degeneration (“stage 1-2” in Fig. S1) from the F1 population described above. The mRNA-Seq reads were mapped to the 61 hypothetical candidate genes. Only one of them exhibited male-specific expression (RPKM > 1). This gene included a fasciclin domain, which are typically involved in cell adhesion (Table S1). This fasciclin-like gene was named “*Friendly Boy* (*FrBy*)”, as a potential counterpart of the Su^F^ sex determinant, *Shy Girl* (*SyGI*). *FrBy* is nested within the monophyletic MTR1 family (Fig. 1a). In rice, MTR1 contributes to tapetum degradation via programmed cell death (PCD), resulting in male fertility (20). The kiwifruit *FrBy* and its orthologs in *Nicotiana tabacum* (FAS1 domain protein), *Arabidopsis thaliana* (AT1G30800) and rice (*Oriza sativa*, MTR1), showed no significant differentiation according to site-branch-specific evolutionary rate analysis against the other branches (Fig. 1b), suggesting conserved protein function. The presence of the *FrBy* gene was male-specific in a wide variety of *Actinidia* species (Fig. 1c). The expression of *FrBy* was specific to early developing androecia (Fig. 1d-e). Tapetum cell-specific qRT-PCR using laser capture microdissection (Supplemental Figure S2) and *in situ* RNA hybridization analysis (Fig. 1f-h) both indicated that *FrBy* expression in androecia was confined to tapetal cells in stage 1-2 and possibly to meiocyte or tetrads. This is consistent with previous observations of MTR1 in rice (Tan et al. 2012) and with its putative function to contribute to tapetum degradation following PCD in kiwifruit (Supplemental Fig. S1 and S3) (19). Differentially expressed genes (DEGs) in male and female anthers (Supplemental Table S2, Figure S4) were also consistent with the potential function of *FrBy*. We identified 538 DEGs (FDR < 0.1) when analyzing transcriptome data from developing anthers (as described above). In those 538 DEGs, GO terms involving PCD and phosphorylation signals were highly enriched (Supplemental Table S3). Not only PCD, but abundant phosphorylation signals are indispensable for proper tapetum maturing and degradation (21–23). Furthermore, an ortholog of *Tapetal Development and Function 1* (*TDF1*) or *MYB35*, a key gene in tapetum maturation in Arabidopsis (Zhu et al. 2008) and one of the two sex determinants in dioecious garden asparagus (10, 24, 25), was detected as one of the male-biased DEGs in kiwifruit, although this gene (Acc30672.1) was not located within the MSY (Supplementary Table S2, Figure S5).

**Figure 1:**
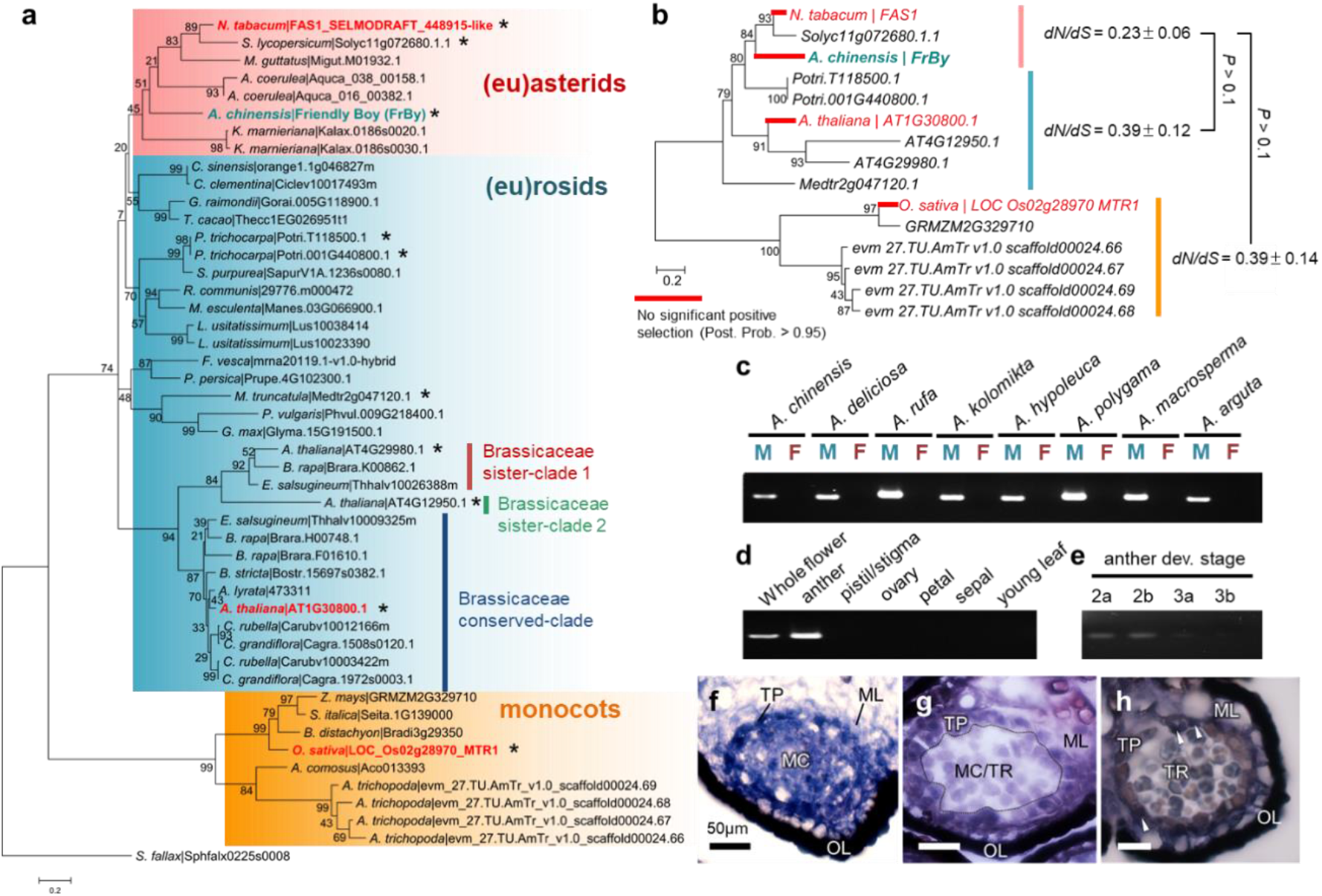
Identification of the *fasciclin*-like *FrBy* as the candidate of the M factor. **a**, Phylogenetic tree of fasciclin-like genes from 32 angiosperm species, including the kiwifruit *FrBy* (light green). The orthologs from (eu)asterids, (eu)rosids, and monocots constitute three putatively monophyletic clades, shown in red, blue and yellow, respectively. The family Brassicaceae forms three subclades, although only one of them is conserved across the Brasicaceae species. The gene functions of the orthologs written in red have been validated previously (for MTR1 in rice) or in this study (for AT1G30800 for Arabidopsis, FAS1 for *Nicotiana tabacum*). The orthologs with asterisks were used for further evolutionary analysis in panel (**b**). **b**, pairwise evolutionary rates (*dN/dS*) in each clade suggested no protein functional differentiation between these orthologs. No site-branch specific positive selection was detected in red branches, suggesting that protein function and key domains were conserved in MTR1, AT1G30800, FAS1, and kiwifruit FrBy. **c**, PCR analysis specific to *FrBy*. Male-specificity of *FrBy* is conserved in a wide variety of *Actinidia* species. M: male, F: female. **d-e**, expression pattern of *FrBy*. (**d**), *FrBy* is expressed specifically in anthers and not on other organs. Within the anthers (**e**), *FrBy* is substantially expressed in stage 2a-b, and faintly at stage 3a. **f-h**, RNA *in situ* hybridization using antisense *FrBy* probe. *FrBy* is expressed highly in tapetum cells (TP) and meiocyte (MC) at stage 2a (**f**), in tapetum cells only at stage 2b (**g**), and faintly in tapetum cells (shown by arrows) at stage 3a (**h**). These results of RNA in situ hybridization were consistent with tapetum-specific expression analysis using laser capture microdissection (Supplemental Figure S2).

To investigate the function of *FrBy*, we first used the CRISPR/Cas9 gene editing system in two distantly related model plants, *Arabidopsis thaliana* and *Nicotiana tabacum* (Supplemental Table S4). Although the Arabidopsis genome includes three paralogs of *FrBy*, only one, AT1G30800, was in the cluster which was conserved across the *Brassicaceae* species (Fig. 1a). In Arabidopsis, the AT1G30800-null lines (Supplemental Figure S6) were self-sterile, with low pollen germination rates (Fig. 2g), but could successfully produce seed after being crossed to control male plants (Fig. 2a-e). The null line showed substantial delay in tapetal layer degradation (Supplemental Fig. S7), which is consistent with the development of female kiwifruit plants (Fig. S1) (18, 19). On the other hand, the lack of AT1G30800 had no significant effect on female reproductive function (*P* > 0.1, Supplemental Fig. S8). In *N. tabacum*, knock-out mutation of the *FrBy* ortholog, *FAS1* (*fas1*) (Supplemental Figure S9) resulted in male sterility, with substantial reduction in pollen germination rate, and was accompanied by a delay in tapetum degradation. The other organs, including the gynoecium, showed no differentiation compared to the control plants (Fig. 2h-o). The transgenic *N. tabacum* lines expressing the kiwifruit Su^F^ gene, *SyGI*, under the control of native promoter (p*SyGI-SyGI*) exhibited female-sterility (15). Reciprocal crossing using control plants, p*SyGI-SyGI*, and *fas1* indicated that *SyGI* and *FAS1* independently promote gynoecium and androecium development, respectively (Fig. 2p). Importantly, male function in a *fas1* null line could be complemented by introduction of the kiwifruit *FrBy* under the control of its native promoter (Fig. 2q-r, Supplemental Figure S10), indicating that *FrBy* can act to maintain male fertility via proper tapetum degradation in *N. tabacum*. These results all suggest that *FrBy* is likely to be the male promoting factor, and that the two sex determining genes, *SyGI* and *FrBy* work independently for female and male fertility, respectively, in kiwifruit. Furthermore, our phylogenetic and evolutionary analyses indicated that the function of this fasciclin-like monophyletic gene is highly conserved across angiosperm species.

**Figure 2:**
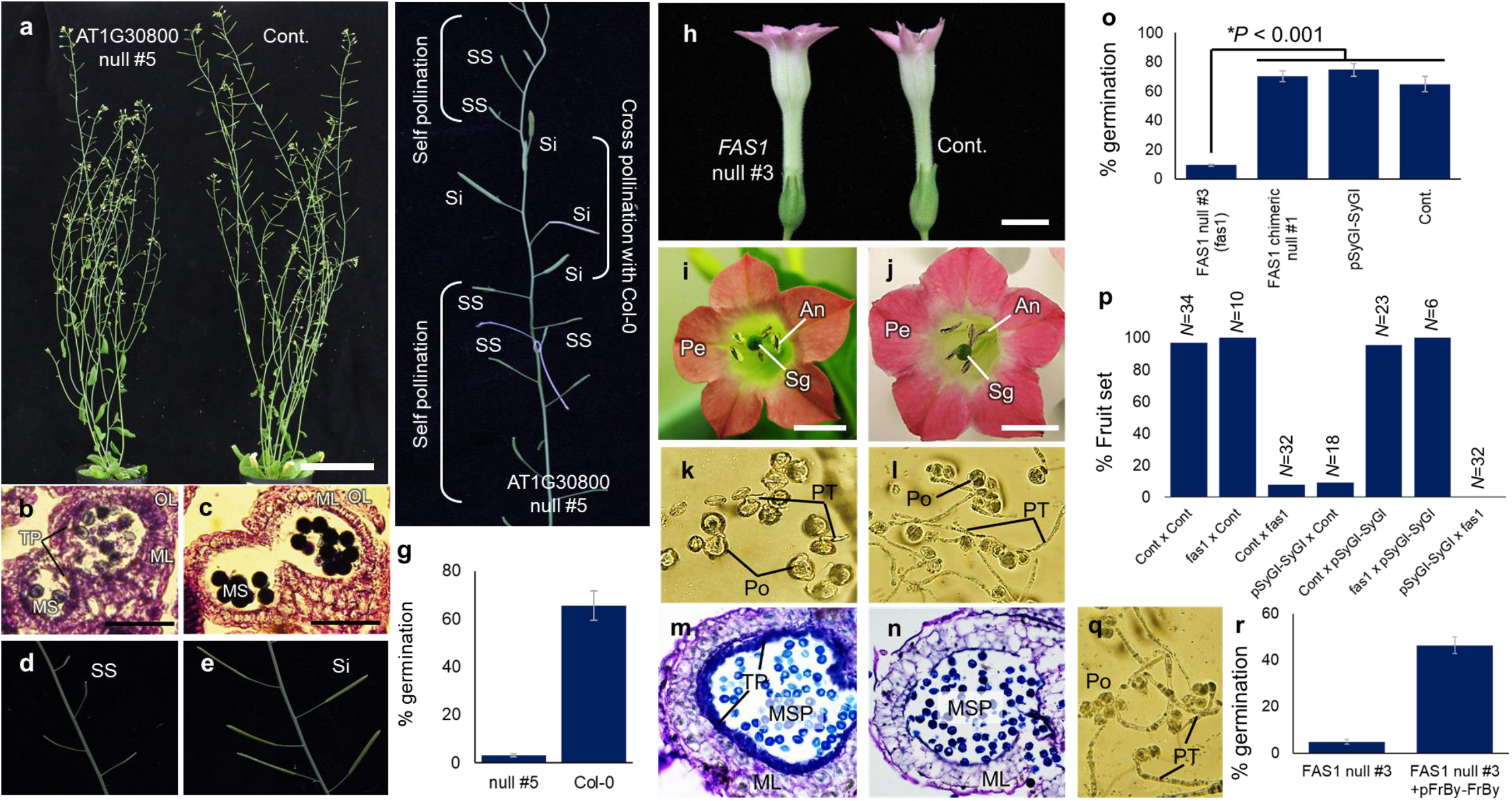
Functional validation of *FrBy* in two model plants. **a-g**, A knock-out mutation in the *FrBy* ortholog, *AT1G30800*, resulted in male sterility in Arabidopsis. **a**, phenotype of the *AT1G30800* null mutant #5 and control lines. Developing anthers showed no tapetum degradation in the AT1G30800 null mutant (**b**), while controls showed complete degeneration of tapetum cells and matured microspore (MS) (**c**). TP: tapetum cells, OL: outer layer, ML: middle layer. **d-e**, reduced seed production in the *AT1G30800* null mutant (**d**) and control (**e**). SS: sterile silique bearing no seeds, Si: silique. **f**, silique development after self- and cross-pollination of the *AT1G30800* null mutant line #5. Only cross pollination with Col-0 control plants resulted in seeds production. **g**, pollen germination ratios in *AT1G30800* null mutant #5 and #2, and control plants (Col-0). Null mutant #5 mostly lost the ability to produce fertile pollen, and chimeric null mutant #3 showed substantial reduction in pollen fertility. **h-p**, In *N. tabacum*, knock-out mutants of the *FrBy* ortholog, *FAS1*, showed male sterility. Whole plant (**h**) and flower organs (**i** for the *FAS1* mutant and **j** for control) phenotypes were not different between the *FAS1* null mutant and control lines. An: anther, Sg: stigma, Pe: petal. Pollen germination ability was substantially reduced in the *FAS1* null mutant (**k**), in comparison to the control lines (**l**). Po: pollen grain, PT: pollen tube. Thick tapetum layers were observed after anther dissection in the microspore stage of the *FAS1* null mutant (**m**), while control lines showed complete degradation of tapetum cells (**n**). In comparison to the *FAS1* null mutant, the SuF gene, *SyGI*, had no significant effect on pollen germination rate (**o**). Reciprocal crosses with control, *FAS1* null, and *SyGI*-expressing lines indicated that *FAS1* and *SyGI* act only for male and female functions, respectively (**p**). **q-r**, complementation of male function in *FAS1* null, using the kiwifruit *FrBy* gene under the control of its native promoter (p*FrBy-FrBy*). Observation of pollen germination (**q**) and germination ratio (**r**) in *FAS1* null mutants transformed with p*FrBy-FrBy*.

Reference genome sequences for kiwifruit have been assembled from female (2A+XX) cultivars (26, 27) but the Y chromosome of kiwifruit (or *Actinidia* spp.) has not been sequenced to date. Here, we constructed the whole genome reference sequence of a male cultivar, Soyu, which is one of the main pollinizers used in Japan. The sequences were assembled using 10X Genomics Supernova v1.2.2, which is based on a long haploblocking method suitable for assembly of highly heterozygous diploid genomes (28). Downstream genomic analysis for the Soyu cultivar was conducted on the “pseudohaploid” version of the whole genome assembly. The drafted genome sequence covered ca 710Mb which corresponds to 94% of the estimated size of kiwifruit genome (758 Mb) (Hopping, 1994, Huang 2013), with N50 = *ca* 318kb for scaffolds (Supplemental Table S5). The genomic short-read sequences were generated from large DNA molecules partitioned and barcoded using the Gel Bead in Emulsion (GEM) microfluidic method of 10X Genomics (28). Thus, the information present in the GEM barcodes anchored to the assembled contigs reflect their physical distance, in which proximal contig pairs often share the same GEM barcodes. GEM barcodes were extracted from read pairs and linkage information was assigned using a custom script. To construct longer scaffolds within the MSY, we first identified 9 scaffolds, which together spanned 1.43 Mb, and contained ca 87% of the Y-specific contigs previously assembled (15) (Supplemental Table S6). To organize these scaffolds relative to each other, we applied DelMapper (6), an approach that employed the traveling salesman problem (TSP), used in radiation hybrid mapping of mammalian chromosomes (29, 30) or deletion mapping of Y-chromosome in dioecious *Silene latifolia* (6). Using this method, 8 of the 9 scaffolds were successfully organized. In this new assembled super-scaffold, the two sex determinants, *SyGI* and *FrBy*, are located on adjacent scaffolds, at an estimated distance of ca 500kb (Fig. 3a, Supplemental Figure S11). Mapping of genomic reads from male and female individuals in the KE population (15) indicated that a ~800-kb region enriched in male-specific sequences (putative MSY) was located at the center of this assembled super-scaffold (Fig. 3a). The putative MSY includes the two sex determinants and highly repetitive sequences, which is consistent with the structure of MSY in other plants or animals (10, 31). The putative pseudoautosomal region (PAR) appears to be single copy and exhibited no substantial gender-bias, and mostly flanked the putative MSY, although some PAR-like sequences were also located inside the putative MSY (Fig. 3a). In this super-scaffold, 145 genes were predicted using AUGUSTUS (32), and 30 of these genes were fully male-specific (Fig. 3b-c, Supplemental Table S7). Of the 30 male-specific genes, only *SyGI* and *FrBy* were substantially expressed (RPKM > 1) in carpel and anther, respectively (Fig. 3d-e, Supplemental Table S8), based on transcriptome information from the KE population (Akagi et al. 2018 for carpel) (15). These data support the hypothesis that they are the two factors determining sexuality in kiwifruit.

**Figure 3:**
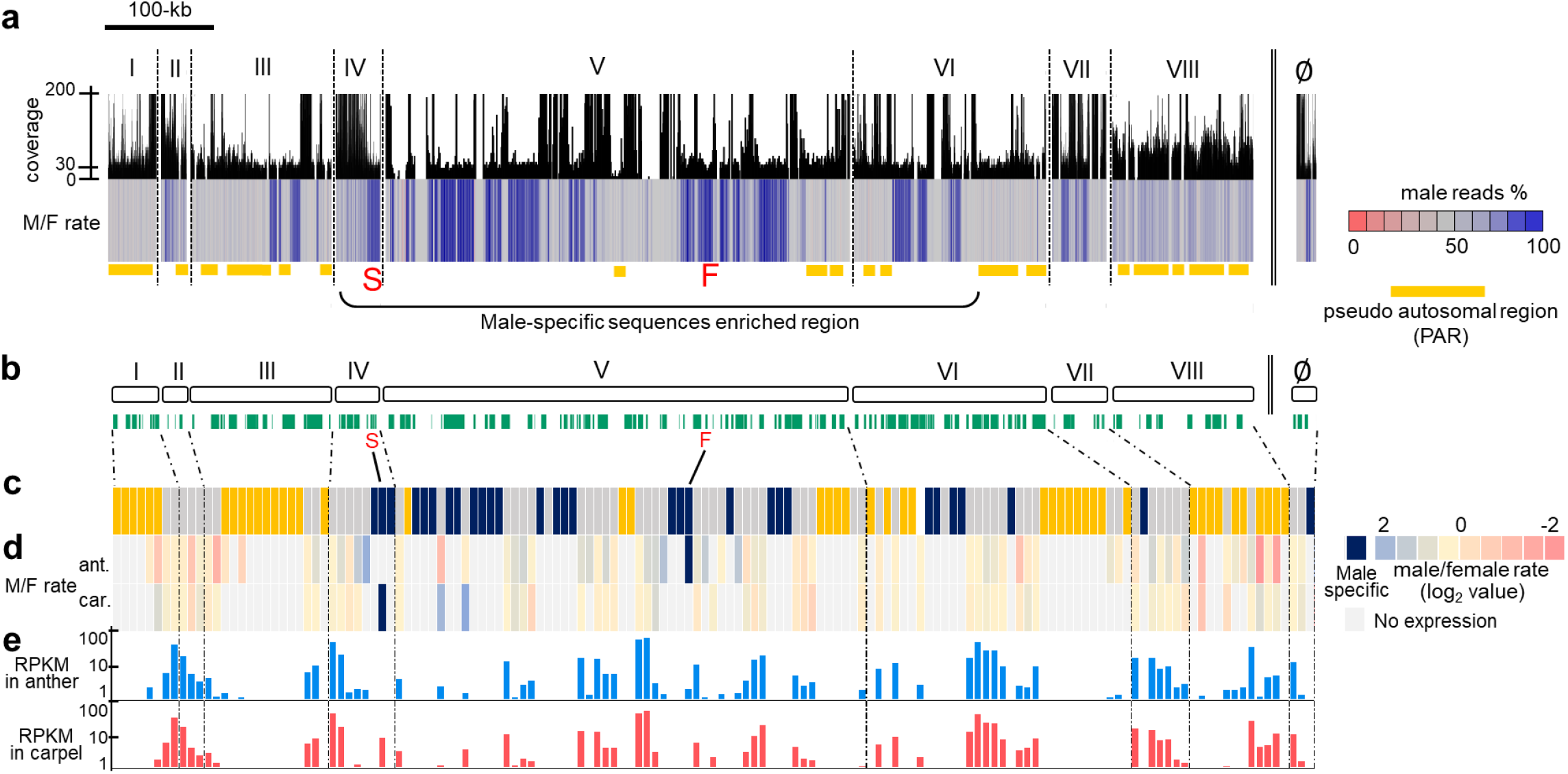
Sequence architecture of the kiwifruit Y chromosome, including the two sex determinants. **a**, Super-scaffold constituted of anchored 8 contigs (I-VIII) and unanchored one (∅), with coverage of male genomic read (bars) and percentage of male genomic reads (color chart). The coverage from simplex alleles at single sites (or male-specific region with no X-chromosome counterpart) is estimated at 20-23X. Repeated regions, such as transposable elements, were identified as those with high coverage (>50X). Putative pseudo-autosomal regions (PAR) were defined as non-repetitive regions with unbiased reads coverage (ca 25-75% male reads), and highlighted with orange bars. *SyGI* (S) and *FrBy* (F) were located on male-specific regions in the middle of the Y-chromosomal contigs. **b**, models of the nine contigs (white bars) and the genes predicted in these contigs (green bars). **c**, male-specificity in the predicted genes in each contig. Genes in male-specific, PAR, and repetitive regions, are shown in blue, orange, and gray, respectively. **d-e**, expression patterns of the predicted genes in each contig. **d**, expression bias between male and female individuals, in anthers (ant.) and carpels (car.). Complete male-specific expression was given in thick blue. **e**, expression levels (RPKM) in anther and carpel. Across the Y-specific genes as given in deep blue, only *FrBy* and *SyGI* were substantially expressed (RPKM > 1) in differentiating anther and carpel, respectively.

We further corroborated the role of these two sex determinants in two ways. Breeding programs have generated a few hermaphrodite accessions in hexaploid *A. deliciosa* (33). We sequenced one of them, the KH-line (Supplementary Table S9), which was thought to be a Y-dependent (or possibly X-dependent) hermaphrodite (6A+XXXXXY^h^ or 6A+XXXXXX^h^, Supplementary Figure S12). Mapping of the KH and control *A. deliciosa* [male cv. Matua (6A+XXXXXY)] genomic reads to the Y-chromosomal scaffolds described above (Fig. 3) demonstrated that the KH-line carries the *FrBy* gene, but not the *SyGI* gene, either through one or several long deletion(s), including the loss of *SyGI*, or through gain of *FrBy* on the X chromosome from recombination with the Y chromosome (Figure 4a, Supplementary Figure S13). Consistent with this result, *SyGI* could not be amplified in the KH-line, while *FrBy* could (Figure 4b). This suggested that loss of *SyGI* (the Su^F^) in the Y, or gain of *FrBy* in the X, resulted in a natural hermaphrodite line (Figure 4c). Next, we set out to develop hermaphrodite kiwifruit artificially, as well as to further validate the *FrBy* function. We introduced the *FrBy* ORF under the control of its native promoter (p*FrBy-FrBy*) into a “rapid flowering” *A. chinensis* female cv. Hort16A. In this line, the *CENTRORADIALIS* (*CEN*) genes have been truncated by gene-editing, resulting in lines that bypass the long juvenile phase (*ca* 3-4 years) and flower precociously (34). As anticipated, the p*FrBy-FrBy* lines, which flowered 4 months after regeneration, were hermaphroditic, exhibiting restored androecium function. These produced fruits including fertile seeds after self-pollination (Fig. 4d-h, Supplementary Table S10). Pollen tubes from the p*FrBy-FrBy* lines grew similarly to those from male accessions, in contrast to the control lines (Figure 4g-i, Supplementary Figure S14). These results clearly indicated that *FrBy* acts as the M factor in kiwifruit sex determination. They also provide valuable insight into new breeding approaches for dioecious species.

**Figure 4:**
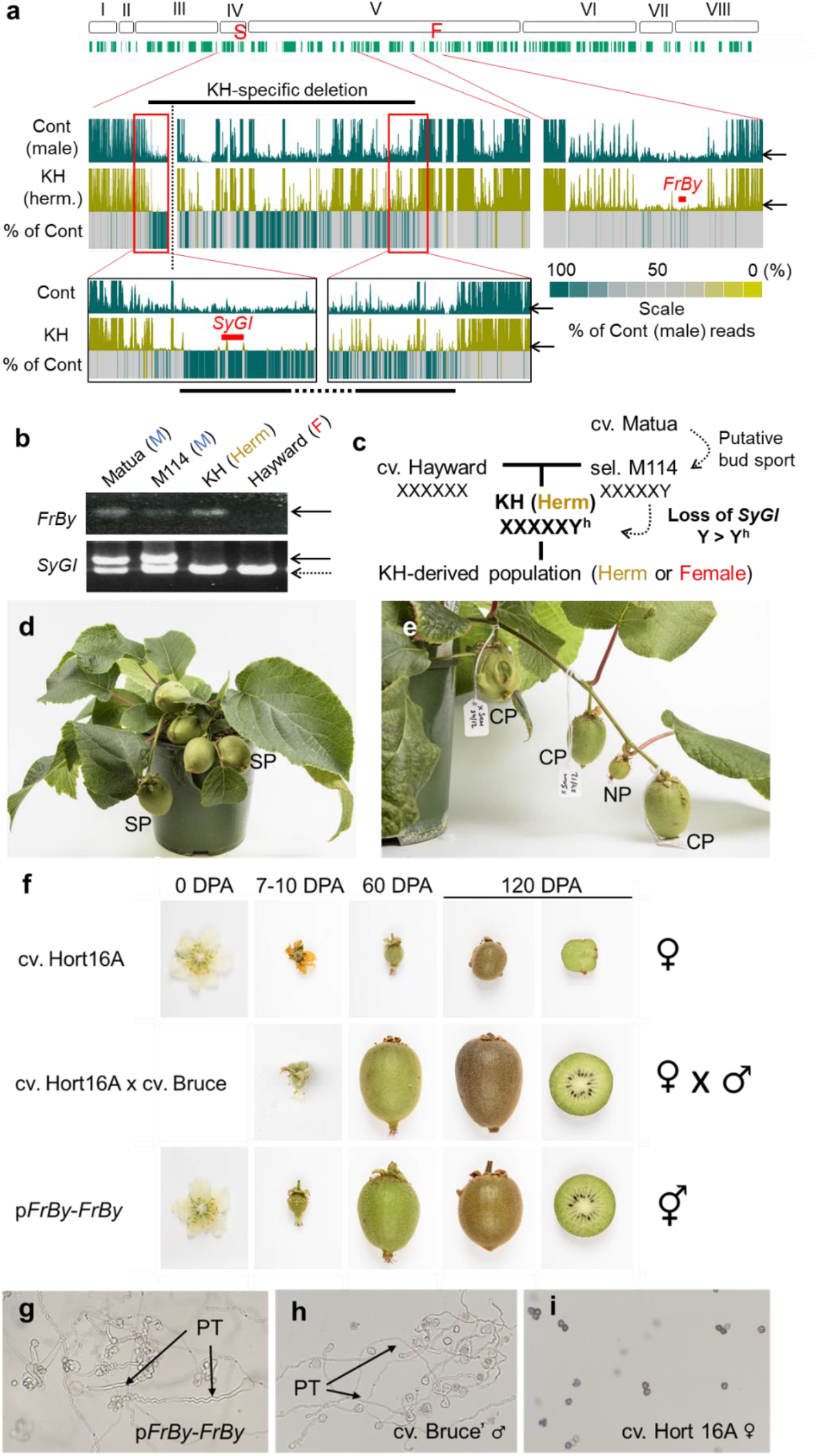
Lost of *SyGI*, or gain of *FrBy* resulted in a natural and synthetic hermaphrodite kiwifruit, respectively. **a**, Genomic context in the Y-chromosomal contigs, in male cv. Matua (control), and a natural hermaphrodite line, KH. The KH line carries long deletions across contigs IV-V, including *SyGI*.This was confirmed by PCR analyses (Supplemental Figure S13). The right arm of contig V was conserved in the KH line. Arrows indicated estimated coverage of simplex Y allele. **b**, conservation of *FrBy* and *SyGI* in KH and its parental cultivars/lines, based on the pedigree of the KH lines (as shown in **c**). Solid arrows indicated *FrBy* and *SyGI*. Dotted arrow showed a paralog of *SyGI* with high sequence homology located on an autosome (Akagi et al. 2018). M: male, F: female. *SyGI* is absent specifically in the KH line, while *FrBy* was conserved in both parental males and the KH lines. **c**, pedigree of the KH line and model for the establishment of the *SyGI* null Y^h^ chromosome. **d-i**, development of hermaphrodite kiwifruit by transformation with *FrBy* under the native promoter (p*FrBy-FrBy*). **d**, The p*FrBy-FrBy* lines showed fertilized fruits with self-pollination (SP) in 120 days post anthesis (DPA). **e**, Control female cv. Hort16A showed unfertilized fruit in non-pollination (NP), while cross-pollination with p*FrBy-FrBy* pollen (CP) resulted in fertilized fruit. **f**, In control female cv. Hort16A (top panels), flowers aborted or developed small parthenocarpic fruit without seeds. In cross pollination of control female and a male cv. Bruce (middle), normal fruits bearing fertile seeds were developed on pollinated cv. Hort16A. In self-pollination of transgenic cv. Hort 16A with p*FrBy-FrBy* (bottom), normal fruits with fertile seeds were developed, comparable to normal female x male. Pollen grains from pFrBy-FrBy-induced female cv. Hort16A (**g**) had the ability to grow pollen tubes (PT), as well as from male cv. Bruce (**h**), while pollen grains from control female cv. Hort16A were sterile (**i**).

Taken together, our results are consistent with the following evolutionary path for the transition from hermaphroditism to dioecy in *Actinidia*, based on two tightly linked genes within a small MSY: loss-of-function of *FrBy* established a proto-X chromosome, while lineage-specific gain-of-function in *SyGI* (15) derived a dominant suppressor of gynoecium development, establishing a proto-Y chromosome (Figure 5). This evolutionary process and the predicted function of the determinants are consistent with the “two-mutation model” (12, 13). This proposed evolutionary history of *SyGI* and *FrBy* is also consistent with those of the two-locus type sex determinants in *Asparagus* or *Phoenix* (10, 35), although the specific function of these sex-determinants are different. The putative SuF genes, *SOFF* for *Asparagus* and *LOG1*-like for *Phoenix*, were established by lineage-specific gene duplication/translocation on the Y chromosome; while the putative M genes, *MYB35* (*TDF1*) for *Asparagus* and *CPY703/GPAT3* for *Phoenix* were lost from the X chromosomes. Within the order Ericales, kiwifruit (*Actinidia*) evolved these Y-encoded sex determinants while persimmons (*Diospyros*), evolved a single sex determinant on the Y chromosome, *OGI*, which encodes small-RNAs repressing an autosomal feminizing gene, *MeGI* (9, 36). Despite their different specific functions, *SyGI* in kiwifruit and *OGI* in persimmon both act as dominant suppressors and were both derived from lineage-specific duplications, suggesting evolutionary consistency in how the diverse sex determination systems have evolved in angiosperms.

**Figure 5:**
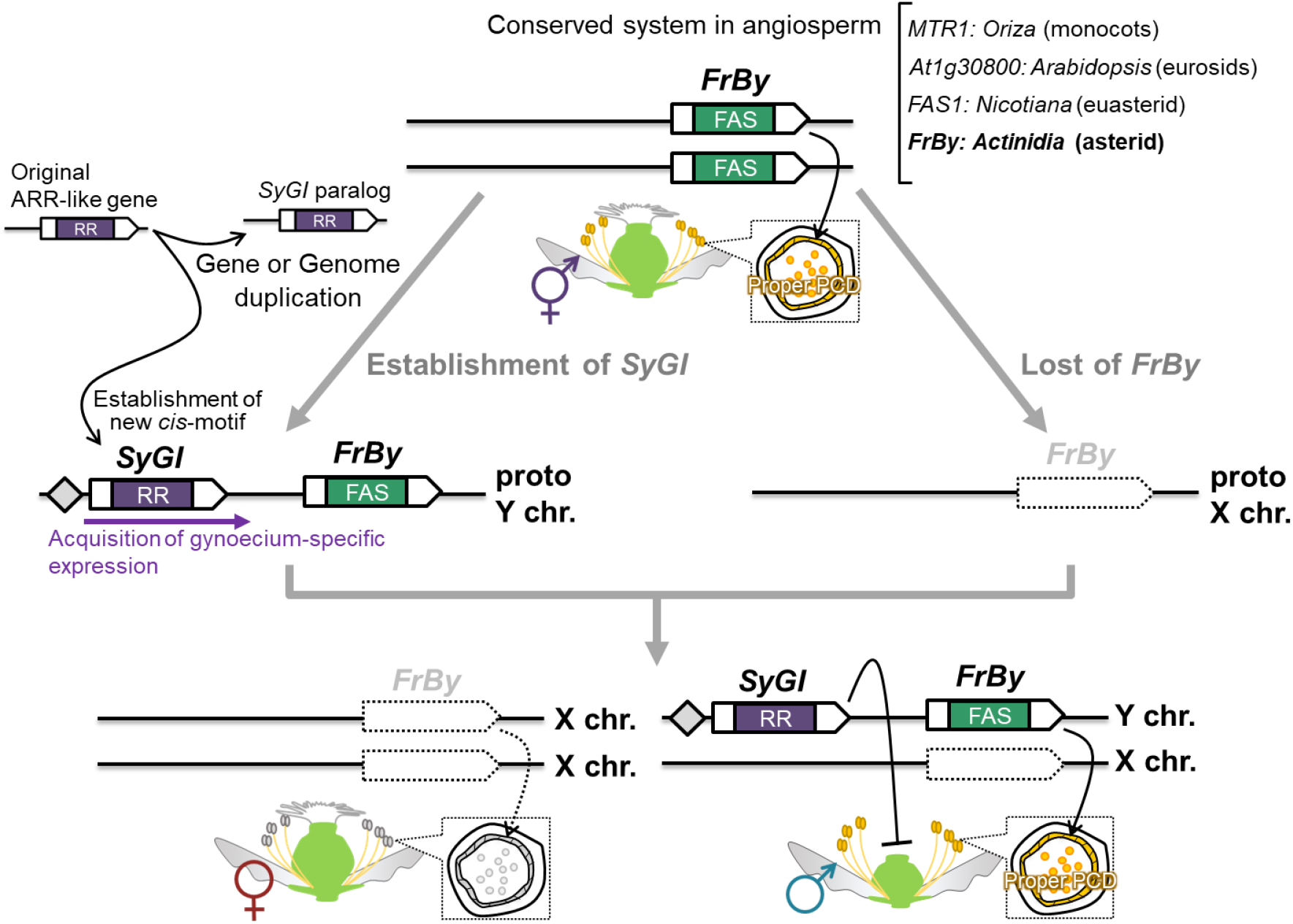
Evolutionary model for the establishment dioecy in *Actinidia*. The putative hermaphrodite ancestor bears an intact *FrBy* ortholog, which carries a function that is conserved across angiosperm. The loss of function in *FrBy* generated a proto X-chromosome, while the gain of function in *SyGI* (15), which occurred next to the intact copy of *FrBy*, resulted in a proto Y-chromosome. Together, these proto X and Y chromosomes evolved to derive the current XY system in the *Actinidia* genus.

## Supporting information

Supplementary Mateials

## SUPPLEMENTARY INFORMATION

### Materials and Methods (S1-S11)

**Method S1:** Screening of the expressed candidate sex determinants

**Method S2:** Expression profiling in kiwifruit anthers

**Method S3:** *in situ* RNA hybridization

**Method S4:** Phylogenetic analysis and detection of positive selection

**Method S5:** 10X Genomics (Chromium) library construction and genome assembly with Supernova

**Method S6:** Anchoring and characterization of the scaffolds surrounding the two sex determinants

**Method S7:** Breeding of hermaphrodite kiwifruit

**Method S8:** Genomic analysis of hermaphrodite kiwifruit

**Method S9:** Construction of vectors

**Method S10:** Transformation

**Method S11:** Floral phenotyping in transformed plants

### Accession numbers

**Supplemental Figures (S1-S14)**

**Supplemental Figure S1:** Transition of anther and definition of the developmental stage

**Supplemental Figure S2:** Tapetum cells-specific *FrBy* expression pattern with laser capture microdissection (LCM).

**Supplemental Figure S3:** Visualization of programmed cell death in tapetum cells of male and female kiwifruit

**Supplemental Figure S4:** Gene expression pattern in male and female anthers

**Supplemental Figure S5:** Phylogenetic analysis of TDF1/MYB35-like genes in *Actinidia*

**Supplemental Figure S6:** Gene disruption of *AT1G30800* in Arabidopsis using CIRISPR/Cas9

**Supplemental Figure S7:** Characterization of anther development in *AT1G30800* null line

**Supplemental Figure S8:** Silique and seed production in *AT1G30800* null line

**Supplemental Figure S9:** Gene disruption of *FAS1* in *Nicotiana tabacum* using CRISPR/Cas9

**Supplemental Figure S10:** Complementation of male function in *FAS1* null lines by kiwifruit *FrBy*

**Supplemental Figure S11:** Correspondence between GEM barcodes and the anchored scaffolds

**Supplemental Figure S12:** Segregation of *FrBy*, in a KH-derived segregating populations.

**Supplemental Figure S13:** Characterization of the putative KH-specific deletion

**Supplemental Figure S14:** Cross pollination with p*FrBy-FrBy*-induced cv. Hort 16A

### Supplemental Tables (S1-S12)

**Supplemental Table S1:** 61 hypothetical genes identified in the MSY contigs and their expression levels in anther

**Supplemental Table S2:** List of DEGs between male and female anthers

**Supplemental Table S3:** GO enrichment analysis in the DEGs between male and female anthers

**Supplemental Table S4:** Characterization of gene-edited *Arabidopsis* and *N. tabacum* lines

**Supplemental Table S5:** Summary of the draft genome assembly in cv. Soyu by 10X Genomics data

**Supplemental Table S6:** List of the assembled scaffolds of cv. Soyu anchored by Y-specific contigs

**Supplemental Table S7:** List of genes predicted in the 9 scaffolds anchored by the Y-specific contigs

**Supplemental Table S8:** Expression patterns of the genes predicted in the 9 scaffolds anchored by the Y-specific contigs

**Supplemental Table S9:** Pedigree of the KH line and the segregating population derived from the KH line

**Supplemental Table S10:** Phenotypic characterization in p*FrBy-FrBy*-induced kiwifruit.

**Supplemental Table S11:** List of plant materials.

**Supplemental Table S12:** List of primers used in this study.

## ACKNOWLEDGEMENTS

We thank Dr. Luca Comai (UC Davis Dept. Plant Biology and Genome Center) for technical advice and bioinformatics support, Drs. Yusuke Kazama and Kotaro Ishii (Riken Institute) for technical support for using the DelMapper program, and Niels Nieuwenhuizen and Jane (Lei) Zhang for vector construction. The KE population were originally provided from Kagawa Prefectural Agricultural Experiment Station. Some of this work was performed at the Vincent J. Coates Genomics Sequencing Laboratory at UC Berkeley, supported by NIH S10 OD018174 Instrumentation Grant. This work was supported by PRESTO Grant Number JPMJPR15Q1 (to TA) and JPMJPR15Q6 (to SSS) from the Japan Science and Technology Agency (JST), by a Grant-in-Aid for Scientific Research on Innovative Areas No. J16H06471 (to TA) from JSPS, and by the National Science Foundation (NSF) IOS award under Grant No. 1457230 (to IMH)

## AUTHOR CONTRIBUTION

TA, IK, and RT conceived the study. TA designed the experiments. TA, SMP, EV, SSS, MS, AF, MJD, TW, RR and CV conducted the experiments. TA, SMP, EV, SSS, MS, IMH, and AF analyzed the data, SMP, MAM, PD, ACA, KB, and IK initiated/bred and maintained the plant materials. TA, SMP, EV, and IMH drafted the manuscript. All authors approved the manuscript.

## AUTHOR INFORMATION

All sequence data generated in the context of this manuscript has been deposited in the appropriate DDBJ database: Illumina reads for gDNAseq and mRNAseq in the Short Read Archives (SRA) database (SRA IDs), the genomic contig sets constructed with 10X Genomics reads were submitted to Genbank (IDs).

Reprints and permissions information is available at www.nature.com/reprints.

The authors declare no competing financial interests.

Correspondence and requests for materials should be addressed to Takashi Akagi.

