## Supplementary Mateials for "Two Y chromosome-encoded genes determine sex in kiwifruit"

*Corresponding author

#### Methods

##### Method S1: Screening of the expressed candidate sex determinants

Developing anthers at stage 1-2, which correspond to the differentiation stage of male or female androecium (see Supplementary Figure S1), were sampled from F1 sibling vines derived from an interspecific cross, *A. rufa* sel. Fuchu × *A. chinensis* sel. FCM1, named KE population (15), planted on Kagawa University, Japan (N34.28, E134.13), in 10-22 April in 2016-2017. Total RNA was extracted using the Plant RNA Reagent (Invitrogen) and purified by phenol/chloroform extraction. Two micrograms of total RNA were processed in preparation for Illumina Sequencing, according to a previous report (15). The constructed libraries were sequenced on Illumina's HiSeq 4000 sequencer (50-bp single-end or 150-bp paired-end reads). All Illumina sequencing were conducted at the Vincent J. Coates Genomics Sequencing Laboratory at UC Berkeley, and the raw sequencing reads were processed using custom Python scripts developed in the Comai laboratory and available online (http://comailab.genomecenter.ucdavis.edu/index.php/), as previously described (9). Male-specific Y-chromosomal sequences in kiwifruit, defined MSY contigs, were comprehensively identified in previous study (15). The mRNA-Seq reads from each 5 male and female individuals from the KE population (Supplemental Table S11) (15) were used to identify the genes substantially expressed in developing anthers. The mRNA-Seq reads were aligned to the hypothetical 61 genes located on the 249 MSY contigs (Akagi et al. 2018), using the Burrows–Wheeler Aligner (BWA) (37) allowing up to ca 3% mismatches. The number of reads mapping to each contigs was recorded from the alignment file produced by the Sequence Alignment/Map (SAM) tool (38) (http://samtools.sourceforge.net/). For *Friendly Boy* (*FrBy*), which showed male-specific and anther-enriched expression, the expression patterns were further examined using various plant organs and developing anthers (stage 2a, 2b, 3a, and 3b, see Supplementary Figure S1).

##### Method S2: Expression profiling in kiwifruit anther

The described mRNA-Seq reads from each 5 male and female individuals of the KE population were aligned to the whole CDS sequences sets in *A. chinensis* (27), using BWA with default parameters. The number of reads mapped to each reference sequences was recorded from the alignment file produced by the Sequence Alignment/Map (SAM) tool (38) (http://samtools.sourceforge.net/). The read counts per gene were generated from the aligned SAM files using a custom R script. Differential expression between male and female individuals was analysed in R (version 3.0.1) using the R package DESeq (Anders and Huber, 2010) (version 1.14; http://bioconductor.org/packages/release/bioc/html/DESeq.html). We conducted DESeq analysis using 5 biological replicates from male and female individuals, with the following parameters: method='per-condition' and sharingMode='maximum'. An FDR threshold of 0.1 was used to identify differentially expressed genes.

##### Method S3: *in situ* RNA hybridization

RNA *in situ* hybridization was conducted as previously described (15). Briefly, developing anthers (stage 2-3), were fixed in FAA (1.8% formaldehyde, 5% acetic acid, 50% ethanol), subsequently displaced by 20% sucrose solution before being sliced by a cryostat (Leica CM1520) using cryofilm (Kawamoto 2003). The tissues were sliced into 10-μm sections, and mounted on FRONTIER coated glass slides (Matsunami Glass Ind., Kishiwada, Japan). The rehydrated tissue sections were incubated in a Proteinase K solution (700 U/mL Proteinase K) for 30 min at 37°C, followed by acetylation for 10 min. Full length *FrBy* cDNA sequences were cloned into the pGEM-T Easy vector (Promega, WI, USA) to synthesize DIG-labelled antisense RNA probes using the DIG-labelling RNA synthesis kit (Roche, Switzerland). The probe solution including RNaseOUT (Thermo Fisher Scientific, USA) was applied to the slides and covered with parafilm. Hybridization was performed at 48°C for >16 h. For detection, 0.1% Anti-Digoxigenin-AP Fab fragments (Sigma-Aldrich) was used as the secondary antibody to stain with NBT/BCIP solutions.

**Method S4: Phylogenetic analysis and detection of positive selection**

The fasciclin-like full-length orthologs of *Friendly Boy* were extracted from the genomes of 31 angiosperm using BLASTp ($<1e^{-10}$) in Phytozome (JGI release version 12.0, https://phytozome.jgi.doe.gov/pz/portal.html), and in the genome database of *Nicotiana* from Sol Genomics Network (https://solgenomics.net/). A single ortholog of the *FrBy* in *Sphagnum fallax*, Sphfalx0225s0008, was used as the outgroup gene. Alignment analyses on amino acid sequences were conducted using MAFFT ver. 7 with L-INS-i model, followed by manual revision using SeaView ver. 4. The evolutionary topology was examined using the maximum likelihood method (ML) by MEGAX (39) with WAG+G model and 1,000 replications for bootstraps. All sites, including missing and gap data, were used for the construction of phylogenic trees, and the nearest neighbour interchange technique was used. Bootstraps were shown on the branches as 1/10 of the calculated values. The orthologous genes in *Oryza sativa*, *Zea mays*, *Amborella trichopoda*, *Populus trichocarpa*, *Medicago truncatula*, *A. thaliana*, *Nicotiana tabacum*, *Solanum lycopersicum*, were subjected to in-codon frame alignment using Pal2Nal (40) and MAFFT ver. 7. Based on the aligned nucleotide sequences, pairwise evolutionary rates (*dN/dS*) were detected by using DnaSP (41). The alignments were also used to estimate evolutionary topology using ML method by Mega, with GTR (+G) model with 1,000 replications for bootstrap. Based on these alignments and topology, the codon-based detection of branch- and site-specific positive selection test was performed using PAML (42). The statistical significance of positive selection on the foreground branches was evaluated using the likelihood ratio test of the null hypothesis that $dN/dS = 1$. Site-specific positive selection was assessed by Empirical Bayes analysis.

**Method S5: 10X Genomics (Chromium) library construction and genome assembly with Supernova**

Genomic DNA of kiwifruit cv. Soyu (*A. chienensis*) was extracted with Genomic-tip 100/G (Qiagen), and then was adjusted to a concentration of 1.0 ng/μl and 1.25 ng of template gDNA was loaded on a Chromium Genome Chip. Whole genome sequencing libraries were prepared using Chromium Genome Library & Gel Bead Kit v.2 (10X Genomics, cat. 120258), Chromium Genome Chip Kit v.2 (10X Genomics, cat. 120257), Chromium i7 Multiplex Kit (10X Genomics, cat. 120262) and Chromium controller according to manufacturer's instructions, with just one modification. Briefly, gDNA was combined with Master Mix, a library of Genome Gel Beads, and partitioning oil to create Gel Bead-in-Emulsions (GEMs) on the Chromium Genome Chip. The GEMs were isothermally amplified with primers containing an Illumina Read 1 sequencing primer, a unique 16-bp 10x barcode and a 6-bp random primer sequence. Next, bar-coded DNA fragments were recovered for Illumina library construction. The amount and fragment size of post-GEM DNA was quantified using a Bioanalyzer 2100 with an Agilent High sensitivity DNA kit (Agilent, cat. 5067-4626). Prior to Illumina library construction, the GEM amplification product was sheared on an E220 Focused Ultrasonicator (Covaris, Woburn, MA) to approximately 350bp (55 seconds at peak power = 175, duty factor = 10, and cycle/burst = 200). Next, the sheared GEMs were converted to a sequencing library following the 10X standard operating procedure and using the following 4 indexed adaptors: TACTCTTC, CCTGTGCG, GGACACGT, ATGAGAAA. The library was quantified by qPCR with a Kapa Library Quant kit (Kapa Biosystems-Roche) and sequenced on one lane of HiSeq4000 sequencer (Illumina, San Diego, CA) with paired-end 150 bp reads. The 10X barcoded reads were demultiplexed and assembled using the Supernova 2.1.1 pipeline as described previously (28). A number of assembly versions were generated, on average resulting in assemblies that were 700Mb in length with an N50 of 290kb.

**Method S6: Anchoring and characterization of the scaffolds surrounding the two sex determinants**

The assembled Supernova scaffolds were anchored with the 249 genomic MSY contigs derived from male *A. chinensis* sel. FCM1 (15), which comprehensively covered Y-specific sequences by kmer cataloging in segregated populations (9, 15), by using blastn (>99% nucleotide homology, in 100-bp window). To rule out the possibility that X-allelic scaffolds were anchored, we further mapped the genomic Illumina reads from male and female individuals in the KE population (15), to detect male-specific polymorphisms throughout the scaffolds. The 10X Genomics indexes (called "gem") sequences of the anchored 9 scaffolds, to which ca 87% of the MSY contigs were anchored (Supplemental Table S6), were subjected to DelMapper program (6), which utilizes the traveling salesman problem (TSP) method, according to Kazama et al. 2016. The sequences of the 9 anchored scaffolds were assessed by AUGUSTUS (32) to identify putative genes located on the scaffolds. The mRNA-Seq reads from each 5 male and female individuals of the KE F1 population (15) in anthers (stage 1-2), described above, in carpels (stage 1), and in whole flowers (stage 1-2) (15) were aligned to these predicted genes by using BWA with default parameters, to calculate expression levels. The male-specificity in genomic sequences were assessed with rate of the mapped reads from male and female. gDNA-Seq reads from each 20 male and female individuals of the KE population (15) were aligned into the 9 scaffolds, by using BWA allowing 10% mismatches, to calculate the rate of male/female reads coverages. The non-repetitive regions or genes with <66.6% male reads (coverage rate in male/female < 2) and with >95% male reads (coverage rate in male/female > 19) were defined as pseudo autosomal region (PAR) and MSY, respectively, in this study.

**Method S7: Breeding of hermaphrodite kiwifruit**

The seven plants tested were from three types of cross: inconstant male x inconstant male; inconstant males selfed; and hermaphrodite x inconstant male. The inconstant male KFM, which came from a cross between the female cv. Hayward and in inconstant male M114 collected from a grower's orchard, was used as a pollen parent and, when selfed, also as an ovule parent. The inconstant male M121, also collected from a grower's orchard, was used as an ovule parent. The hermaphrodite KH, which also came from the cv. Hayward x M114 cross (i.e. is full-sib of KFM), was used as an ovule parent. The hermaphrodites KH, o01_02 (from M121 x KFM) and s01_03 (from KFM selfed) are considered to be de novo hermaphrodites from recombinant gametes produced by inconstant males. The male cv. Matua was tested as control. For both selfing and crossing, flowering shoots on ovule parents were enclosed within paperbags before anthesis. Inconstant male and hermaphrodite ovule parents had all bisexual flowers within bags emasculated and inconstant males also had all staminate flowers removed. Anthers were collected from bisexual flowers of the pollen parent KFM and dried overnight in covered petri dishes at 25°C to induce dehiscence. Pollen was collected into labelled vials. At anthesis, bags on flowering shoots of ovule parents were briefly removed and flowers were hand-pollinated using small brushes. Bags were replaced and retained for several weeks after pollination. After fruit harvest, seeds were extracted using a pectinase/cellulase solution to degrade the fruit flesh, then sown in small trays of seed raising mix. After 2 weeks stratification at 5°C and 2 weeks of alternating temperatures, trays were placed on a heat-bed. Seedlings were potted up and grown on over the winter in a glasshouse with 16 hour 25°C-days maintained by use of lights and heating units. Seedlings were hardened off in a shade-house before being planted out in T-bar orchard blocks at the Te Puke Research Centre, Bay of Plenty, New Zealand with six rows per block, 5 m between rows, 96 seedlings per row and 1 m between seedlings within rows. Families were randomly assigned to 6-seedling plots along rows. Seedlings were maintained for seven years, after this only individuals of interest were retained.

**Method S8: Genomic analysis on hermaphrodite kiwifruit**

Genomic DNA was extracted from young kiwifruit leaves using the Qiagen Plant Mini Kit, following the standard protocol. To check the amplification of *SyGI* and *FrBy*, PCR was performed with specific primers (Supplementary Table S12) on using PrimeSTAR® GXL DNA Polymerase, following manufacturer's instructions. The PCR followed a 3-step reaction (95 °C for 30 seconds, 55 °C for 30 seconds, then 72 °C for 30 seconds) for 35 cycles. Approximately 0.5 μg of genomic DNA of sel. KH and male cv. Matua (Supplementary Table S9) was used for the construction of Illumina genomic libraries. The Illumina genomic libraries were constructed using KAPA HyperPlus Library Preparation Kit (KAPA Biosystems), according to the manufacture's instruction. For barcoding, NEXTflex adaptors (Bioo Scientific) were ligated. To remove contamination of self-ligated adapter dimers, libraries were size-selected using AMPure in 0.8:1 (v/v) AMPure:reaction volume. Eluted DNA was enriched by PCR reaction using Prime STAR Max (Takara) at the following PCR conditions: 30 s at 98°C, 6 cycles of 10 s at 98°C, 30 s at 65°C and 30 s at 72°C and a final extension step of 5 min at 72°C. Enriched libraries were purified with AMPure (0.7:1 v/v AMPure to reaction volume), and quality and quantity were assessed using the Agilent BioAnalyzer (Agilent Technologies) and Qubit fluorometer (Invitrogen). Libraries were sequenced using Illumina's HiSeq4000 (100-bp paired-end reads). The genomic library reads were trimmed as described in trimming of mRNA-Seq reads, and then aligned to the described 9 scaffolds covering the two sex determinants and the MSY, with BWA allowing up to 5% mismatches. The transition of the coverages of the mapped reads were visualized with IGV (43).

**Method S9: Construction of vectors**

For Arabidopsis (*A. thaliana*), pKI1.1R vector (44) was digested by *Aar*I. The ds-oligos, which target At1g30800, were ligated with the linearized vector by TaKaRa Mighty mix (TaKaRa). For *N. tabacum,* pPLV2 was digested by *Bam*HI. Using pKI1.1R vector (44) as the template, gRNA expression cassettes and Cas9 expression cassettes were amplified by PCR with the primers pPVL02_AtU6_infBamF and pPVL02_Hsp_infBamR (see Supplemental Table S12). The linearized pPLV2 and the PCR fragment was conjugated using In-Fusion™ reaction. The vector harboring Cas9 and gRNA expression cassettes in the pPLV2 backbone were designated as pPLV2-GE. pPLV2-GE vector was digested by *Aar*I. The *Aar*I digested products were ligated with the annealed oligos which target the *FAS1* gene. The oligos were described as Supplemental Table S12. For complementation test with *FrBy* in kiwifruit and *N. tabacum,* a sequence comprising 1571 nt of the promoter region and the 723 nt coding region of *Actinidia chinensis FrBy*, followed by the *OCS* 3′ terminator, was synthesized by GenScript (http://www.genscript.com) then cloned into *Apa*I/*Not*I digested pDE-KRS-HYG, derived from pDE-KRS (34). The fragment containing the CaMV *35S* promoter, *hptII* and *35S* terminator was PCR amplified from pHYGREX5 (45) using the forward 35S_NheIF (5'-TTAAGCTAGCGTATTGGCTAGAGCAG-3') and reverse 35S_UTR_R_MauBI_R (5'-TATACGCGCGCGTAATTCGGGGGATCTGGA-3') oligonucleotide primers and cloned into *Nhe*I/*Mau*BI digested pDE-KRS, replacing the *nptII* expression cassette with the *hptII* expression cassette. The resulting plasmid was transformed into *Agrobacterium tumefaciens* strain EHA105 by electroporation.

**Method S10: Transformation**

**1. Arabidopsis**

Arabidopsis (*A. thaliana*) ecotype Columbia-0 (Col-0), *lig4* (SALK_044027), and *ku70* (SALK_123114) was grown under white light (400-750 nm) with continuous light condition at 22°C until transformation. The binary construct was introduced into *Agrobacterium tumefaciens* strain GV3101 by electroporation. Next, Arabidopsis plants were transformed using the flower-dipping method, according to the previous protocol (9). Screening of transgenic plants was conducted by red fluorescence of the T1 seeds using fluorescence stereomicroscope. The genotyping was conducted using the T1 cauline leaves.

**2. *Nicotiana tabacum***

Tobacco plants (*N. tabacum*) cv. Petit Havana SR1 were grown in vitro under white light (400-750nm) with 16-h-light and 8-h-dark cycles at 22°C until transformation. The binary construct was introduced into the A. tumefaciens strain EHA101 using the helper vector pSOUP by electroporation. Young petioles and leaves of tobacco plants were transformed by the leaf disk method. Transgenic plants were selected on Murashige and Skoog medium supplemented with 100 μg/mL kanamycin.

**3. Kiwifruit**

Plant material from a female kiwifruit cultivar cv. Hort16A (*A. chinensis* Planch. var. *chinensis*) was previously edited (Varkonyi-Gasic et al., 2018). This rapid-flowering material was used for further transformation and expression studies. The media were described before (46). *Agrobacterium tumefaciens* mediated transformation of *A. chinensis* was performed as previously described (46), with the exception that hygromycin (15mg/l) was used for selection instead of kanamycin. Briefly, leaf strips excised from *in vitro*-grown shoots were cocultivated with *Agrobacterium* suspension culture and transferred to regeneration and selection medium. Individual calli were excised from the leaf strips for further selection and bud induction, and adventitious buds regenerated from the calli were excised and transferred to shoot elongation medium. When shoots had grown to 1–2 cm high, they were transplanted onto rooting medium. Rooted transgenic plants were then potted and grown in a containment greenhouse at ambient conditions (temperature min 18 °C/ max 30 °C night/day, 14 h/10 h light/dark in summer).

**Method S11: Floral phenotyping in transformed plants**

Developing anthers were fixed in FAA (1.8% formaldehyde, 5% acetic acid, 50% ethanol), subsequently displaced by 10-30% sucrose solution series before being sliced by a cryostat (Leica CM1520) using cryofilm (47). The tissues were sliced into 5-μm sections, and then stained by toluidine blue or Fast Green. The pollens from flesh anthers were then placed on agar slides (1% agar containing 20% sucrose and 0.01% boric acid) at 25 °C for 12 hours, maintaining moisture with damp filter paper. For Arabidopsis and *N. tabacum*, the pollen germination ratio was counted as average percentages in batches of 200 pollen grains from the first five flowers. For kiwifruit crossing using p*FrBy-FrBy*-induced lines, flowers were pollinated at full bloom using a sterile paint brush. Fruit was harvested 100-120 days post anthesis (DPA).

**Accession numbers**

All sequence data generated in the context of this manuscript have been deposited in the appropriate DDBJ database: Illumina reads for gDNA-Seq and mRNA-Seq in the Short Read Archives (SRA) database (DRA accession)

**Supplemental Figure S1**

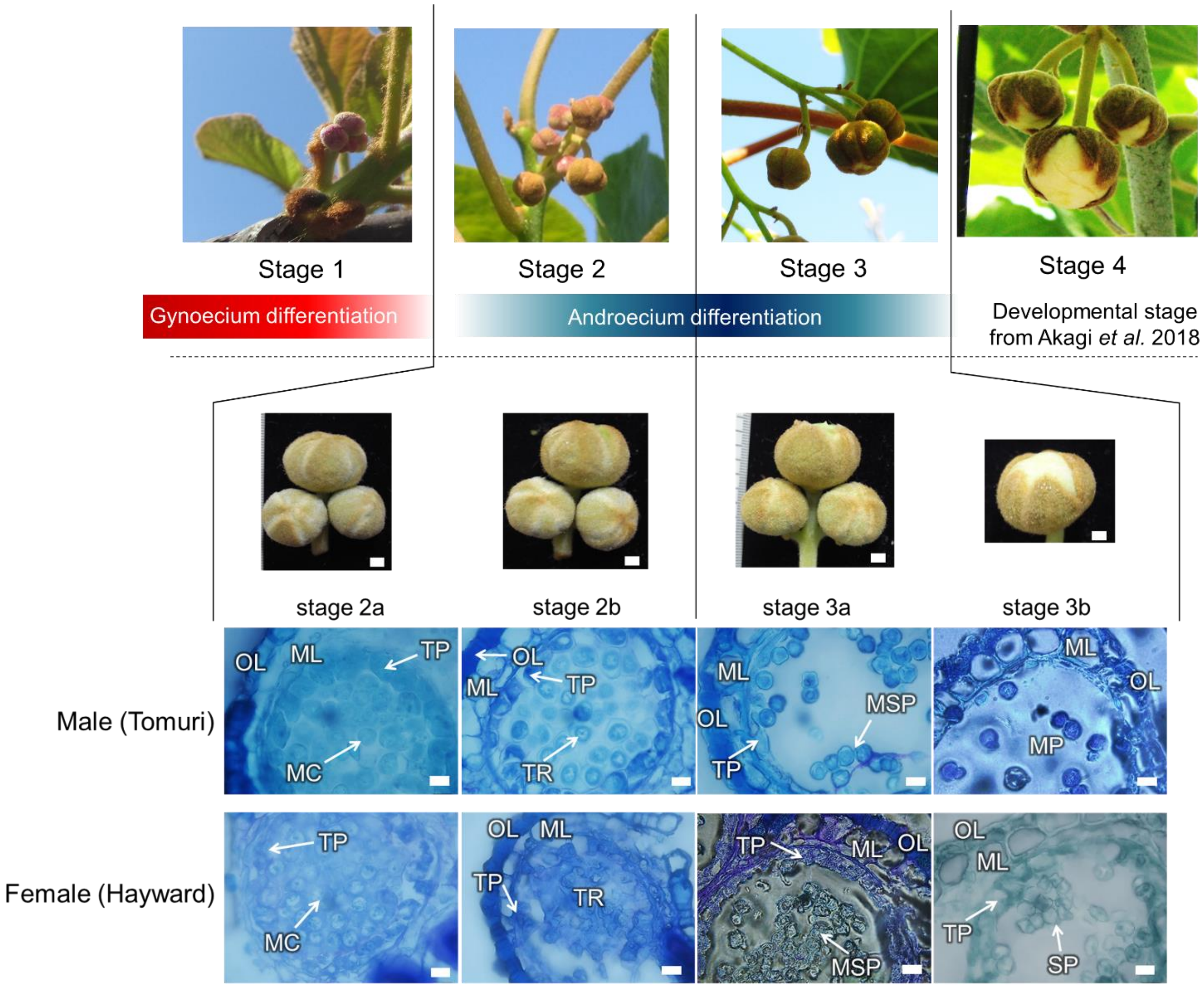


**Supplementary Figure S1 Anther development and definition of the developmental stages**

Kiwifruit flower development has been divided into 4 stages (15). Gynoecium differentiation in male and female is completed in stage 1, while androecium differentiation occurs in stage 2-3. Stages 2 and 3 were further subdivided into stages 2a and 2b, and 3a and 3b, based on anther morphogenesis and especially microspore development. In stage 2a, meiocytes (MC) and tapetal layers (TP) are differentiated. In stage 2b, tetrads (TR) and clear tapetal layers are observed. In stage 3a, microspores (MSP) are separated and tapetal cells degenerate in males only. In stage 3b, tapetal layers completely disappear in male to develop mature pollen (MP), while female maintain thick tapetal layers with sterile pollen (SP). OL: outer layer, ML: middle layer. Bars indicate 20 $\mu$m.

**Supplemental Figure S2**

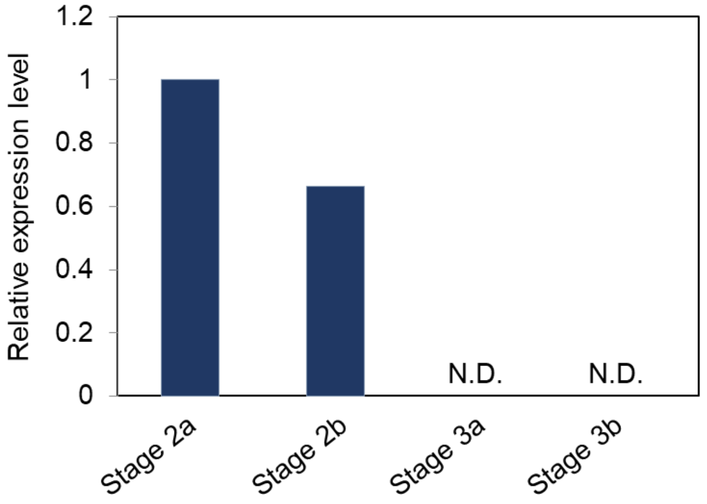


**Supplementary Figure S2 Tapetum cells-specific *FrBy* expression pattern using laser capture microdissection (LCM)**

Tapetum cells in anthers of developmental stages 2a, 2b, 3a, and 3b were isolated with LCM, and the expression levels of *FrBy* were analyzed by qRT-PCR analysis. Consistent with our results from RNA *in situ* hybridization, *FrBy* was expressed specifically in stage 2a-2b. N.D.: not detected with 45-cycles in PCR.

**Supplemental Figure S3**

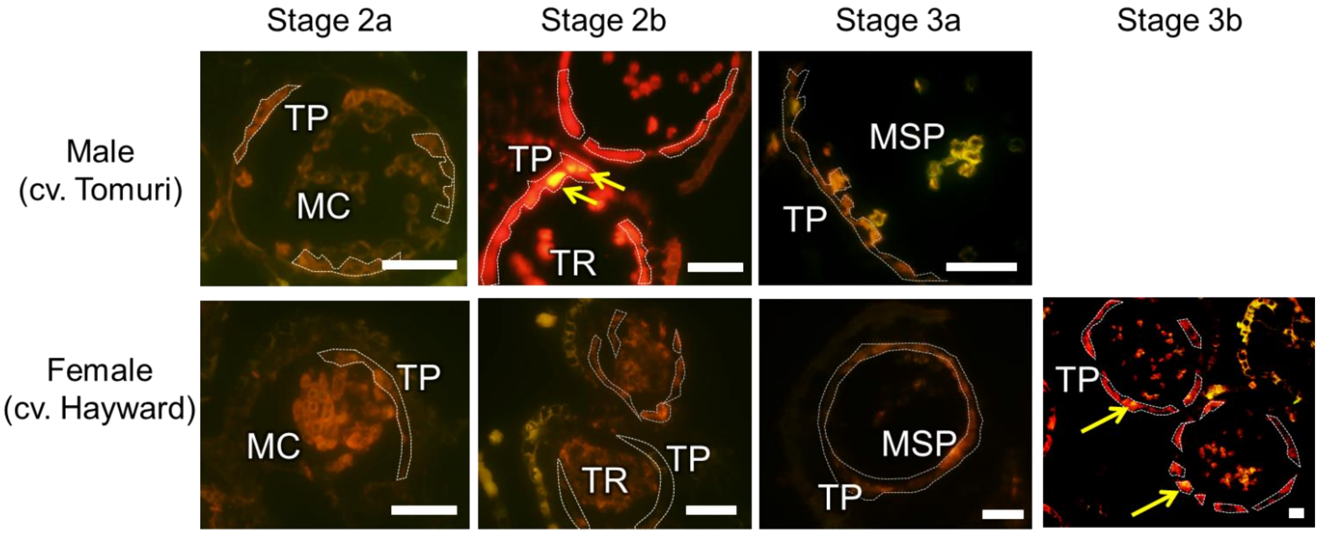


**Supplementary Figure S3 Visualization of programmed cell death in tapetum cells of male and female kiwifruit**

Programmed cell death (PCD) signals were detected with TUNEL method in which PCD signals could be detected by yellow fluorescence. In stage 2b, which is immediately before tapetum degradation (Figure S1), substantial PCD signal was detected in tapetum cells (TP) in males, as shown by yellow arrows. Female tapetum cells rarely showed PCD signals in tapetum cells until stage 3a. In stage 3b, female also showed PCD in the tapetum. MC: meiocyte, TR: tetrads, MSP: microspore.

**Supplemental Figure S4**

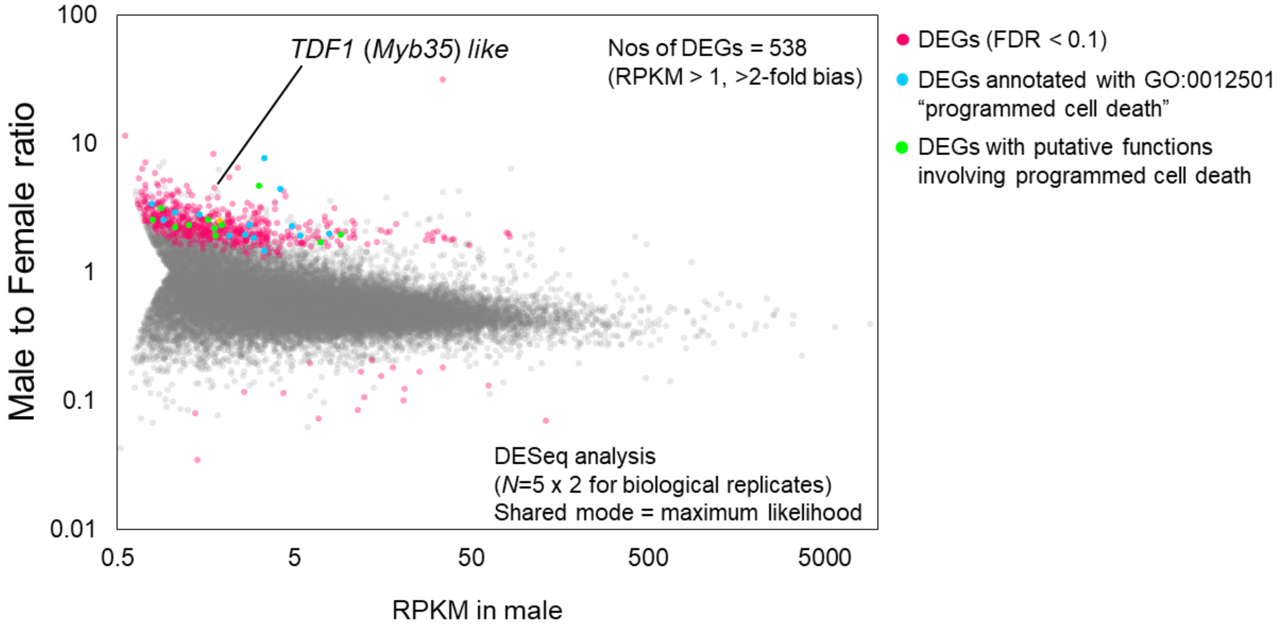


**Supplementary Figure S4 Gene expression pattern in male and female anthers**

Transcriptomic data from male and female anthers in stage 2 were analyzed to identify genes differentially expressed (DEGs) between male and female. Five male and female lines from the KE population (15) were used as biological replicates. The genes were distributed based on expression level (X axis) and the ratio of male/female reads (Y axis). The 538 DEGs (FDR < 0.1 and >2-fold bias between male and female, see Supplementary Table S2) are shown in pink circles, including *TDF1* (*Myb35*)-like gene, with the exception of the DEGs annotated with programmed cell death in GO (GO:0012501) or with programmed cell death-associated functions, which were highlighted in sky blue and light green, respectively.

**Supplemental Figure S5**

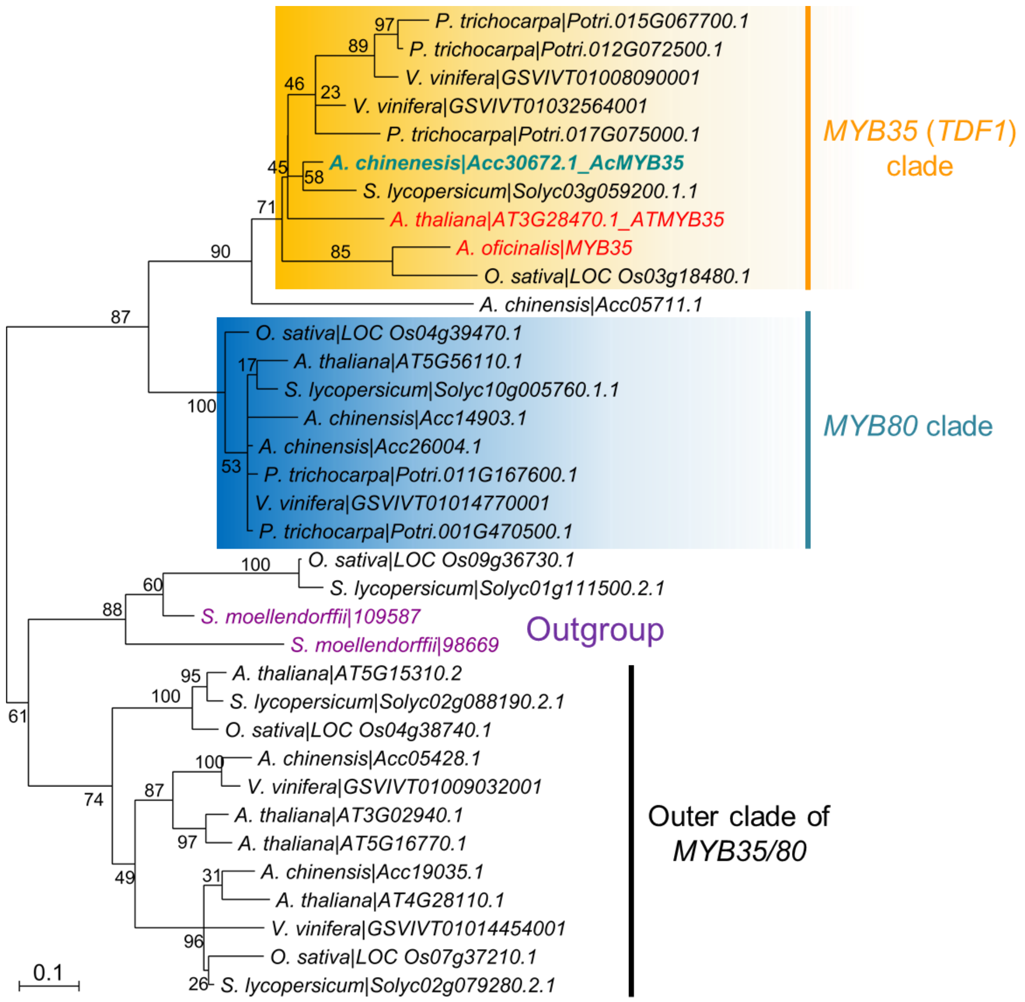


**Supplementary Figure S5 Phylogenetic analysis of *TDF1/MYB35*-like genes in *Actinidia***

Phylogenetic topology for MYB35/MYB80-like genes was constructed using the orthologous sequences from *Actinidia chinensis* (27), Arabidopsis (*Arabidopsis thaliana*), rice (*Oriza sativa*), poplar (*Populus trichocarpa*), tomato (*Solanum lycopersicum*), and grape (*Vitis vinifera*), and the male promoting sex determining gene in garden asparagus (*Asparagus officinalis*), shown in red. Two major clades (MYB35/MYB80) and an outer clade were defined, with *Selaginella* orthologs as an outgroup shown in purple. The *TDF1/MYB35*-like Acc30672.1_*AcMYB35*, a DEG between male and female anthers (Figure S4 and shown in light green here), and the *Asparagus MYB35*, are both nested with the TDF1/MYB35 clade, with significant statistic supports (bootstrap = 71/100).

**Supplemental Figure S6**

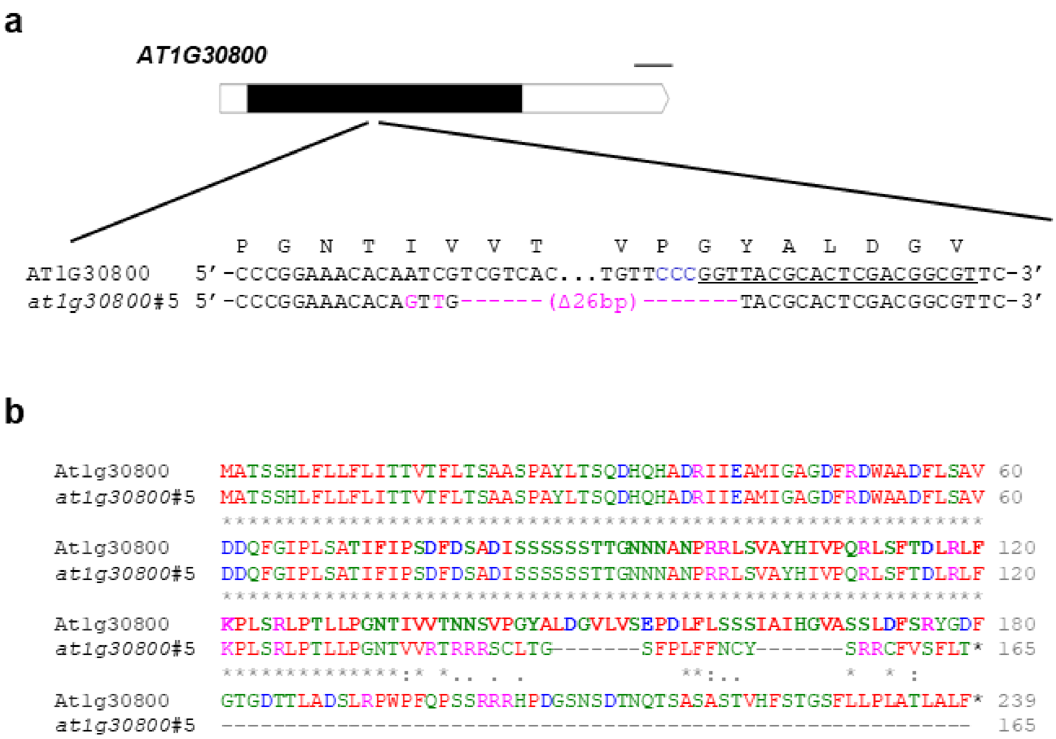


**Supplementary Figure S6 Gene disruption of *AT1G30800* in Arabidopsis using CRISPR/Cas9.**

**a,** Gene model of *AT1G30800* and alignment of the genomic sequence of *AT1G30800* from the genome-edited line #5. (Upper) Filled box: CDS, open box: UTRs, scale bar = 100 bp. (Lower) Blue: PAM sequence, underlined: target sequence of gRNA, magenta: substitution and deletion induced by CRISPR/Cas9. The *AT1G30800* sequence was obtained by an Arabidopsis Col-0 annotation database, Araport11 (https://www.araport.org/). **b,** Sequence alignment of the putative translation product of the edited *at1g30800* from line #5. The putative FAS1 domain (position 71-176, ID:SM00554 from SMART database) is shown in bold.

**Supplemental Figure S7**

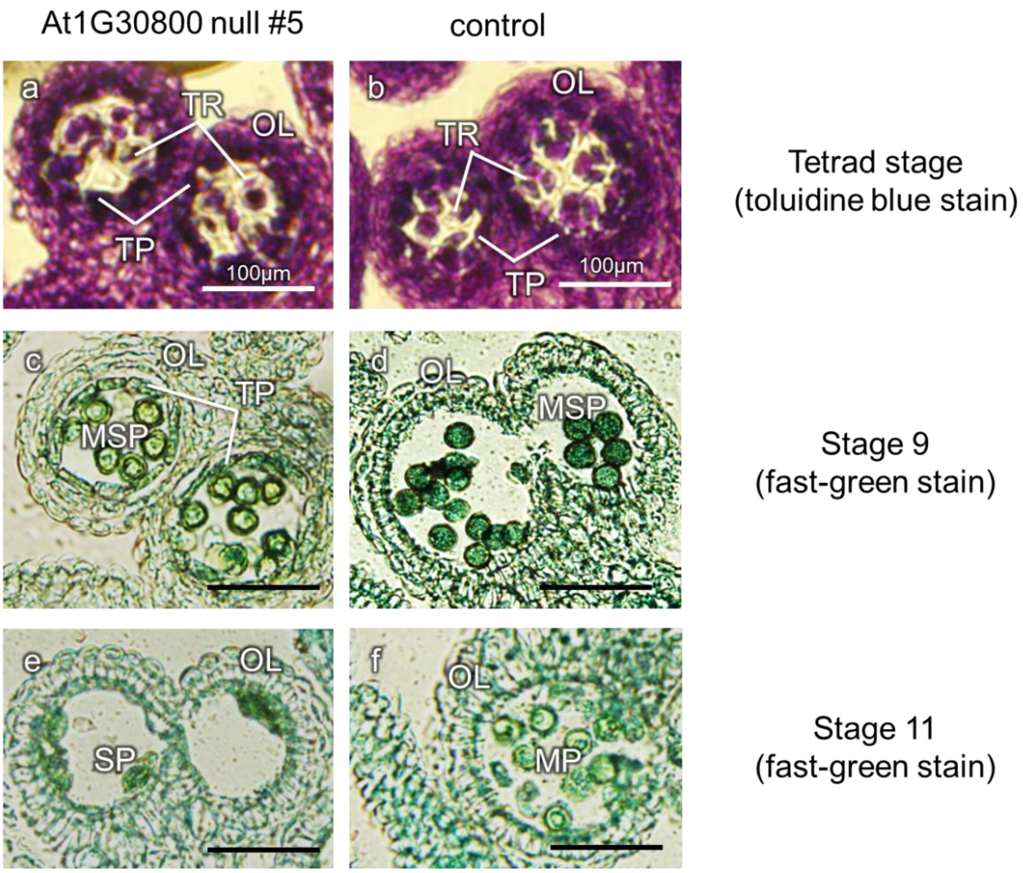


**Supplementary Figure S7 Characterization of anther development in *AT1G30800* null line**

Anthers of *AT1G30800* null mutant line #5 and control lines were dissected at three developmental stages. **a-b**, in the tetrad (TR) stage, both *AT1G30800* null and control lines exhibited proper tapetum layer (TP) formation. OL: outer layer. **c-d**, in stage 9, which corresponds to anther developmental stage 3a-b in kiwifruit (Figure S1), the *AT1G30800* null mutant line maintained thick tapetum layers, while control lines showed complete degradation of tapetum cells. This differentiation between *AT1G30800* null and control is consistent with the difference between male and female anthers in kiwifruit (Figure S1). The microspores (MSP) in control lines were stained in deep green with fast-green, indicating their fertility, while *AT1G30800* null mutant line showed faint staining with fast-green. **e-f**, at the later maturation stages, the tapetum layers were also degenerated in the *AT1G30800* null mutant line, but they only produced shrunk sterile pollen grains (SP). MP: mature pollen.

**Supplemental Figure S8**

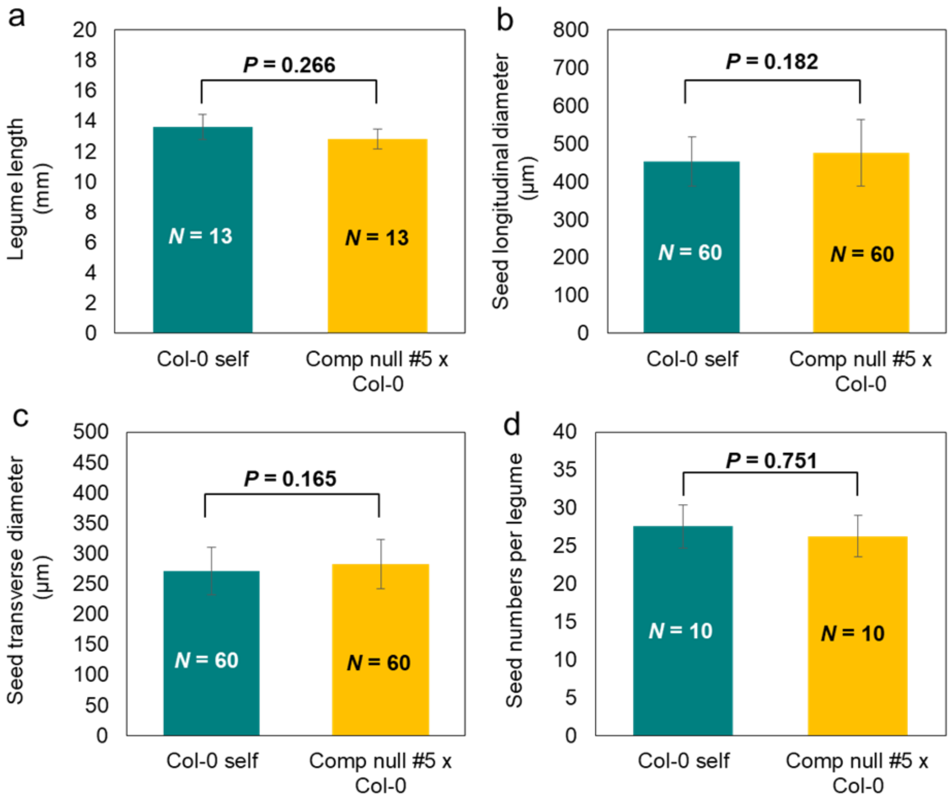


**Supplementary Figure S8 legume and seed production in *AT1G30800* null line**

The female fertility of control (Col-0) and *AT1G30800* mutant null line #5 in cross with control (Col-0) were examined. We assessed silique length (**a**), seed longitudinal diameter (**b**), seed transverse diameter (**c**), and seed numbers per silique (**d**). For silique length and seed numbers per silique, mean value in an individual was used as a biological replicate ($N$ = 13 and 10, respectively). For seed diameter, mean values in a silique was used as a biological replicate ($N$ = 60). All tests showed no significant differences between *AT1G30800* null mutant and control lines ($p > 0.165$), suggesting that *AT1G30800* has no effects on gynoecium functions.

**Supplemental Figure S9**

**a** ***FAS1* (SELMODRAFT_448915-like)**

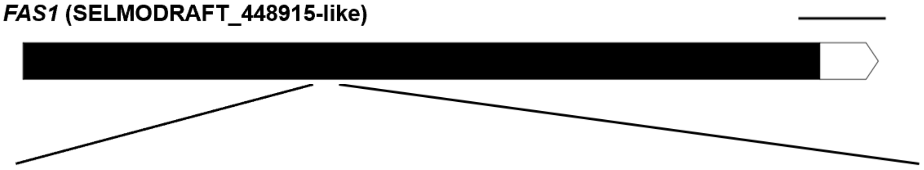

```
              D   A   M   S   N   H   Q   T   P     G   D   Q   N   I   N   L   D
FAS1      5'-GACGCCATGTCAAATCATCAAACAC--CTGGCGATCAAAATATCAACTTAGAC-3'
fas1 #3   5'-GACGCCATGTCAAATCATCAAACATGATAGGCGATCAAAATATCAACTTAGAC-3'
```

**b**

```
FAS1      MIRNGRTIAPNSIPLYLNDTHFDLAILKSLPEKTEYKSPAEALKRSSLKVNNEATEIPTT 60
fas1_#3   MIRNGRTIAPNSIPLYLNDTHFDLAILKSLPEKTEYKSPAEALKRSSLKVNNEATEIPTT 60
          ************************************************************

FAS1      PPPYQQPTTTTFPPPPPPPPLTQHQIQEQQLNNIVEALIGAGDFAGWANLLSSTDLSSLP 120
fas1_#3   PPPYQQPTTTTFPPPPPPPPLTQHQIQEQQLNNIVEALIGAGDFAGWANLLSSTDLSSLP 120
          ************************************************************

FAS1      LSATFFIPGNDAMSNHQTPGDQNINLDPFLIAYHIVPQRLSFADLQQFKSNTRLPTLLPS 180
fas1_#3   LSATFFIPGNDAMSNHQT*----------------------------------------- 171
          ******************

FAS1      KFIVITNNSLTNFTVDGSQITYPDVYVNSAFTVHGVNKVLEYSVYGADPVFSPPENNTVP 240
fas1_#3   ------------------------------------------------------------ 178

FAS1      TETVAPIPPKPKPKPKPKPKPLLFPEGGGGFIDGWRSSNSTRAFPIEKLMLDISVLFFIF 300
fas1_#3   ------------------------------------------------------------ 178

FAS1      FWV* 303
fas1_#3   ---- 178
```

**Supplementary Figure S9 Gene disruption of *FAS1* in *Nicotiana tabacum* using CRISPR/Cas9.**

**a**, Gene model of *FAS1* and alignment of the genomic sequence of *fas1* genome-edited line #3. (Upper) Filled box: CDS, open box: UTRs, scale bar = 100 bp. (Lower) Blue: PAM sequence, underlined: target sequence of the gRNA, magenta: substitution and insertion induced by CRISPR/Cas9. The DNA sequence of *FAS1* was obtained from NCBI (XM_016649347.1). **b**, Sequence alignment of the putative translation product of the edited *fas1* line #3. The putative FAS1 domain (position 59-198, ID:PRU00082 from PRORULE database) is shown in bold.

**Supplemental Figure S10**

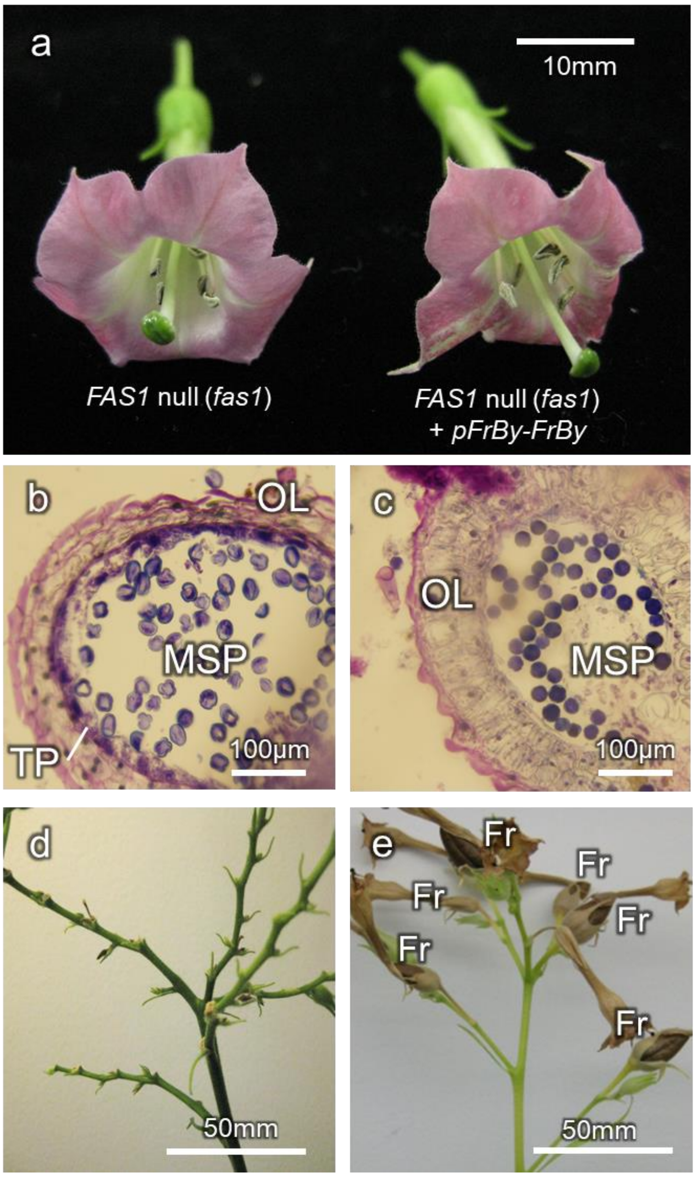


**Supplementary Figure S10 Complementation of male function in *FAS1* null lines by kiwifruit *FrBy*.**

**a**, flower appearance of *FAS1* null (*fas1*) and *fas1* transformed with kiwifruit (*A. chinensis*) *FrBy* under the control of the *FrBy* native promoter (*fas1+pFrBy-FrBy*). **b-c**, dissection of stage 3 anthers in kiwifruit (Figure S1), in *fas1* (**b**) and *fas1+pFrBy-FrBy* (**c**). OL: outer layer, MSP: microspore. *fas1* maintained thick tapetum cells (TP), while *fas1+pFrBy-FrBy* showed complete degradation of tapetum layers. **d-e**, crosses between control (Col-0) as maternal parents and *fas1* (**d**) or *fas1+pFrBy-FrBy* (**e**), as paternal parents. *fas1* pollen grains were sterile, while introduction of kiwifruit *FrBy* restored the male function of *fas1*, resulting in fruit (Fr) production.

#### Supplemental Figure S11

Contigs anchored with Y-specific/Y-linked markers (Akagi et al. 2018)

| | I | II | III | IV | V | VI | VII | VIII | Ø |
|---|---|---|---|---|---|---|---|---|---|
| gem-index seq | Contig-6851 | Contig-8318 | Contig-6898 | Contig-423383 | Contig-426301 | Contig-7598 | Contig-422073 | Contig-5670 | Contig-8003 |
| GATCGGAAGAGCACAC | 0 | 0 | 0 | 0 | 0 | 1 | 1 | 1 | 0 |
| GACGCTCTTCCGATCT | 0 | 0 | 1 | 1 | 1 | 1 | 0 | 0 | 0 |
| GGCGTTTGGACTACAA | 0 | 0 | 0 | 1 | 1 | 1 | 1 | 0 | 0 |
| TCTTGGCGTTCGAACG | 0 | 1 | 1 | 1 | 1 | 0 | 0 | 0 | 0 |
| AAAACCCTAAACCCTA | 0 | 0 | 1 | 1 | 0 | 0 | 0 | 0 | 0 |
| AAACCCTAAACCCTAA | 0 | 0 | 1 | 1 | 0 | 0 | 0 | 0 | 0 |
| AACCCTAAACCCTAAA | 0 | 0 | 1 | 1 | 0 | 0 | 0 | 0 | 0 |
| ACCCTAAACCCTAAAC | 0 | 0 | 1 | 1 | 0 | 0 | 0 | 0 | 0 |
| CCCTAAACCCTAAACC | 0 | 0 | 1 | 1 | 0 | 0 | 0 | 0 | 0 |
| CCTAAACCCTAAACCC | 0 | 0 | 1 | 1 | 0 | 0 | 0 | 0 | 0 |
| CTAAACCCTAAACCCT | 0 | 0 | 1 | 1 | 0 | 0 | 0 | 0 | 0 |
| GGGTTTAGGGTTTAGG | 0 | 0 | 1 | 1 | 0 | 0 | 0 | 0 | 0 |
| GGTTTAGGGTTTAGGG | 0 | 0 | 1 | 1 | 0 | 0 | 0 | 0 | 0 |
| GTTTAGGGTTTAGGGT | 0 | 0 | 1 | 1 | 0 | 0 | 0 | 0 | 0 |
| TAAACCCTAAACCCTA | 0 | 0 | 1 | 1 | 0 | 0 | 0 | 0 | 0 |
| AAATATTCCTAAAAAC | 0 | 0 | 0 | 1 | 1 | 0 | 0 | 0 | 0 |
| ATATATATATATATAT | 0 | 0 | 0 | 0 | 1 | 1 | 0 | 0 | 0 |
| GAACATGCATCCTCAA | 0 | 0 | 1 | 1 | 1 | 0 | 0 | 0 | 0 |
| GAGAATCAAGTGGTAA | 0 | 1 | 0 | 1 | 1 | 0 | 0 | 0 | 0 |
| GATTCGCTCCCAGAAG | 0 | 0 | 0 | 1 | 1 | 0 | 0 | 0 | 0 |
| GCTTACGTTTTCGAAA | 0 | 0 | 0 | 0 | 1 | 1 | 0 | 0 | 0 |
| GGAAGCCACATGGGAT | 0 | 0 | 0 | 0 | 1 | 1 | 0 | 0 | 0 |
| GGAATGATCAGTATAT | 0 | 0 | 0 | 0 | 1 | 1 | 0 | 0 | 0 |
| GGCCATCACACGCCCA | 0 | 0 | 0 | 1 | 1 | 0 | 0 | 0 | 0 |
| GGCGTTCGAACGCCCA | 0 | 0 | 0 | 1 | 1 | 0 | 0 | 0 | 0 |
| GGGAAGCCACATGGGA | 0 | 0 | 0 | 0 | 1 | 1 | 0 | 0 | 0 |
| GGGGAAGCCACATGGG | 0 | 0 | 0 | 0 | 1 | 1 | 0 | 0 | 0 |
| GGGGAATACGACTCTA | 0 | 1 | 1 | 1 | 1 | 0 | 0 | 0 | 0 |
| GGTAAGGAATTCACTT | 0 | 0 | 0 | 0 | 1 | 1 | 0 | 0 | 0 |
| GGTCTTTTCGATAAAT | 0 | 0 | 0 | 0 | 1 | 1 | 0 | 0 | 0 |
| GGTGGTGGTATAGTTC | 0 | 0 | 0 | 0 | 1 | 1 | 0 | 0 | 0 |
| GGTGTTTGGACTACAA | 0 | 0 | 0 | 0 | 1 | 1 | 0 | 0 | 0 |
| GGTTTGTATGATCCCT | 0 | 0 | 0 | 1 | 1 | 0 | 0 | 0 | 0 |
| TGATTCGCTCCCAGAA | 0 | 0 | 0 | 1 | 1 | 0 | 0 | 0 | 0 |
| CAAAACCCTAAACCCT | 0 | 0 | 1 | 1 | 0 | 0 | 0 | 0 | 0 |
| CTAAAACCCTAAACCC | 0 | 0 | 1 | 1 | 0 | 0 | 0 | 0 | 0 |
| CTTGGCGTTCGAACGC | 0 | 0 | 1 | 1 | 0 | 0 | 0 | 0 | 0 |
| GAAAACCCTAAACCCT | 0 | 0 | 1 | 1 | 0 | 0 | 0 | 0 | 0 |
| GCATTTGTATGTCCTC | 0 | 0 | 1 | 1 | 0 | 0 | 0 | 0 | 0 |
| GGGCGTTCGAACGTCC | 0 | 0 | 0 | 1 | 0 | 1 | 0 | 0 | 0 |
| GTAAACCCTAAACCCT | 0 | 0 | 1 | 1 | 0 | 0 | 0 | 0 | 0 |
| GAGGAACCACCCTAAT | 0 | 0 | 0 | 0 | 0 | 1 | 1 | 1 | 0 |
| CCACCACCACCTCCTC | 1 | 1 | 1 | 0 | 0 | 0 | 0 | 0 | 0 |
| GGTGGTGGAGGAGGAG | 1 | 1 | 1 | 0 | 0 | 0 | 0 | 0 | 0 |

**Supplementary Figure S11 Correspondence between GEM barcodes and the anchored scaffolds.**

From the scaffolds assembled with 10X Genomics Supernova v1.2.2, nine scaffolds (I-VIII and Ø) were anchored with Y-specific/Y-linked markers (Akagi et al. 2018) (Table S6). Forty-four GEM index sequences (16-bp), which were mapped to more than 2 of these 9 scaffolds, are listed on the left. From mapped (1, green) and unmapped (0, white) information, using traveling salesman problem (TSP), the most likely physical order of the scaffolds was estimated, and aligned as shown in this figure. Scaffolds III-VII, shown in thick green, are thought to be male-specific sequences enriched regions.

**Supplemental Figure S12**

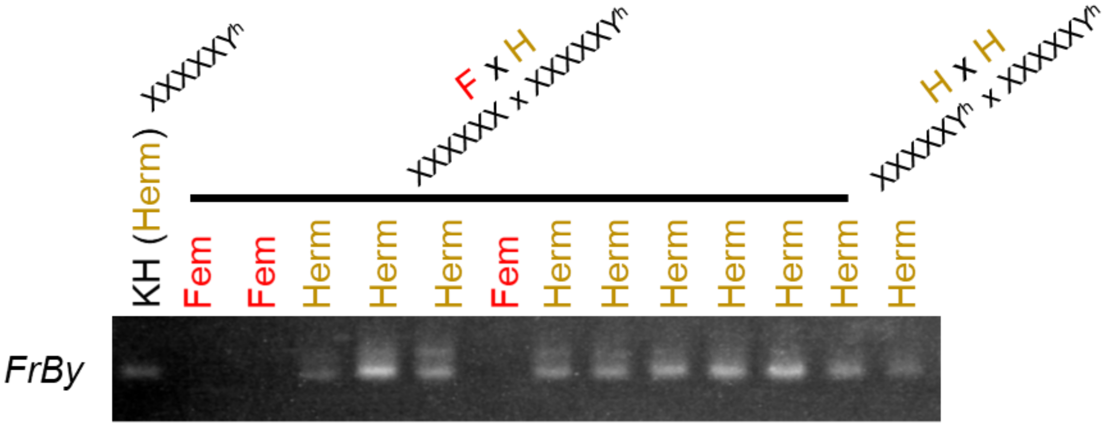


**Supplementary Figure S12 Segregation of *FrBy*, in a KH-derived segregating populations.**

Hermaphroditism (Herm) in the KH-derived population (Table S9) co-segregated with the presence of *FrBy*, as a Y-chromosomal marker. Selfed KH line produced only female (Fem) and hermaphrodite offspring (Table S9). These results suggest that the hermaphrodite phenotype derived from KH line is due to mutation on the Y chromosome (Y to $Y^h$) [or possibly gain of *FrBy* in X chromosome by recombination (X to $X^h$)].

**Supplemental Figure S13**

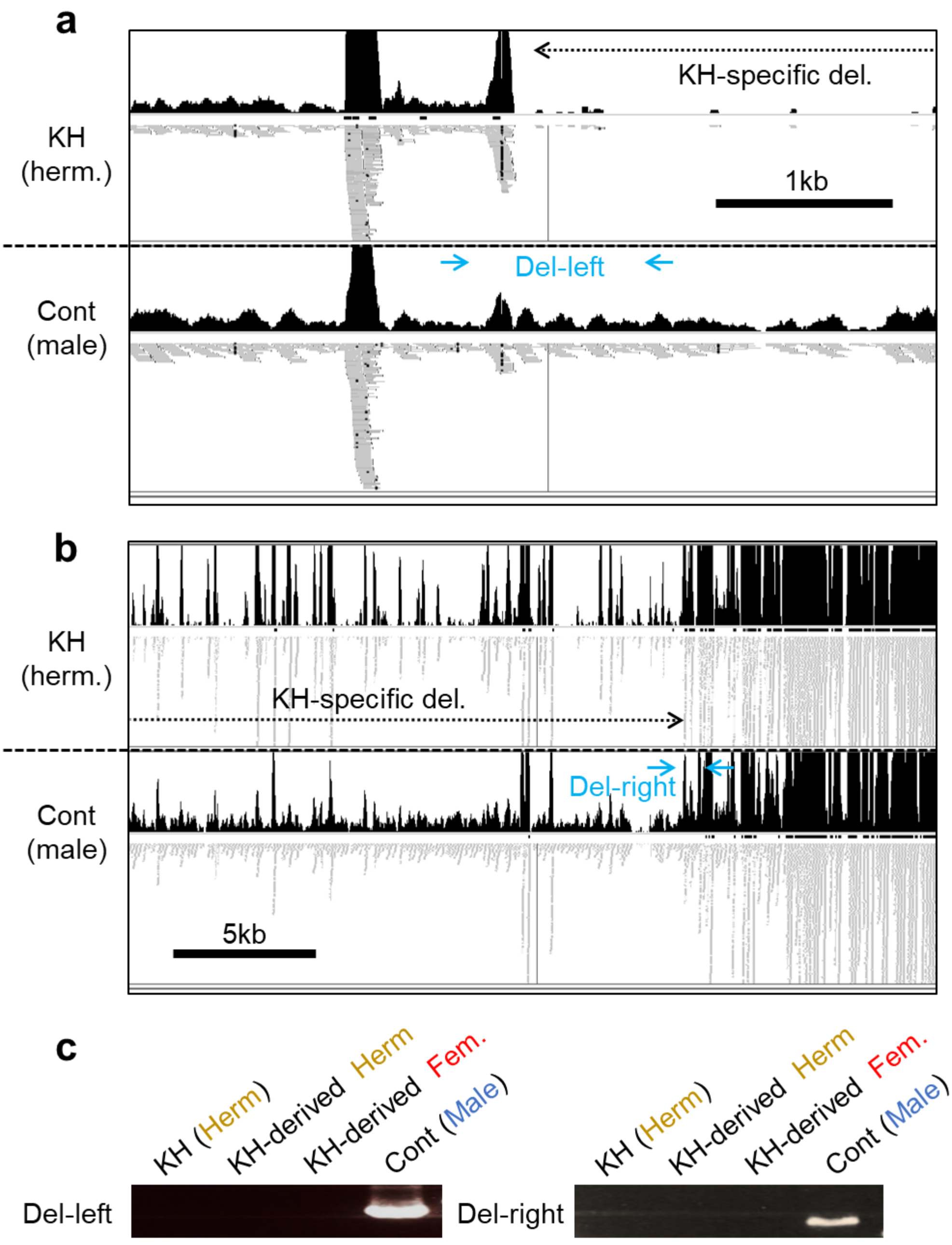


**Supplementary Figure S13 Characterization of the putative KH-specific deletion.**
Mapping of the gDNA Illumina reads from the KH line and control in the contig III (left arm) (**a**) and contig IV (right arm) (**b**). Top and bottom areas in each lane showed reads coverage and mapped reads, respectively. Arrows in sky blue indicated the primers for PCR characterization of the deletion (Del-left and Del-right), corresponding to the panel c. **c,** Both Del-left and Del-right primer sets showed amplicon only in the control male. The KH and its offspring hermaphrodite individuals showed no amplicon, as well as in female individuals.

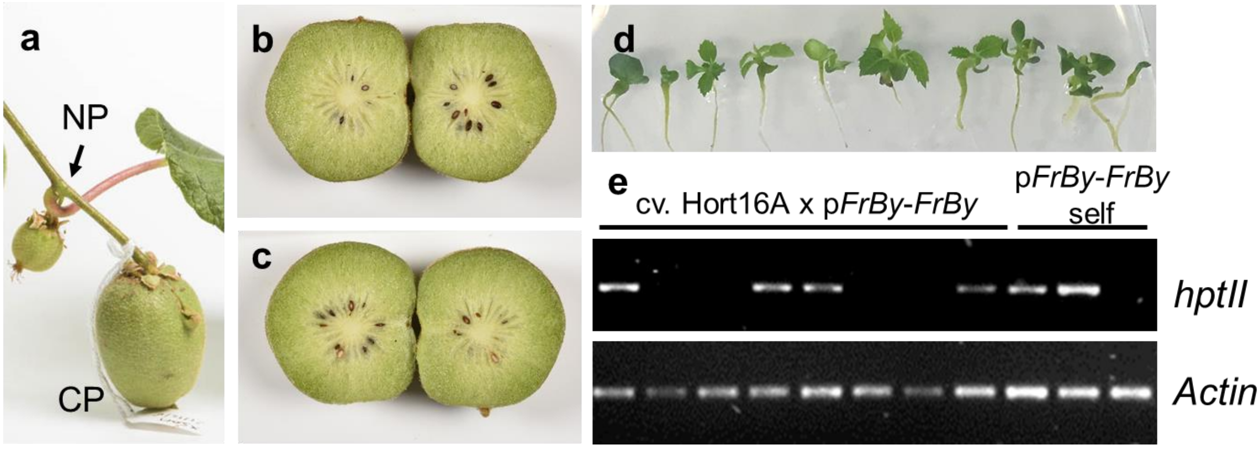


**Supplementary Figure S14 Cross pollination with p*FrBy-FrBy*-induced cv. Hort16A.**

**a,** Pollination of rapid flowering *A. chinensis* female cv. Hort16A with p*FrBy*-*FrBy* (hyg$^{R}$) pollen gives rise to normal fruit (**a**). NP, non-pollinated, CP, cross-pollinated. Both cross pollination of cv. Hort16A x p*FrBy*-*FrBy* (**b**) and self-pollination of p*FrBy*-*FrBy* (**c**) produced fertile seeds, similarly. **d**, Seeds from cross-pollination of cv. Hort16A x p*FrBy*-*FrBy* showed normal germination and seedling development. Approximately ½ of the seedlings are hyg$^{R}$ (**e**).

#### Supplemental Table S1 61 hypothetical genes identified in the MSY contigs and their expression levels in anther

| Contig ID | RPKM ± SE | | | Highest hit in the TAIR10 DB | |
|---|---|---|---|---|---|
| | Male | Female | | gene ID | putative function |
| kiwi151209_m20_contig_9:g1.t1 | 0.00 ± 0.00 | 0.00 ± 0.00 | | AT2G26450.1 | Plant invertase/pectin methylesterase |
| kiwi151209_m20_contig_25:g2.t1 | 0.22 ± 0.05 | 0.00 ± 0.00 | | AT5G24470.1 | APRR5, PRR5 \| pseudo-response regulator |
| kiwi151209_m20_contig_38:g3.t1 | 0.30 ± 0.27 | 0.12 ± 0.11 | | AT3G22980.1 | Ribosomal protein S5/Elongation factor |
| kiwi151209_m20_contig_40:g4.t1 | 0.08 ± 0.07 | 0.00 ± 0.00 | | AT4G03100.1 | Rho GTPase activating protein |
| kiwi151209_m20_contig_59:g5.t1 | 0.44 ± 0.30 | 0.13 ± 0.06 | | AT4G35785.5 | RNA-binding (RRM/RBD/RNP motifs) family |
| kiwi151209_m20_contig_92:g6.t1 | 0.00 ± 0.00 | 0.00 ± 0.00 | | AT2G39480.1 | PGP6 \| P-glycoprotein 6 |
| kiwi151209_m20_contig_98:g7.t1 | 0.33 ± 0.12 | 0.03 ± 0.03 | | AT1G64260.1 | MuDR family transposase |
| kiwi151209_m20_contig_105:g8.t1 | 3.22 ± 0.78 | 4.02 ± 0.29 | | ATCG00490.1 | RBCL \| ribulose-bisphosphate carboxylase |
| kiwi151209_m20_contig_154:g9.t1 | 2.40 ± 0.40 | 3.66 ± 0.34 | | ATCG00480.1 | ATPB, PB \| ATP synthase subunit beta |
| kiwi151209_m20_contig_155:g10.t1 | 0.00 ± 0.00 | 0.06 ± 0.05 | | AT3G29010.1 | Biotin/lipoate A/B protein ligase |
| kiwi151209_m20_contig_182:g11.t1 | 1.49 ± 0.23 | 2.96 ± 0.28 | | ATCG00480.1 | ATPB, PB \| ATP synthase subunit beta |
| kiwi151209_m20_contig_229:g12.t1 | 0.46 ± 0.14 | 0.35 ± 0.08 | | AT4G15720.1 | Tetratricopeptide repeat (TPR)-like |
| kiwi151209_m20_contig_229:g13.t1 | 0.27 ± 0.15 | 0.18 ± 0.11 | | AT1G28327.1 | unknown protein; Has 52 |
| kiwi151209_m20_contig_248:g14.t1 | 0.00 ± 0.00 | 0.00 ± 0.00 | | AT1G68010.1 | HPR, ATHPR1 \| hydroxypyruvate reductase |
| kiwi151209_m20_contig_256:g15.t1 | 0.31 ± 0.12 | 0.05 ± 0.03 | | AT2G04940.1 | scramblase-related |
| kiwi151209_m20_contig_263:g16.t1 | 0.36 ± 0.10 | 0.02 ± 0.02 | | ATMG00860.1 | ORF158 \| DNA/RNA polymerases superfamily |
| kiwi151209_m20_contig_267:g17.t1 | 0.14 ± 0.12 | 0.00 ± 0.00 | | AT4G29420.1 | F-box/RNI-like superfamily protein |
| kiwi151209_m20_contig_322:g18.t1 | 0.00 ± 0.00 | 0.08 ± 0.07 | | AT2G19650.1 | Cysteine/Histidine-rich C1 domain |
| kiwi151209_m20_contig_338:g19.t1 | 1.23 ± 0.09 | 0.62 ± 0.11 | | AT1G42190.1 | GAG/POL/ENV polyprotein |
| kiwi151209_m20_contig_408:g20.t1 | 10.24 ± 1.06 | 6.66 ± 0.62 | | AT4G04740.2 | CPK23 \| calcium-dependent protein kinase |
| kiwi151209_m20_contig_496:g21.t1 | 0.23 ± 0.09 | 0.01 ± 0.01 | | AT1G19835.2 | Plant protein of unknown function |
| kiwi151209_m20_contig_524:g22.t1 | 1.38 ± 0.24 | 0.60 ± 0.06 | | AT2G15180.1 | Zinc knuckle (CCHC-type) family |
| kiwi151209_m20_contig_545:g23.t1 | 0.38 ± 0.14 | 0.02 ± 0.02 | | AT5G51480.1 | SKS2 \| SKU5 similar 2 |
| kiwi151209_m20_contig_545:g24.t1 | 0.19 ± 0.05 | 0.00 ± 0.00 | *Shy Girl* | AT5G26594.1 | ARR24, RR24 \| response regulator 24 |
| kiwi151209_m20_contig_578:g25.t1 | 0.34 ± 0.13 | 0.28 ± 0.08 | | AT3G57300.1 | INO80, ATINO80 \| INO80 ortholog |
| kiwi151209_m20_contig_621:g26.t1 | 1.65 ± 0.47 | 0.00 ± 0.00 | *Friendly Boy* | AT1G30800.1 | Fasciclin-like arabinogalactan family |
| kiwi151209_m20_contig_624:g27.t1 | 0.09 ± 0.06 | 0.01 ± 0.01 | | AT1G09640.2 | Translation elongation factor EF1B |
| kiwi151209_m20_contig_679:g28.t1 | 0.04 ± 0.04 | 0.00 ± 0.00 | | AT2G46230.1 | PIN domain-like family protein |
| kiwi151209_m20_contig_743:g29.t1 | 0.44 ± 0.07 | 0.10 ± 0.03 | | AT5G45130.1 | ATRAB5A, ATRABF2A \| Rab5-related gene |
| kiwi151209_m20_contig_930:g30.t1 | 0.16 ± 0.11 | 0.02 ± 0.01 | | AT5G53790.1 | Protein of unknown function (DUF295) |
| kiwi151209_m20_contig_951:g31.t1 | 0.00 ± 0.00 | 0.00 ± 0.00 | | AT3G04240.1 | SEC \| Tetratricopeptide repeat (TPR)-like |

| | | | | | |
|---|---|---|---|---|---|
| kiwi151209_m20_contig_962:g32.t1 | 0.08 ± 0.07 | 0.04 ± 0.03 | | AT5G21326.1 | Ca2+regulated serine-threonine protein |
| kiwi151209_m20_contig_1037:g33.t1 | 0.44 ± 0.24 | 0.12 ± 0.05 | | AT3G26730.1 | RING/U-box superfamily protein |
| kiwi151209_m20_contig_1060:g34.t1 | 0.12 ± 0.05 | 0.00 ± 0.00 | | AT3G24255.1 | RNA-directed DNA polymerase |
| kiwi151209_m20_contig_1070:g35.t1 | 0.50 ± 0.14 | 0.29 ± 0.16 | | AT1G58390.1 | Disease resistance protein (CC-NBS-LRR) |
| kiwi151209_m20_contig_1092:g36.t1 | 3.82 ± 0.82 | 2.70 ± 0.58 | | AT2G31650.1 | ATX1, SDG27 \| homologue of trithorax |
| kiwi151209_m20_contig_1107:g37.t1 | 0.91 ± 0.16 | 0.45 ± 0.14 | | ATMG00860.1 | ORF158 \| DNA/RNA polymerases superfamily |
| kiwi151209_m20_contig_1148:g38.t1 | 0.42 ± 0.24 | 0.00 ± 0.00 | | AT5G15090.2 | VDAC3 \| voltage dependent anion channel |
| kiwi151209_m20_contig_1182:g39.t1 | 0.21 ± 0.06 | 0.09 ± 0.04 | | no hit | NA |
| kiwi151209_m20_contig_1217:g40.t1 | 0.17 ± 0.13 | 0.05 ± 0.04 | | AT3G20810.3 | JMJD5 \| 2-oxoglutarate (2OG) and Fe(II)-oxygenases |
| kiwi151209_m20_contig_1263:g41.t1 | 0.00 ± 0.00 | 0.00 ± 0.00 | | AT4G11730.1 | Cation transporter/ E1-E2 ATPase family |
| kiwi151209_m20_contig_1434:g42.t1 | 0.18 ± 0.13 | 0.07 ± 0.04 | | no hit | NA |
| kiwi151209_m20_contig_1436:g43.t1 | 0.00 ± 0.00 | 0.00 ± 0.00 | | AT3G28790.1 | Protein of unknown function (DUF1216) |
| kiwi151209_m20_contig_1494:g44.t1 | 0.03 ± 0.02 | 0.16 ± 0.10 | | AT5G61200.1 | unknown protein |
| kiwi151209_m20_contig_1494:g45.t1 | 6.81 ± 1.36 | 9.35 ± 1.17 | | AT1G68990.1 | MGP3 \| male gametophyte defective 3 |
| kiwi151209_m20_contig_1724:g46.t1 | 0.00 ± 0.00 | 0.00 ± 0.00 | *YFT* | AT1G65480.1 | FT \| PEBP (phosphatidylethanolamine-binding) |
| kiwi151209_m20_contig_1854:g47.t1 | 0.24 ± 0.11 | 0.06 ± 0.05 | | AT3G13590.1 | Cysteine/Histidine-rich C1 domain family |
| kiwi151209_m20_contig_1986:g48.t1 | 5.97 ± 0.71 | 11.27 ± 1.46 | | no hit | NA |
| kiwi151209_m20_contig_2004:g49.t1 | 0.00 ± 0.00 | 0.00 ± 0.00 | | AT3G12350.2 | F-box family protein |
| kiwi151209_m20_contig_2011:g50.t1 | 0.28 ± 0.11 | 0.03 ± 0.03 | | AT3G63190.1 | RRF, HFP108 \| ribosome recycling factor |
| kiwi151209_m20_contig_2206:g51.t1 | 0.21 ± 0.18 | 0.04 ± 0.04 | | AT4G26900.1 | AT-HF, HISN4 \| HIS HF |
| kiwi151209_m20_contig_2243:g52.t1 | 0.16 ± 0.07 | 0.00 ± 0.00 | | AT3G55950.1 | CCR3, ATCRR3 \| CRINKLY4 related 3 |
| kiwi151209_m20_contig_2578:g53.t1 | 0.03 ± 0.03 | 0.08 ± 0.05 | | AT3G60980.1 | Tetratricopeptide repeat (TPR)-like |
| kiwi151209_m20_contig_2616:g54.t1 | 0.76 ± 0.30 | 0.95 ± 0.28 | | AT5G51630.2 | Disease resistance protein (TIR-NBS-LRR) |
| kiwi151209_m20_contig_2826:g55.t1 | 1.58 ± 0.35 | 1.07 ± 0.16 | | AT3G57280.1 | Transmembrane proteins 14C |
| kiwi151209_m20_contig_2924:g56.t1 | 0.08 ± 0.07 | 0.00 ± 0.00 | | AT3G12250.4 | TGA6, BZIP45 \| TGACG motif-binding factor |
| kiwi151209_m20_contig_3379:g57.t1 | 0.00 ± 0.00 | 0.00 ± 0.00 | | AT4G19930.1 | F-box and associated interaction domain |
| kiwi151209_m20_contig_3626:g58.t1 | 0.00 ± 0.00 | 0.00 ± 0.00 | | AT5G17520.1 | RCP1, MEX1 \| root cap 1 (RCP1) |
| kiwi151209_m20_contig_3804:g59.t1 | 0.00 ± 0.00 | 0.00 ± 0.00 | | AT1G70040.1 | Protein of unknown function (DUF1163) |
| kiwi151209_m20_contig_3872:g60.t1 | 0.00 ± 0.00 | 0.00 ± 0.00 | | AT2G30500.2 | Kinase interacting (KIP1-like) family |
| kiwi151209_m20_contig_3940:g61.t1 | 0.03 ± 0.03 | 0.00 ± 0.00 | | AT4G03020.2 | transducin family protein / WD-40 repeat |

#### Supplemental Table S2 List of DEGs between male and female anthers

| Gene ID | expressio level (RPKM) | | FDR value | $log_2$ (Male/Female) | Gene annotation by TAIR | |
|---|---|---|---|---|---|---|
| | Female ave | Male ave | | | Gene ID | Gene abbreviation |
| Acc22078.1 | 0.79 | 6.06 | 2.0E-06 | 3.88 | AT3G27060.1 | TSO2, ATTSO2 |
| Acc02105.1 | 0.37 | 3.10 | 5.1E-06 | 3.90 | AT4G14430.1 | IBR10, ATECI2, ECI2, ECHIB, PEC12 |
| Acc29816.1 | 0.64 | 4.18 | 8.1E-06 | 3.71 | AT3G48160.2 | DEL1, E2L3, E2FE |
| Acc23350.1 | 21.06 | 1.76 | 1.4E-04 | -2.69 | AT5G12010.1 | |
| Acc01225.1 | 1.13 | 5.24 | 1.4E-04 | 3.16 | AT5G65090.1 | MRH3, BST1, DER4 |
| Acc17794.1 | 2.14 | 67.12 | 1.4E-04 | 5.99 | No | |
| Acc13477.1 | 0.66 | 3.61 | 2.6E-04 | 3.38 | No | |
| Acc14927.1 | 1.66 | 7.01 | 2.6E-04 | 3.01 | AT2G05160.1 | |
| Acc24741.1 | 4.03 | 10.88 | 4.2E-04 | 2.35 | No | |
| Acc24019.1 | 4.07 | 12.75 | 5.4E-04 | 2.59 | AT1G76740.1 | |
| Acc28464.1 | 6.11 | 15.21 | 5.4E-04 | 2.25 | No | |
| Acc13986.1 | 1.22 | 4.22 | 1.4E-03 | 2.74 | AT5G53670.1 | |
| Acc18343.1 | 1.82 | 7.12 | 1.4E-03 | 2.86 | AT5G08020.1 | ATRPA70B, RPA70B |
| Acc21103.1 | 0.32 | 1.65 | 1.4E-03 | 3.25 | AT4G34131.1 | UGT73B3 |
| Acc27496.1 | 1.74 | 6.01 | 1.4E-03 | 2.72 | AT5G43530.1 | |
| Acc28184.1 | 0.09 | 1.02 | 1.5E-03 | 4.44 | AT5G45500.2 | |
| Acc31257.1 | 0.42 | 1.63 | 1.6E-03 | 2.87 | AT1G68740.1 | PHO1;H1 |
| Acc26447.1 | 1.61 | 3.89 | 1.6E-03 | 2.15 | AT1G53270.1 | |
| Acc13203.1 | 0.98 | 2.75 | 1.6E-03 | 2.44 | AT3G50120.1 | |
| Acc14090.1 | 0.30 | 1.53 | 1.6E-03 | 3.27 | AT2G03370.1 | |
| Acc15608.1 | 1.30 | 3.96 | 1.6E-03 | 2.57 | AT5G41790.1 | CIP1 |
| Acc19788.1 | 12.77 | 0.92 | 1.6E-03 | -2.97 | AT3G61680.1 | |
| Acc31647.1 | 1.35 | 3.52 | 1.6E-03 | 2.29 | AT3G47570.1 | |
| Acc32283.1 | 1.59 | 4.65 | 2.3E-03 | 2.48 | AT1G05700.1 | |
| Acc10059.1 | 0.76 | 2.92 | 2.4E-03 | 2.87 | AT1G58190.2 | RLP9 |
| Acc29667.1 | 0.21 | 1.11 | 2.7E-03 | 3.38 | AT5G52450.1 | |
| Acc31650.1 | 6.56 | 15.38 | 2.8E-03 | 2.10 | AT2G47990.1 | SWA1, EDA13, EDA19 |
| Acc18575.1 | 10.06 | 21.05 | 3.1E-03 | 1.96 | ATMG00860.1 | ORF158 |
| Acc28342.1 | 2.02 | 5.00 | 3.2E-03 | 2.19 | AT4G04650.1 | |
| Acc19957.1 | 0.46 | 1.63 | 3.3E-03 | 2.75 | AT4G22900.1 | |
| Acc24910.1 | 3.00 | 6.72 | 3.3E-03 | 2.01 | AT5G35450.1 | |
| Acc11663.1 | 22.39 | 2.36 | 3.3E-03 | -2.32 | AT5G64260.1 | EXL2 |
| Acc00694.1 | 0.25 | 1.18 | 3.4E-03 | 3.21 | AT1G07390.2 | AtRLP1, RLP1 |
| Acc29498.1 | 1.57 | 4.88 | 3.5E-03 | 2.43 | AT5G45275.1 | |
| Acc02256.1 | 0.41 | 1.63 | 3.9E-03 | 2.95 | AT5G15900.1 | TBL19 |
| Acc33091.1 | 3.53 | 8.17 | 3.9E-03 | 2.13 | AT5G49110.1 | |
| Acc00125.1 | 1.55 | 4.40 | 4.2E-03 | 2.48 | AT4G31980.1 | |
| Acc00600.1 | 0.49 | 1.66 | 4.2E-03 | 2.73 | AT2G26710.1 | BAS1, CYP734A1, CYP72B1 |
| Acc02653.1 | 0.33 | 1.26 | 4.2E-03 | 2.83 | AT1G64080.1 | |
| Acc03541.1 | 6.98 | 14.71 | 4.2E-03 | 1.94 | AT4G23160.1 | CRK8 |
| Acc07843.1 | 1.53 | 4.43 | 4.2E-03 | 2.49 | AT3G05130.1 | |
| Acc08805.1 | 0.28 | 1.02 | 4.2E-03 | 2.73 | AT3G48880.2 | |
| Acc15922.1 | 11.00 | 22.46 | 4.2E-03 | 1.93 | No | |
| Acc18609.1 | 5.78 | 12.16 | 4.2E-03 | 1.94 | AT1G43760.1 | |
| Acc24016.1 | 1.85 | 4.36 | 4.2E-03 | 2.11 | AT4G08850.1 | |
| Acc33026.1 | 1.54 | 4.90 | 4.2E-03 | 2.62 | AT1G14430.1 | |
| Acc24562.1 | 247.58 | 17.29 | 4.4E-03 | -3.13 | AT5G49360.1 | BXL1, ATBXL1 |
| Acc20479.1 | 0.41 | 1.78 | 4.6E-03 | 3.00 | AT2G36770.1 | |
| Acc24742.1 | 11.39 | 23.45 | 4.8E-03 | 1.94 | AT1G48120.1 | |
| Acc04665.1 | 1.42 | 3.82 | 4.9E-03 | 2.32 | AT4G36870.2 | BLH2, SAW1 |
| Acc18659.1 | 0.19 | 1.19 | 4.9E-03 | 3.63 | AT1G54540.1 | |
| Acc16057.1 | 1.91 | 5.84 | 4.9E-03 | 2.58 | AT1G62310.1 | |
| Acc26925.1 | 1.83 | 4.78 | 5.0E-03 | 2.32 | AT1G52190.1 | |
| Acc06511.1 | 1.34 | 4.42 | 5.5E-03 | 2.68 | AT1G78580.1 | ATTPS1, TPS1 |
| Acc06438.1 | 7.38 | 14.37 | 6.1E-03 | 1.86 | AT3G42170.1 | |
| Acc10104.1 | 0.29 | 1.15 | 6.1E-03 | 2.72 | AT2G18690.1 | |
| Acc01442.1 | 37.48 | 4.62 | 6.3E-03 | -2.23 | AT5G51910.2 | |
| Acc20835.1 | 0.77 | 2.14 | 6.3E-03 | 2.36 | AT3G61170.1 | |
| Acc18495.1 | 19.38 | 37.96 | 6.4E-03 | 1.88 | AT4G23160.1 | CRK8 |
| Acc00053.1 | 1.90 | 4.68 | 6.8E-03 | 2.19 | AT3G51740.1 | IMK2 |
| Acc02315.1 | 0.70 | 2.07 | 7.0E-03 | 2.43 | AT3G16520.3 | UGT88A1 |
| Acc13682.1 | 0.48 | 2.05 | 7.3E-03 | 2.95 | AT4G21700.1 | |
| Acc01263.1 | 0.62 | 2.04 | 7.9E-03 | 2.57 | AT3G19380.1 | PUB25 |
| Acc19156.1 | 0.28 | 1.10 | 7.9E-03 | 2.91 | AT5G41040.1 | |
| Acc02399.1 | 0.53 | 1.74 | 8.0E-03 | 2.65 | AT2G26730.1 | |
| Acc09308.1 | 0.87 | 2.56 | 8.0E-03 | 2.47 | AT3G08500.1 | MYB83, AtMYB83 |

| | | | | | | |
|---|---|---|---|---|---|---|
| Acc12879.1 | 0.40 | 1.47 | 8.0E-03 | 2.90 | AT2G29110.1 | ATGLR2.8, GLR2.8 |
| Acc13840.1 | 1.09 | 3.15 | 8.0E-03 | 2.46 | AT2G19130.1 | |
| Acc25258.1 | 0.88 | 2.37 | 8.0E-03 | 2.28 | AT5G44440.1 | |
| Acc32063.1 | 0.51 | 1.63 | 8.1E-03 | 2.50 | AT3G11840.1 | PUB24 |
| Acc06831.1 | 5.69 | 11.74 | 8.6E-03 | 1.91 | AT4G38070.1 | |
| Acc13779.1 | 4.82 | 8.94 | 8.6E-03 | 1.78 | No | |
| Acc02232.1 | 20.97 | 43.32 | 8.7E-03 | 1.97 | AT4G23160.1 | CRK8 |
| Acc19497.1 | 5.38 | 10.51 | 8.7E-03 | 1.84 | AT4G10780.1 | |
| Acc30443.1 | 4.31 | 10.70 | 8.7E-03 | 2.24 | AT1G69770.1 | CMT3 |
| Acc26295.1 | 3.23 | 6.32 | 8.9E-03 | 1.89 | AT3G26330.1 | CYP71B37 |
| Acc28127.1 | 0.52 | 1.67 | 8.9E-03 | 2.61 | AT4G18210.1 | ATPUP10, PUP10 |
| Acc01693.1 | 0.47 | 1.67 | 9.4E-03 | 2.78 | AT2G37900.1 | |
| Acc05285.1 | 3.27 | 7.34 | 9.4E-03 | 2.09 | AT4G23160.1 | CRK8 |
| Acc06585.1 | 0.68 | 2.56 | 9.4E-03 | 2.78 | AT4G02330.1 | ATPMEPCRB |
| Acc18614.1 | 19.88 | 35.59 | 9.5E-03 | 1.74 | No | |
| Acc05425.1 | 0.38 | 1.51 | 9.6E-03 | 2.82 | AT1G17840.1 | WBC11, ABCG11, DSO, COF1, ATWBC11 |
| Acc18692.1 | 7.68 | 14.18 | 9.6E-03 | 1.80 | No | |
| Acc11901.1 | 1.02 | 3.46 | 9.9E-03 | 2.70 | AT1G20060.1 | |
| Acc30420.1 | 2.10 | 4.21 | 9.9E-03 | 1.93 | AT5G12890.1 | |
| Acc09188.1 | 0.30 | 1.40 | 1.0E-02 | 3.08 | AT4G01140.1 | |
| Acc29423.1 | 2.15 | 4.83 | 1.0E-02 | 2.03 | AT3G47570.1 | |
| Acc06843.1 | 1.70 | 3.78 | 1.0E-02 | 2.00 | AT5G10530.1 | |
| Acc12153.1 | 0.66 | 1.98 | 1.1E-02 | 2.44 | AT2G22490.2 | CYCD2;1 |
| Acc15925.1 | 6.27 | 13.00 | 1.1E-02 | 1.99 | No | |
| Acc00881.1 | 1.10 | 2.70 | 1.1E-02 | 2.21 | AT4G10310.1 | HKT1, ATHKT1 |
| Acc12988.1 | 37.99 | 3.82 | 1.1E-02 | -2.58 | AT5G12010.1 | |
| Acc19629.1 | 0.64 | 1.79 | 1.1E-02 | 2.44 | AT5G07900.1 | |
| Acc25261.1 | 0.26 | 1.16 | 1.1E-02 | 3.00 | AT5G44440.1 | |
| Acc28254.1 | 20.15 | 41.20 | 1.1E-02 | 1.97 | No | |
| Acc31722.1 | 0.53 | 2.10 | 1.1E-02 | 2.80 | AT1G03070.2 | |
| Acc32516.1 | 0.43 | 1.79 | 1.1E-02 | 2.96 | AT1G55230.1 | |
| Acc09149.1 | 1.34 | 2.99 | 1.1E-02 | 2.09 | AT2G46060.1 | |
| Acc17167.1 | 0.72 | 1.80 | 1.1E-02 | 2.22 | AT1G32450.1 | NRT1.5 |
| Acc32255.1 | 0.50 | 1.42 | 1.1E-02 | 2.46 | AT3G50160.1 | |
| Acc13740.1 | 0.50 | 1.52 | 1.1E-02 | 2.50 | AT5G23960.2 | TPS21 |
| Acc00029.1 | 1.71 | 3.89 | 1.2E-02 | 2.12 | AT1G66910.1 | |
| Acc30381.1 | 0.33 | 1.38 | 1.2E-02 | 2.90 | AT1G51405.1 | |
| Acc25188.1 | 1.78 | 4.40 | 1.2E-02 | 2.17 | AT5G19410.1 | |
| Acc13971.1 | 5.65 | 10.59 | 1.2E-02 | 1.74 | AT1G06520.1 | ATGPAT1, GPAT1 |
| Acc06059.1 | 0.38 | 1.33 | 1.3E-02 | 2.64 | AT2G23620.1 | ATMES1, MES1 |
| Acc09534.1 | 7.80 | 0.89 | 1.3E-02 | -2.35 | No | |
| Acc32510.1 | 4.64 | 10.61 | 1.3E-02 | 2.16 | AT1G69770.1 | CMT3 |
| Acc21309.1 | 0.29 | 1.39 | 1.3E-02 | 3.28 | AT1G30040.1 | ATGA2OX2, GA2OX2 |
| Acc21983.1 | 0.35 | 1.07 | 1.3E-02 | 2.49 | AT5G50120.1 | |
| Acc14254.1 | 0.50 | 1.57 | 1.4E-02 | 2.58 | AT1G24430.1 | |
| Acc16643.1 | 0.64 | 1.61 | 1.4E-02 | 2.23 | AT4G27190.1 | |
| Acc31877.1 | 0.60 | 1.91 | 1.4E-02 | 2.59 | AT4G39950.1 | CYP79B2 |
| Acc33420.1 | 2.41 | 4.57 | 1.4E-02 | 1.81 | AT4G38180.1 | FRS5 |
| Acc20272.1 | 1.96 | 4.92 | 1.4E-02 | 2.28 | AT5G52950.1 | |
| Acc09737.1 | 6.43 | 11.29 | 1.4E-02 | 1.69 | AT4G04480.1 | |
| Acc10467.1 | 0.37 | 1.30 | 1.4E-02 | 2.73 | AT5G08520.1 | |
| Acc25212.1 | 2.14 | 4.50 | 1.4E-02 | 1.98 | AT5G46220.1 | |
| Acc32722.1 | 0.46 | 1.37 | 1.5E-02 | 2.55 | AT5G14090.1 | |
| Acc05523.1 | 0.95 | 2.65 | 1.6E-02 | 2.46 | No | |
| Acc21202.1 | 0.70 | 2.23 | 1.6E-02 | 2.58 | AT1G22150.1 | SULTR1;3 |
| Acc03044.1 | 0.59 | 1.63 | 1.6E-02 | 2.43 | AT2G38510.1 | |
| Acc02403.1 | 0.26 | 1.09 | 1.6E-02 | 3.01 | No | |
| Acc19680.1 | 1.72 | 4.47 | 1.6E-02 | 2.23 | AT2G47430.1 | CKI1 |
| Acc22934.1 | 10.88 | 18.70 | 1.6E-02 | 1.69 | No | |
| Acc28557.1 | 1.07 | 2.54 | 1.6E-02 | 2.14 | AT1G13230.1 | |
| Acc30705.1 | 1.49 | 3.16 | 1.6E-02 | 1.99 | AT4G08850.2 | |
| Acc31874.1 | 0.31 | 1.04 | 1.6E-02 | 2.64 | AT3G26310.1 | CYP71B35 |
| Acc22629.1 | 0.36 | 1.20 | 1.6E-02 | 2.69 | AT5G17540.1 | |
| Acc18607.1 | 21.21 | 38.84 | 1.7E-02 | 1.80 | AT4G23160.1 | CRK8 |
| Acc28223.1 | 0.92 | 2.33 | 1.7E-02 | 2.19 | AT5G51160.1 | |
| Acc01300.1 | 4.07 | 6.85 | 1.8E-02 | 1.63 | AT5G54160.1 | ATOMT1, OMT1 |
| Acc21739.1 | 0.82 | 1.93 | 1.8E-02 | 2.15 | AT1G53270.1 | |
| Acc26823.1 | 1.55 | 6.80 | 1.8E-02 | 3.12 | AT3G27060.1 | TSO2, ATTSO2 |
| Acc28802.1 | 2.45 | 4.61 | 1.8E-02 | 1.79 | AT3G59140.1 | ATMRP14, MRP14, ABCC10 |
| Acc30924.1 | 0.31 | 1.17 | 1.8E-02 | 2.81 | AT3G09290.1 | TAC1 |
| Acc32284.1 | 3.80 | 7.15 | 1.8E-02 | 1.78 | AT1G07390.3 | AtRLP1, RLP1 |
| Acc33533.1 | 6.35 | 12.24 | 1.8E-02 | 1.87 | No | |

| | | | | | | |
|---|---|---|---|---|---|---|
| Acc09655.1 | 1.81 | 3.81 | 1.8E-02 | 2.02 | AT1G11330.1 | |
| Acc16157.1 | 0.71 | 1.64 | 1.9E-02 | 2.08 | No | |
| Acc06356.1 | 0.65 | 1.80 | 1.9E-02 | 2.29 | AT1G14040.1 | |
| Acc08635.1 | 5.48 | 9.37 | 1.9E-02 | 1.66 | AT3G47570.1 | |
| Acc16802.1 | 1.34 | 3.18 | 1.9E-02 | 2.15 | AT5G58630.1 | |
| Acc17599.1 | 0.89 | 2.73 | 1.9E-02 | 2.50 | AT2G29110.1 | ATGLR2.8, GLR2.8 |
| Acc19493.1 | 1.18 | 2.76 | 1.9E-02 | 2.20 | AT4G10780.1 | |
| Acc15833.1 | 0.31 | 1.10 | 2.0E-02 | 2.69 | AT3G57430.1 | OTP84 |
| Acc18550.1 | 3.16 | 5.82 | 2.0E-02 | 1.82 | AT1G07390.3 | AtRLP1, RLP1 |
| Acc24726.1 | 2.92 | 5.50 | 2.0E-02 | 1.79 | No | |
| Acc32890.1 | 5.27 | 8.94 | 2.0E-02 | 1.68 | AT1G43760.1 | |
| Acc00507.1 | 2.97 | 6.20 | 2.0E-02 | 1.95 | No | |
| Acc08006.1 | 1.55 | 3.22 | 2.0E-02 | 1.92 | AT3G52430.1 | PAD4, ATPAD4 |
| Acc09652.1 | 1.21 | 2.88 | 2.0E-02 | 2.16 | AT1G61610.1 | |
| Acc18523.1 | 0.46 | 1.25 | 2.0E-02 | 2.39 | AT5G46330.1 | FLS2 |
| Acc00950.1 | 0.24 | 1.38 | 2.0E-02 | 3.40 | AT2G26490.1 | |
| Acc05289.1 | 1.65 | 3.39 | 2.0E-02 | 1.96 | AT2G17525.1 | |
| Acc19054.1 | 0.58 | 1.65 | 2.0E-02 | 2.36 | AT3G49950.1 | |
| Acc19150.1 | 0.91 | 2.44 | 2.0E-02 | 2.32 | AT4G32300.1 | SD2-5 |
| Acc33502.1 | 1.29 | 3.21 | 2.0E-02 | 2.30 | AT2G01050.1 | |
| Acc23755.1 | 1.26 | 3.10 | 2.0E-02 | 2.15 | AT5G67090.1 | |
| Acc33278.1 | 3.58 | 8.72 | 2.0E-02 | 2.21 | AT2G05160.1 | |
| Acc23743.2 | 4.32 | 7.05 | 2.1E-02 | 1.59 | AT2G29120.1 | ATGLR2.7, GLR2.7 |
| Acc04973.1 | 2.20 | 4.23 | 2.1E-02 | 1.85 | AT1G35710.1 | |
| Acc06038.1 | 0.46 | 1.36 | 2.1E-02 | 2.44 | AT4G36990.1 | HSF4, HSFB1, AT-HSFB1, ATHSF4 |
| Acc31789.1 | 0.47 | 1.35 | 2.1E-02 | 2.45 | AT5G04110.1 | GYRB3 |
| Acc24077.1 | 1.27 | 2.78 | 2.1E-02 | 2.03 | AT1G21270.1 | WAK2 |
| Acc31517.1 | 24.50 | 45.95 | 2.1E-02 | 1.83 | No | |
| Acc21101.1 | 0.47 | 1.14 | 2.1E-02 | 2.19 | AT4G34131.1 | UGT73B3 |
| Acc23674.1 | 1.53 | 3.41 | 2.1E-02 | 2.07 | AT1G75100.1 | JAC1 |
| Acc13107.1 | 1.13 | 3.10 | 2.1E-02 | 2.44 | AT4G13370.1 | |
| Acc28344.1 | 27.89 | 49.80 | 2.2E-02 | 1.74 | No | |
| Acc18899.1 | 0.39 | 1.30 | 2.2E-02 | 2.60 | AT1G29340.1 | PUB17, ATPUB17 |
| Acc10060.1 | 1.53 | 3.02 | 2.3E-02 | 1.83 | AT1G74190.1 | AtRLP15, RLP15 |
| Acc16189.1 | 0.47 | 1.36 | 2.3E-02 | 2.41 | AT5G60900.1 | RLK1 |
| Acc12456.1 | 1.19 | 2.58 | 2.3E-02 | 2.04 | AT3G15730.1 | PLDALPHA1, PLD |
| Acc01351.1 | 0.84 | 2.15 | 2.3E-02 | 2.33 | AT1G66910.1 | |
| Acc32436.1 | 0.52 | 1.61 | 2.4E-02 | 2.60 | AT3G10330.1 | |
| Acc21010.1 | 2.23 | 4.31 | 2.4E-02 | 1.83 | AT5G48060.1 | |
| Acc23858.1 | 1.81 | 4.15 | 2.4E-02 | 2.05 | AT5G49660.1 | |
| Acc29581.1 | 26.82 | 4.14 | 2.4E-02 | -1.85 | AT1G67940.1 | ATNAP3, AtSTAR1, NAP3 |
| Acc21323.1 | 0.92 | 2.28 | 2.4E-02 | 2.20 | AT2G13620.1 | ATCHX15, CHX15 |
| Acc12345.1 | 3.60 | 6.53 | 2.4E-02 | 1.73 | No | |
| Acc16953.1 | 1.13 | 2.35 | 2.4E-02 | 1.94 | AT3G09640.2 | APX2, APX1B |
| Acc21398.1 | 1.32 | 2.62 | 2.4E-02 | 1.83 | AT2G27430.1 | |
| Acc27443.1 | 1.61 | 4.25 | 2.5E-02 | 2.33 | AT5G03250.1 | |
| Acc19343.1 | 2.36 | 5.64 | 2.5E-02 | 2.24 | AT2G05160.1 | |
| Acc19746.1 | 25.53 | 46.15 | 2.5E-02 | 1.76 | No | |
| Acc28412.1 | 22.28 | 38.43 | 2.5E-02 | 1.68 | AT5G13710.2 | SMT1, CPH |
| Acc32636.1 | 0.80 | 2.06 | 2.5E-02 | 2.24 | AT3G25830.1 | ATTPS-CIN, TPS-CIN |
| Acc22571.1 | 0.40 | 1.35 | 2.5E-02 | 2.68 | AT5G25840.1 | |
| Acc20822.1 | 2.13 | 5.01 | 2.6E-02 | 2.14 | AT3G03300.3 | DCL2 |
| Acc21521.1 | 7.01 | 11.25 | 2.6E-02 | 1.60 | AT1G19260.1 | |
| Acc04137.1 | 1.98 | 3.82 | 2.7E-02 | 1.85 | AT1G70170.1 | MMP |
| Acc32413.1 | 1.95 | 3.41 | 2.8E-02 | 1.68 | AT5G27110.1 | |
| Acc04786.1 | 0.47 | 1.43 | 2.8E-02 | 2.40 | AT5G43190.1 | |
| Acc06827.1 | 4.20 | 7.15 | 2.8E-02 | 1.69 | AT5G36930.1 | |
| Acc09812.1 | 0.73 | 1.72 | 2.8E-02 | 2.15 | AT1G72610.1 | GLP1, ATGER1, GER1 |
| Acc04642.1 | 2.32 | 4.03 | 2.8E-02 | 1.74 | AT5G42710.1 | |
| Acc13761.1 | 0.67 | 1.46 | 2.8E-02 | 2.01 | AT2G03200.1 | |
| Acc05988.1 | 0.71 | 1.82 | 2.8E-02 | 2.19 | AT5G67385.1 | |
| Acc15801.1 | 0.32 | 1.01 | 2.9E-02 | 2.62 | AT5G47580.1 | |
| Acc06352.1 | 1.24 | 3.07 | 2.9E-02 | 2.17 | AT1G26760.1 | ATXR1, SDG35 |
| Acc08145.1 | 1.37 | 2.90 | 2.9E-02 | 1.94 | AT4G39490.1 | CYP96A10 |
| Acc09739.1 | 0.95 | 2.02 | 2.9E-02 | 1.97 | AT4G04480.1 | |
| Acc13698.1 | 0.49 | 1.36 | 2.9E-02 | 2.29 | AT2G14830.1 | |
| Acc16243.1 | 1.29 | 2.72 | 2.9E-02 | 2.03 | AT3G23330.1 | |
| Acc24522.1 | 0.35 | 1.12 | 2.9E-02 | 2.58 | AT1G02400.1 | ATGA2OX4, ATGA2OX6, DTA1, GA2OX6 |
| Acc32146.1 | 0.87 | 1.96 | 2.9E-02 | 1.96 | AT2G34930.1 | |
| Acc33541.1 | 0.78 | 2.53 | 2.9E-02 | 2.55 | No | |
| Acc13356.1 | 0.61 | 1.84 | 2.9E-02 | 2.38 | AT3G11840.1 | PUB24 |
| Acc03042.1 | 0.46 | 1.43 | 2.9E-02 | 2.58 | AT3G51630.2 | WNK5 |

| | | | | | | |
|---|---|---|---|---|---|---|
| Acc09940.1 | 0.88 | 2.06 | 2.9E-02 | 2.07 | AT2G26710.1 | BAS1, CYP734A1, CYP72B1 |
| Acc20688.1 | 1.13 | 2.21 | 2.9E-02 | 1.88 | AT1G53230.1 | TCP3 |
| Acc18867.1 | 1.01 | 2.33 | 3.0E-02 | 2.06 | AT4G21700.1 | |
| Acc01557.1 | 2.11 | 4.15 | 3.0E-02 | 1.87 | AT1G59740.1 | |
| Acc02163.1 | 0.72 | 1.84 | 3.0E-02 | 2.27 | AT3G55550.1 | |
| Acc11554.1 | 1.93 | 3.39 | 3.0E-02 | 1.70 | AT1G43980.1 | |
| Acc19411.1 | 0.80 | 2.13 | 3.0E-02 | 2.22 | AT4G27190.1 | |
| Acc29159.1 | 0.52 | 1.53 | 3.0E-02 | 2.38 | AT5G11320.1 | YUC4 |
| Acc02195.1 | 28.67 | 51.36 | 3.0E-02 | 1.74 | AT1G21280.1 | |
| Acc06618.1 | 1.95 | 3.80 | 3.0E-02 | 1.89 | AT5G48910.1 | LPA66 |
| Acc25709.1 | 11.17 | 26.59 | 3.1E-02 | 2.22 | AT3G23750.1 | |
| Acc03347.1 | 1.23 | 2.58 | 3.2E-02 | 1.96 | AT2G18193.1 | |
| Acc03815.1 | 1.77 | 4.18 | 3.2E-02 | 2.10 | AT4G27190.1 | |
| Acc11685.1 | 1.07 | 2.28 | 3.2E-02 | 2.01 | AT5G60900.1 | RLK1 |
| Acc10017.1 | 1.71 | 3.33 | 3.2E-02 | 1.93 | AT2G29120.1 | ATGLR2.7, GLR2.7 |
| Acc13496.1 | 0.36 | 1.11 | 3.2E-02 | 2.49 | AT5G02070.1 | |
| Acc19431.1 | 1.34 | 3.04 | 3.2E-02 | 2.13 | AT4G04890.1 | PDF2 |
| Acc33613.1 | 0.66 | 1.68 | 3.2E-02 | 2.25 | AT4G21410.1 | CRK29 |
| Acc21360.1 | 0.62 | 1.44 | 3.3E-02 | 2.17 | AT1G59740.1 | |
| Acc30419.1 | 1.05 | 2.15 | 3.3E-02 | 1.93 | AT5G12890.1 | |
| Acc19858.1 | 2.23 | 4.45 | 3.3E-02 | 1.86 | AT3G47570.1 | |
| Acc25789.1 | 1.46 | 3.14 | 3.4E-02 | 1.97 | AT2G30150.1 | |
| Acc30446.1 | 0.46 | 1.27 | 3.4E-02 | 2.43 | AT1G55230.1 | |
| Acc19634.1 | 0.57 | 1.34 | 3.5E-02 | 2.11 | AT1G03840.1 | MGP |
| Acc29312.1 | 3.03 | 6.69 | 3.5E-02 | 2.06 | AT5G08020.1 | ATRPA70B, RPA70B |
| Acc11323.1 | 0.31 | 1.12 | 3.5E-02 | 2.69 | AT4G37850.1 | |
| Acc12428.1 | 3.89 | 8.21 | 3.5E-02 | 2.00 | AT4G21070.1 | ATBRCA1, BRCA1 |
| Acc10095.1 | 1.60 | 3.22 | 3.5E-02 | 1.92 | AT3G25570.2 | |
| Acc00128.1 | 0.83 | 1.77 | 3.5E-02 | 1.96 | AT3G50120.1 | |
| Acc03298.1 | 58.98 | 10.55 | 3.5E-02 | -1.63 | AT5G53370.1 | ATPMEPCRF, PMEPCRF |
| Acc09831.1 | 1.62 | 2.88 | 3.5E-02 | 1.71 | AT4G26090.1 | RPS2 |
| Acc03323.1 | 4.07 | 6.90 | 3.5E-02 | 1.66 | AT1G43722.1 | |
| Acc11994.1 | 0.39 | 1.39 | 3.5E-02 | 2.80 | AT3G46940.1 | DUT1 |
| Acc12710.1 | 0.82 | 1.91 | 3.5E-02 | 2.10 | AT1G19900.1 | |
| Acc20597.1 | 0.43 | 1.13 | 3.5E-02 | 2.34 | AT4G21390.1 | B120 |
| Acc30063.1 | 2.10 | 3.83 | 3.6E-02 | 1.83 | AT2G27610.1 | |
| Acc07241.1 | 0.38 | 1.30 | 3.6E-02 | 2.62 | AT2G32300.1 | UCC1 |
| Acc18015.1 | 2.03 | 4.29 | 3.6E-02 | 2.01 | AT5G53150.1 | |
| Acc33656.1 | 4.66 | 0.55 | 3.6E-02 | -2.26 | AT5G59320.1 | LTP3 |
| Acc04514.1 | 0.38 | 1.18 | 3.7E-02 | 2.69 | AT5G11880.1 | |
| Acc08636.1 | 2.74 | 4.26 | 3.7E-02 | 1.52 | AT3G47570.1 | |
| Acc28628.1 | 2.43 | 4.46 | 3.7E-02 | 1.82 | AT5G25930.1 | |
| Acc25329.1 | 1.25 | 2.52 | 3.7E-02 | 1.89 | AT1G71400.1 | AtRLP12, RLP12 |
| Acc05384.1 | 0.49 | 1.26 | 3.7E-02 | 2.22 | AT5G65590.1 | |
| Acc04412.1 | 3.22 | 5.14 | 3.8E-02 | 1.58 | AT3G06880.2 | |
| Acc16281.1 | 1.87 | 3.62 | 3.8E-02 | 1.80 | AT3G16620.1 | ATTOC120, TOC120 |
| Acc17307.1 | 0.50 | 1.15 | 3.8E-02 | 2.13 | AT5G46080.1 | |
| Acc20077.1 | 9.86 | 16.26 | 3.8E-02 | 1.67 | ATCG00380.1 | RPS4 |
| Acc20235.1 | 0.36 | 1.19 | 3.8E-02 | 2.60 | AT3G49070.1 | |
| Acc27062.1 | 1.87 | 3.50 | 3.8E-02 | 1.81 | AT1G02520.1 | PGP11 |
| Acc31875.1 | 0.90 | 2.44 | 3.8E-02 | 2.26 | AT5G05260.1 | CYP79A2 |
| Acc04990.1 | 0.40 | 1.19 | 3.8E-02 | 2.47 | AT3G43660.1 | |
| Acc13157.1 | 1.39 | 2.79 | 3.8E-02 | 1.90 | AT2G33680.1 | |
| Acc01412.1 | 0.81 | 1.89 | 3.8E-02 | 2.13 | AT5G52400.1 | CYP715A1 |
| Acc05462.1 | 0.58 | 1.80 | 3.8E-02 | 2.51 | AT5G58170.1 | SVL5 |
| Acc14082.1 | 2.19 | 3.73 | 3.8E-02 | 1.65 | AT1G69440.1 | AGO7, ZIP |
| Acc18080.1 | 1.24 | 3.20 | 3.8E-02 | 2.28 | AT3G48270.1 | CYP71A26 |
| Acc29175.1 | 0.82 | 1.80 | 3.9E-02 | 1.97 | AT4G31980.1 | |
| Acc19470.1 | 1.51 | 3.12 | 3.9E-02 | 2.03 | AT5G63020.1 | |
| Acc18257.2 | 1.21 | 2.68 | 3.9E-02 | 2.05 | AT2G22070.1 | |
| Acc19282.1 | 1.64 | 3.27 | 4.0E-02 | 1.90 | AT3G15200.1 | |
| Acc07300.1 | 1.03 | 2.02 | 4.1E-02 | 1.86 | No | |
| Acc19469.1 | 1.20 | 2.14 | 4.1E-02 | 1.69 | AT5G23960.2 | TPS21 |
| Acc16807.1 | 2.03 | 4.04 | 4.1E-02 | 1.84 | AT1G21270.1 | WAK2 |
| Acc10519.1 | 1.97 | 3.92 | 4.1E-02 | 1.88 | AT2G22560.1 | |
| Acc18081.1 | 0.18 | 1.26 | 4.2E-02 | 3.69 | AT3G48310.1 | CYP71A22 |
| Acc29174.1 | 0.50 | 1.30 | 4.2E-02 | 2.24 | AT2G38290.1 | ATAMT2, AMT2;1, AMT2 |
| Acc31320.1 | 0.84 | 1.70 | 4.2E-02 | 1.91 | AT1G68330.1 | |
| Acc32344.1 | 1.12 | 2.03 | 4.2E-02 | 1.72 | AT1G07530.1 | SCL14, ATGRAS2, GRAS2 |
| Acc18158.1 | 0.64 | 1.70 | 4.3E-02 | 2.26 | AT2G26730.1 | |
| Acc17741.1 | 2.75 | 0.10 | 4.3E-02 | -3.99 | AT3G21360.1 | |
| Acc01923.1 | 1.06 | 2.61 | 4.4E-02 | 2.24 | AT3G51550.1 | FER |

| | | | | | | |
|---|---|---|---|---|---|---|
| Acc10592.1 | 1.19 | 2.20 | 4.4E-02 | 1.72 | AT2G13600.1 | |
| Acc15463.1 | 1.47 | 2.58 | 4.4E-02 | 1.72 | AT1G08070.1 | OTP82 |
| Acc02987.1 | 0.97 | 2.30 | 4.5E-02 | 2.20 | AT3G47570.1 | |
| Acc06916.1 | 0.51 | 1.26 | 4.6E-02 | 2.16 | AT2G15020.1 | |
| Acc16001.1 | 1.12 | 2.43 | 4.6E-02 | 2.01 | AT3G61170.1 | |
| Acc16854.1 | 1.83 | 3.09 | 4.6E-02 | 1.69 | AT5G02390.1 | |
| Acc17596.1 | 1.04 | 2.18 | 4.6E-02 | 1.98 | AT2G29110.1 | ATGLR2.8, GLR2.8 |
| Acc26900.1 | 0.90 | 2.01 | 4.6E-02 | 2.03 | AT1G15670.1 | |
| Acc28858.1 | 0.58 | 1.90 | 4.6E-02 | 2.58 | AT4G23060.1 | IQD22 |
| Acc06113.1 | 1.01 | 1.93 | 4.7E-02 | 1.82 | AT1G79680.1 | WAKL10, ATWAKL10 |
| Acc10089.1 | 1.70 | 3.01 | 4.7E-02 | 1.72 | AT2G24130.1 | |
| Acc30737.1 | 1.47 | 3.23 | 4.7E-02 | 1.94 | AT5G15300.1 | |
| Acc04763.1 | 0.96 | 2.98 | 4.7E-02 | 2.52 | AT1G47210.2 | CYCA3;2 |
| Acc04744.1 | 0.78 | 1.64 | 4.7E-02 | 1.99 | AT2G15490.3 | UGT73B4 |
| Acc06355.1 | 0.43 | 1.21 | 4.7E-02 | 2.50 | AT1G69480.1 | |
| Acc14259.1 | 0.88 | 1.74 | 4.7E-02 | 1.83 | AT1G24430.1 | |
| Acc21115.1 | 0.82 | 2.15 | 4.7E-02 | 2.29 | AT1G49740.1 | |
| Acc31844.1 | 52.49 | 106.61 | 4.7E-02 | 1.91 | AT4G23160.1 | CRK8 |
| Acc16377.1 | 2.89 | 4.91 | 4.7E-02 | 1.66 | AT3G59140.1 | ATMRP14, MRP14, ABCC10 |
| Acc30792.1 | 0.58 | 1.33 | 4.7E-02 | 2.07 | AT1G50420.1 | SCL3, SCL-3 |
| Acc19664.1 | 1.94 | 3.45 | 4.8E-02 | 1.71 | AT3G46530.1 | RPP13 |
| Acc08803.1 | 1.65 | 3.70 | 4.8E-02 | 2.08 | AT4G08850.2 | |
| Acc00747.1 | 0.52 | 1.39 | 4.8E-02 | 2.34 | AT5G52450.1 | |
| Acc17033.1 | 2.37 | 4.07 | 4.8E-02 | 1.70 | AT3G14470.1 | |
| Acc25257.1 | 0.88 | 2.18 | 4.8E-02 | 2.24 | AT5G44440.1 | |
| Acc25621.1 | 4.37 | 8.36 | 4.8E-02 | 1.88 | AT2G38940.1 | ATPT2, PHT1;4 |
| Acc26932.1 | 0.60 | 1.53 | 4.8E-02 | 2.25 | AT1G10340.1 | |
| Acc27957.1 | 0.43 | 1.18 | 4.8E-02 | 2.38 | AT2G45220.1 | |
| Acc28859.1 | 0.43 | 1.08 | 4.8E-02 | 2.27 | AT4G11720.1 | HAP2, GCS1 |
| Acc08016.1 | 2.51 | 4.46 | 4.8E-02 | 1.77 | No | |
| Acc08349.1 | 0.80 | 1.80 | 4.9E-02 | 2.05 | AT2G46570.1 | LAC6 |
| Acc25375.1 | 1.69 | 2.80 | 4.9E-02 | 1.64 | AT1G65090.3 | |
| Acc13534.1 | 1.12 | 2.23 | 5.0E-02 | 1.92 | No | |
| Acc03461.1 | 1.23 | 2.75 | 5.0E-02 | 2.13 | AT2G34930.1 | |
| Acc12397.1 | 1.47 | 2.95 | 5.0E-02 | 1.93 | AT5G10720.1 | AHK5, CKI2, HK5 |
| Acc02259.1 | 54.88 | 109.20 | 5.0E-02 | 1.88 | AT4G23160.1 | CRK8 |
| Acc20979.1 | 0.80 | 1.59 | 5.0E-02 | 1.82 | AT5G08490.1 | |
| Acc32414.1 | 1.00 | 2.13 | 5.0E-02 | 1.96 | AT1G16150.1 | WAKL4 |
| Acc07389.1 | 4.78 | 8.81 | 5.1E-02 | 1.82 | No | |
| Acc29261.1 | 0.85 | 2.23 | 5.1E-02 | 2.25 | AT1G68330.1 | |
| Acc26057.1 | 3.14 | 6.01 | 5.1E-02 | 1.90 | AT5G54650.2 | Fh5, ATFH5 |
| Acc10380.1 | 3.56 | 5.83 | 5.1E-02 | 1.61 | AT5G66850.1 | MAPKKK5 |
| Acc13951.1 | 1.60 | 3.36 | 5.1E-02 | 1.95 | AT1G67000.1 | |
| Acc32062.1 | 0.50 | 1.31 | 5.2E-02 | 2.41 | AT2G35930.1 | PUB23 |
| Acc16812.1 | 0.69 | 1.51 | 5.3E-02 | 2.05 | AT3G47570.1 | |
| Acc01183.1 | 1.84 | 3.26 | 5.3E-02 | 1.82 | AT2G22560.1 | |
| Acc21168.1 | 2.41 | 4.17 | 5.4E-02 | 1.66 | AT1G09970.1 | LRR XI-23, RLK7 |
| Acc22633.1 | 0.45 | 1.33 | 5.4E-02 | 2.50 | AT5G17540.1 | |
| Acc19734.1 | 0.36 | 1.12 | 5.4E-02 | 2.33 | AT2G46770.1 | NST1, ANAC043 |
| Acc00937.1 | 0.46 | 1.27 | 5.4E-02 | 2.44 | AT5G51160.1 | |
| Acc08651.1 | 0.44 | 1.35 | 5.4E-02 | 2.45 | AT5G05600.1 | |
| Acc13584.1 | 1.65 | 3.11 | 5.4E-02 | 1.77 | AT1G45616.1 | AtRLP6, RLP6 |
| Acc30947.1 | 0.62 | 1.37 | 5.4E-02 | 2.05 | AT4G37690.1 | |
| Acc00522.1 | 0.79 | 1.84 | 5.5E-02 | 2.09 | AT4G37680.2 | HHP4 |
| Acc29493.1 | 0.59 | 1.41 | 5.5E-02 | 2.24 | AT4G19230.1 | CYP707A1 |
| Acc15722.1 | 0.46 | 1.14 | 5.6E-02 | 2.25 | AT4G25470.1 | CBF2, DREB1C, FTQ4, ATCBF2 |
| Acc16608.1 | 0.37 | 1.04 | 5.6E-02 | 2.30 | AT4G14130.1 | XTR7, XTH15 |
| Acc21016.1 | 0.48 | 1.21 | 5.6E-02 | 2.14 | AT1G30670.1 | |
| Acc24394.1 | 2.33 | 4.29 | 5.6E-02 | 1.80 | AT1G20560.1 | AAE1 |
| Acc28836.1 | 1.36 | 2.72 | 5.6E-02 | 1.91 | AT5G41315.1 | GL3, MYC6.2 |
| Acc11484.1 | 1.05 | 2.20 | 5.6E-02 | 1.98 | AT4G08850.1 | |
| Acc18720.1 | 1.25 | 2.36 | 5.6E-02 | 1.82 | ATMG00860.1 | ORF158 |
| Acc19434.1 | 1.48 | 2.79 | 5.6E-02 | 1.83 | AT3G28345.1 | |
| Acc23699.1 | 1.20 | 2.46 | 5.6E-02 | 1.92 | AT4G31940.1 | CYP82C4 |
| Acc15800.1 | 0.86 | 1.64 | 5.7E-02 | 1.79 | AT5G47580.1 | |
| Acc20968.1 | 0.88 | 1.68 | 5.7E-02 | 1.81 | AT1G30650.1 | WRKY14, ATWRKY14, AR411 |
| Acc22516.1 | 0.87 | 1.70 | 5.7E-02 | 1.86 | AT5G11320.1 | YUC4 |
| Acc24997.1 | 4.43 | 9.11 | 5.7E-02 | 1.97 | AT4G05520.1 | ATEHD2, EHD2 |
| Acc25945.1 | 1.23 | 2.99 | 5.7E-02 | 2.09 | AT3G45230.1 | |
| Acc29479.1 | 0.58 | 1.36 | 5.7E-02 | 2.08 | No | |
| Acc14589.1 | 0.47 | 1.22 | 5.7E-02 | 2.18 | AT5G19790.1 | RAP2.11 |
| Acc18889.1 | 0.64 | 1.56 | 5.7E-02 | 2.09 | AT4G32300.1 | SD2-5 |

| | | | | | | |
|---|---|---|---|---|---|---|
| Acc19773.1 | 0.43 | 1.29 | 5.7E-02 | 2.46 | AT3G61780.1 | emb1703 |
| Acc20832.1 | 1.17 | 3.05 | 5.7E-02 | 2.24 | AT2G36380.1 | PDR6, ATPDR6 |
| Acc21828.1 | 1.01 | 2.05 | 5.7E-02 | 1.95 | AT3G18230.1 | |
| Acc24867.1 | 0.57 | 1.36 | 5.7E-02 | 2.11 | AT2G28780.1 | |
| Acc31552.1 | 0.48 | 1.19 | 5.7E-02 | 2.14 | AT4G02090.1 | |
| Acc19149.1 | 0.70 | 2.12 | 5.9E-02 | 2.43 | AT3G26320.1 | CYP71B36 |
| Acc20782.1 | 1.00 | 2.33 | 5.9E-02 | 2.06 | AT5G44440.1 | |
| Acc27225.1 | 0.72 | 1.52 | 5.9E-02 | 2.03 | AT4G15720.1 | |
| Acc28028.1 | 0.51 | 1.19 | 5.9E-02 | 1.99 | AT2G45570.1 | CYP76C2 |
| Acc08821.1 | 1.49 | 2.85 | 5.9E-02 | 1.91 | AT2G34930.1 | |
| Acc24570.1 | 0.36 | 1.22 | 5.9E-02 | 2.53 | No | |
| Acc20421.1 | 0.46 | 1.20 | 6.0E-02 | 2.27 | AT5G16770.2 | AtMYB9, MYB9 |
| Acc25259.1 | 0.97 | 2.02 | 6.0E-02 | 1.94 | AT5G44440.1 | |
| Acc29831.1 | 0.68 | 1.54 | 6.0E-02 | 2.06 | AT5G62620.1 | |
| Acc02832.1 | 30.62 | 5.54 | 6.0E-02 | -1.57 | AT2G40140.2 | CZF1, ZFAR1, SZF2, ATSZF2 |
| Acc09901.1 | 0.42 | 1.18 | 6.0E-02 | 2.36 | AT4G28940.1 | |
| Acc23026.1 | 1.10 | 2.10 | 6.0E-02 | 1.69 | AT5G67090.1 | |
| Acc02041.1 | 0.37 | 1.21 | 6.0E-02 | 2.65 | AT5G46050.1 | ATPTR3, PTR3 |
| Acc30489.1 | 0.56 | 1.48 | 6.0E-02 | 2.23 | AT1G55570.1 | sks12 |
| Acc17023.1 | 0.40 | 1.01 | 6.1E-02 | 2.27 | AT1G67856.1 | |
| Acc18919.1 | 1.38 | 2.22 | 6.1E-02 | 1.56 | AT2G21850.1 | |
| Acc23744.1 | 2.27 | 3.68 | 6.1E-02 | 1.57 | AT2G29120.1 | ATGLR2.7, GLR2.7 |
| Acc31577.1 | 0.94 | 2.54 | 6.1E-02 | 2.32 | AT4G02330.1 | ATPMEPCRB |
| Acc15927.1 | 57.67 | 107.29 | 6.2E-02 | 1.77 | No | |
| Acc24767.1 | 0.42 | 1.60 | 6.2E-02 | 2.73 | AT5G60680.1 | |
| Acc01434.1 | 0.98 | 2.20 | 6.2E-02 | 2.05 | AT4G23030.1 | |
| Acc03457.1 | 0.79 | 1.66 | 6.2E-02 | 1.93 | AT5G52400.1 | CYP715A1 |
| Acc28288.1 | 1.62 | 3.45 | 6.2E-02 | 2.01 | AT4G15020.2 | |
| Acc12378.1 | 1.38 | 2.51 | 6.2E-02 | 1.79 | AT1G40390.1 | |
| Acc22989.1 | 1.74 | 3.54 | 6.2E-02 | 1.92 | AT3G22150.1 | |
| Acc06736.1 | 4.05 | 6.78 | 6.2E-02 | 1.68 | AT4G22970.1 | RSW4, AESP, ESP |
| Acc16652.1 | 1.30 | 2.86 | 6.2E-02 | 2.07 | AT3G07290.1 | |
| Acc16928.1 | 0.32 | 1.13 | 6.2E-02 | 2.63 | AT3G26040.1 | |
| Acc26296.1 | 1.75 | 2.89 | 6.2E-02 | 1.66 | AT3G48310.1 | CYP71A22 |
| Acc00808.1 | 37.44 | 60.96 | 6.3E-02 | 1.60 | No | |
| Acc01281.1 | 1.78 | 2.76 | 6.3E-02 | 1.48 | AT5G49690.1 | |
| Acc11016.1 | 0.58 | 1.43 | 6.3E-02 | 2.19 | AT3G46530.1 | RPP13 |
| Acc18674.1 | 2.57 | 4.37 | 6.3E-02 | 1.79 | AT3G42660.1 | |
| Acc20515.1 | 0.73 | 1.60 | 6.3E-02 | 2.15 | AT5G56310.1 | |
| Acc27465.1 | 0.40 | 1.38 | 6.3E-02 | 2.55 | No | |
| Acc19277.1 | 0.75 | 1.99 | 6.4E-02 | 2.35 | AT3G07800.1 | |
| Acc00829.1 | 4.83 | 7.62 | 6.4E-02 | 1.56 | AT1G21280.1 | |
| Acc17656.1 | 43.57 | 7.25 | 6.4E-02 | -1.71 | AT5G07990.1 | TT7, CYP75B1, D501 |
| Acc19460.1 | 1.01 | 1.80 | 6.4E-02 | 1.67 | AT4G27290.1 | |
| Acc20278.1 | 0.69 | 1.56 | 6.4E-02 | 2.01 | AT4G13420.1 | HAK5, ATHAK5 |
| Acc20281.1 | 0.65 | 1.73 | 6.4E-02 | 2.29 | AT5G43360.1 | PHT3, ATPT4, PHT1;3 |
| Acc21382.1 | 1.96 | 3.10 | 6.4E-02 | 1.59 | AT3G47570.1 | |
| Acc32231.1 | 1.00 | 2.18 | 6.4E-02 | 2.00 | AT3G45260.1 | |
| Acc33485.1 | 0.42 | 1.19 | 6.4E-02 | 2.43 | AT1G79180.1 | ATMYB63, MYB63 |
| Acc15384.1 | 1.48 | 2.78 | 6.5E-02 | 1.89 | AT3G18670.1 | |
| Acc01683.1 | 0.81 | 1.74 | 6.6E-02 | 1.89 | AT2G43130.1 | TRAB11F, ATRABA5C, ARA-4, RABA5C ... 357 |
| Acc01459.1 | 2.57 | 0.21 | 6.6E-02 | -2.76 | AT5G62360.1 | |
| Acc24599.1 | 1.61 | 2.93 | 6.6E-02 | 1.79 | No | |
| Acc08654.1 | 1.22 | 2.55 | 6.6E-02 | 1.96 | AT5G58630.1 | |
| Acc23783.1 | 1.08 | 2.22 | 6.6E-02 | 1.81 | AT2G19130.1 | |
| Acc00812.1 | 0.73 | 1.72 | 6.6E-02 | 2.20 | AT5G15900.1 | TBL19 |
| Acc23585.1 | 0.90 | 2.07 | 6.6E-02 | 2.15 | AT4G31500.1 | CYP83B1, SUR2, RNT1, RED1, ATR4 |
| Acc30672.1 | 0.64 | 2.89 | 6.7E-02 | 3.01 | AT3G28470.1 | ATMYB35, TDF1 |
| Acc19290.1 | 3.08 | 5.40 | 6.7E-02 | 1.66 | AT1G55610.2 | BRL1 |
| Acc31543.1 | 0.92 | 1.83 | 6.8E-02 | 1.89 | AT1G02570.1 | |
| Acc18446.1 | 0.36 | 1.19 | 6.8E-02 | 2.72 | No | |
| Acc19700.1 | 1.18 | 2.15 | 6.8E-02 | 1.73 | AT4G02750.1 | |
| Acc23200.1 | 0.55 | 1.57 | 6.9E-02 | 2.50 | AT5G43810.2 | ZLL |
| Acc31243.1 | 0.84 | 1.59 | 6.9E-02 | 1.89 | AT5G59810.1 | ATSBT5.4, SBT5.4 |
| Acc33324.1 | 2.35 | 4.85 | 6.9E-02 | 2.04 | AT1G50700.1 | CPK33 |
| Acc11544.2 | 1.32 | 2.44 | 6.9E-02 | 1.77 | AT4G08850.1 | |
| Acc32635.1 | 0.64 | 1.69 | 6.9E-02 | 2.28 | AT3G25810.1 | |
| Acc02759.1 | 0.38 | 1.08 | 6.9E-02 | 2.35 | AT1G28710.3 | |
| Acc03921.1 | 1.14 | 1.86 | 7.0E-02 | 1.61 | AT1G74630.1 | |
| Acc17432.1 | 0.46 | 1.15 | 7.0E-02 | 2.24 | AT4G23280.1 | CRK20 |
| Acc08842.1 | 1.48 | 2.83 | 7.0E-02 | 1.81 | No | |
| Acc31973.1 | 0.71 | 1.43 | 7.0E-02 | 1.90 | AT1G65450.1 | |

| | | | | | | |
|---|---|---|---|---|---|---|
| Acc09520.1 | 0.81 | 1.93 | 7.1E-02 | 2.06 | AT4G25810.1 | XTR6, XTH23 |
| Acc31544.1 | 0.72 | 1.43 | 7.1E-02 | 1.85 | AT3G03450.1 | RGL2 |
| Acc00749.1 | 1.96 | 3.71 | 7.2E-02 | 1.83 | AT1G21280.1 | |
| Acc18985.1 | 1.90 | 3.28 | 7.2E-02 | 1.67 | AT4G28310.1 | |
| Acc29172.1 | 4.75 | 11.32 | 7.2E-02 | 2.22 | AT4G14770.1 | TCX2, ATTCX2 |
| Acc12262.1 | 0.42 | 1.14 | 7.2E-02 | 2.38 | AT3G18690.1 | MKS1 |
| Acc13355.1 | 0.43 | 1.06 | 7.2E-02 | 2.18 | AT2G35930.1 | PUB23 |
| Acc11630.1 | 0.55 | 1.30 | 7.2E-02 | 2.08 | AT5G09360.1 | LAC14 |
| Acc20786.1 | 1.19 | 1.83 | 7.2E-02 | 1.57 | AT3G19050.1 | POK2 |
| Acc19887.1 | 0.56 | 1.34 | 7.3E-02 | 2.04 | AT1G02030.1 | |
| Acc26652.1 | 0.98 | 1.60 | 7.3E-02 | 1.61 | AT5G60970.1 | TCP5 |
| Acc18731.1 | 0.85 | 1.81 | 7.3E-02 | 1.98 | No | |
| Acc19867.1 | 2.89 | 4.45 | 7.3E-02 | 1.49 | AT3G23330.1 | |
| Acc01255.1 | 0.76 | 1.41 | 7.3E-02 | 1.84 | AT4G39490.1 | CYP96A10 |
| Acc20808.1 | 0.56 | 1.13 | 7.3E-02 | 1.93 | AT5G44360.1 | |
| Acc04081.1 | 1.96 | 3.42 | 7.5E-02 | 1.71 | No | |
| Acc16028.1 | 1.12 | 3.11 | 7.5E-02 | 2.43 | AT3G13980.1 | |
| Acc30466.1 | 0.47 | 1.21 | 7.5E-02 | 2.24 | AT5G19790.1 | RAP2.11 |
| Acc04135.1 | 1.82 | 3.51 | 7.5E-02 | 1.92 | AT4G02750.1 | |
| Acc24071.1 | 0.54 | 1.50 | 7.5E-02 | 2.35 | AT1G76980.1 | |
| Acc27872.1 | 0.87 | 2.22 | 7.6E-02 | 2.25 | AT5G36110.1 | CYP716A1 |
| Acc02118.1 | 1.19 | 2.71 | 7.6E-02 | 2.02 | AT3G52970.1 | CYP76G1 |
| Acc13452.1 | 0.50 | 1.12 | 7.6E-02 | 2.01 | AT2G36450.1 | HRD |
| Acc23766.1 | 0.52 | 1.13 | 7.6E-02 | 2.01 | AT2G36770.1 | |
| Acc19114.1 | 1.03 | 2.02 | 7.6E-02 | 1.85 | AT2G24720.1 | ATGLR2.2, GLR2.2 |
| Acc14662.1 | 2.32 | 4.62 | 7.8E-02 | 1.94 | AT1G04020.1 | ATBARD1, BARD1, ROW1 |
| Acc15716.1 | 0.86 | 1.42 | 7.8E-02 | 1.61 | AT4G23030.1 | |
| Acc32319.1 | 0.54 | 1.22 | 7.9E-02 | 2.04 | AT2G28780.1 | |
| Acc11587.1 | 2.89 | 3.81 | 7.9E-02 | 1.30 | AT1G76770.1 | |
| Acc13287.1 | 1.10 | 2.08 | 7.9E-02 | 1.75 | AT5G39660.2 | CDF2 |
| Acc04228.1 | 1.01 | 1.94 | 7.9E-02 | 1.80 | AT3G63380.1 | |
| Acc05127.1 | 2.21 | 3.43 | 7.9E-02 | 1.52 | AT1G33590.1 | |
| Acc31103.1 | 0.49 | 1.42 | 7.9E-02 | 2.41 | AT1G71250.1 | |
| Acc00627.1 | 3.33 | 6.12 | 7.9E-02 | 1.79 | AT1G21280.1 | |
| Acc04488.1 | 2.13 | 3.93 | 7.9E-02 | 1.81 | AT1G20640.2 | |
| Acc04824.1 | 2.53 | 4.54 | 8.0E-02 | 1.73 | AT5G49660.1 | |
| Acc11325.1 | 1.61 | 3.14 | 8.0E-02 | 1.91 | AT3G17450.1 | |
| Acc32470.1 | 0.75 | 1.58 | 8.1E-02 | 1.90 | AT1G51190.1 | PLT2 |
| Acc04295.1 | 0.40 | 1.03 | 8.2E-02 | 2.25 | AT1G03220.1 | |
| Acc14673.1 | 2.48 | 4.01 | 8.2E-02 | 1.55 | AT4G28650.1 | |
| Acc09736.1 | 1.07 | 1.90 | 8.2E-02 | 1.67 | AT1G62280.1 | SLAH1 |
| Acc02263.1 | 1.66 | 3.39 | 8.2E-02 | 1.96 | AT1G67780.1 | |
| Acc01138.1 | 0.60 | 1.44 | 8.2E-02 | 2.01 | AT1G66120.1 | |
| Acc06298.1 | 1.31 | 2.38 | 8.2E-02 | 1.75 | AT1G69870.1 | NRT1.7 |
| Acc20204.1 | 1.73 | 2.97 | 8.2E-02 | 1.74 | AT4G33990.1 | EMB2758 |
| Acc09287.1 | 3.02 | 4.15 | 8.2E-02 | 1.33 | AT3G14470.1 | |
| Acc03770.1 | 0.65 | 1.37 | 8.2E-02 | 1.90 | AT3G27580.2 | ATPK7 |
| Acc08829.1 | 1.55 | 3.35 | 8.2E-02 | 2.08 | No | |
| Acc22622.1 | 2.17 | 4.22 | 8.3E-02 | 1.90 | AT4G19440.2 | |
| Acc27885.1 | 1.12 | 2.13 | 8.3E-02 | 1.80 | AT5G43360.1 | PHT3, ATPT4, PHT1;3 |
| Acc23786.1 | 0.39 | 1.05 | 8.3E-02 | 2.46 | AT4G37370.1 | CYP81D8 |
| Acc32715.1 | 2.48 | 3.98 | 8.3E-02 | 1.61 | AT5G13150.1 | ATEXO70C1, EXO70C1 |
| Acc16460.1 | 2.16 | 3.12 | 8.3E-02 | 1.41 | AT2G31270.1 | ATCDT1A, CDT1A, CDT1 |
| Acc17748.1 | 0.50 | 1.12 | 8.4E-02 | 2.00 | AT1G79760.1 | DTA4 |
| Acc24986.1 | 0.30 | 1.01 | 8.4E-02 | 2.63 | AT2G43870.1 | |
| Acc04862.1 | 0.93 | 2.10 | 8.4E-02 | 2.03 | AT1G58120.1 | |
| Acc21114.1 | 0.66 | 1.45 | 8.4E-02 | 2.02 | AT1G49780.1 | PUB26 |
| Acc08561.1 | 0.92 | 1.90 | 8.4E-02 | 1.91 | AT2G39490.1 | |
| Acc19794.1 | 1.26 | 3.01 | 8.4E-02 | 2.10 | AT1G01430.1 | TBL25 |
| Acc28585.1 | 1.32 | 2.48 | 8.4E-02 | 1.84 | AT5G25620.1 | YUC6 |
| Acc18516.1 | 0.43 | 2.02 | 8.5E-02 | 3.12 | AT3G47090.1 | |
| Acc32625.1 | 20.33 | 3.43 | 8.6E-02 | -1.81 | AT3G57950.1 | |
| Acc27426.1 | 3.46 | 4.72 | 8.6E-02 | 1.33 | AT5G16250.1 | |
| Acc09649.1 | 0.53 | 1.21 | 8.6E-02 | 2.11 | AT4G27290.1 | |
| Acc13405.1 | 3.25 | 4.70 | 8.6E-02 | 1.42 | AT1G03400.1 | |
| Acc33099.1 | 4.17 | 7.75 | 8.6E-02 | 1.85 | AT4G15890.1 | |
| Acc13118.1 | 0.69 | 1.47 | 8.6E-02 | 1.98 | AT5G43810.2 | ZLL |
| Acc13463.1 | 0.96 | 1.70 | 8.6E-02 | 1.78 | AT5G43630.1 | TZP |
| Acc01931.1 | 1.25 | 2.33 | 8.7E-02 | 1.81 | AT3G51550.1 | FER |
| Acc12267.1 | 1.77 | 3.00 | 8.7E-02 | 1.58 | AT1G65610.1 | ATGH9A2, KOR2 |
| Acc04090.1 | 0.74 | 2.07 | 8.7E-02 | 2.37 | AT5G15240.1 | |
| Acc06058.1 | 0.78 | 1.77 | 8.7E-02 | 2.18 | AT2G45570.1 | CYP76C2 |

| | | | | | | |
|---|---|---|---|---|---|---|
| Acc02319.1 | 0.93 | 2.19 | 8.7E-02 | 2.21 | AT3G16520.3 | UGT88A1 |
| Acc24204.1 | 0.79 | 1.33 | 8.7E-02 | 1.64 | AT1G04500.1 | |
| Acc26190.1 | 0.63 | 1.22 | 8.7E-02 | 1.92 | AT4G32300.1 | SD2-5 |
| Acc18627.1 | 2.28 | 3.18 | 8.8E-02 | 1.37 | AT4G18750.1 | DOT4 |
| Acc19738.1 | 0.85 | 1.64 | 8.9E-02 | 1.84 | AT4G38180.1 | FRS5 |
| Acc30852.1 | 0.58 | 1.23 | 8.9E-02 | 1.98 | AT4G05200.1 | CRK25 |
| Acc18592.1 | 1.01 | 2.15 | 9.0E-02 | 1.94 | AT3G29785.1 | |
| Acc14251.1 | 0.70 | 1.36 | 9.0E-02 | 1.79 | AT1G24430.1 | |
| Acc28313.1 | 0.84 | 1.86 | 9.0E-02 | 1.98 | No | |
| Acc29973.1 | 0.79 | 1.52 | 9.0E-02 | 1.86 | AT5G46050.1 | ATPTR3, PTR3 |
| Acc18077.1 | 1.53 | 3.53 | 9.0E-02 | 2.10 | AT3G47570.1 | |
| Acc16490.1 | 3.25 | 4.71 | 9.1E-02 | 1.46 | AT3G18020.1 | |
| Acc11309.1 | 0.70 | 1.37 | 9.1E-02 | 1.89 | AT1G23770.1 | |
| Acc25493.1 | 0.75 | 1.69 | 9.1E-02 | 2.08 | AT2G47540.1 | |
| Acc20215.1 | 111.63 | 14.62 | 9.1E-02 | -2.11 | AT3G61440.1 | ATCYSC1, ARATH;BSAS3;1, CYSC1 |
| Acc12240.1 | 1.54 | 2.57 | 9.2E-02 | 1.65 | AT4G08850.1 | |
| Acc19538.1 | 22.84 | 4.69 | 9.2E-02 | -1.45 | AT4G14860.1 | atofp11, OFP11 |
| Acc22653.1 | 2.76 | 4.03 | 9.2E-02 | 1.36 | AT3G45900.1 | |
| Acc31510.1 | 1.05 | 2.06 | 9.2E-02 | 1.82 | No | |
| Acc31716.1 | 6.00 | 9.57 | 9.2E-02 | 1.51 | AT1G21280.1 | |
| Acc09654.1 | 1.71 | 2.87 | 9.2E-02 | 1.72 | AT1G11330.1 | |
| Acc27283.1 | 0.60 | 1.21 | 9.2E-02 | 1.93 | AT5G19040.1 | ATIPT5, IPT5 |
| Acc25324.1 | 0.64 | 1.74 | 9.3E-02 | 2.28 | AT3G47570.1 | |
| Acc03401.1 | 0.42 | 1.04 | 9.4E-02 | 2.19 | AT2G46150.1 | |
| Acc15657.1 | 1.60 | 2.41 | 9.5E-02 | 1.46 | AT5G41410.1 | BEL1 |
| Acc11826.1 | 7.23 | 13.24 | 9.6E-02 | 1.81 | AT1G75150.1 | |
| Acc02334.1 | 0.69 | 1.26 | 9.6E-02 | 1.71 | AT5G61430.1 | ANAC100, ATNAC5, NAC100 |
| Acc07299.1 | 0.77 | 1.34 | 9.7E-02 | 1.67 | AT1G75540.1 | STH2 |
| Acc23985.1 | 1.19 | 2.23 | 9.7E-02 | 1.86 | AT5G48100.1 | TT10, LAC15, ATLAC15 |
| Acc06946.1 | 0.52 | 1.32 | 9.7E-02 | 2.24 | AT5G67360.1 | ARA12 |
| Acc05044.1 | 1.75 | 3.30 | 9.7E-02 | 1.80 | AT1G11050.1 | |
| Acc15540.1 | 0.58 | 1.43 | 9.8E-02 | 2.18 | AT1G64625.2 | |
| Acc23789.1 | 0.67 | 1.29 | 9.8E-02 | 1.78 | AT3G47570.1 | |
| Acc14237.1 | 0.58 | 1.37 | 9.8E-02 | 2.05 | AT3G26120.1 | TEL1 |
| Acc15287.1 | 3.90 | 6.49 | 9.8E-02 | 1.71 | AT4G02110.1 | |
| Acc27873.1 | 2.59 | 5.17 | 9.8E-02 | 1.87 | AT5G05240.1 | |
| Acc31723.1 | 10.30 | 2.00 | 9.8E-02 | -1.59 | AT5G43860.1 | ATCLH2, CLH2 |
| Acc14653.1 | 0.59 | 1.36 | 9.9E-02 | 2.15 | AT3G54220.1 | SCR, SGR1 |
| Acc26607.1 | 0.87 | 1.64 | 9.9E-02 | 1.76 | No | |
| Acc32006.1 | 1.15 | 1.94 | 9.9E-02 | 1.59 | AT3G24240.1 | |

**Supplemental Table S3 GO enrichment analysis in the DEGs between male and female anthers**

| GO | Description | p-value | FDR |
|---|---|---|---|
| GO:0006468 | protein phosphorylation | 1.E-08 | 7.E-06 |
| GO:0016310 | phosphorylation | 3.E-06 | 1.E-03 |
| GO:0012501 | programmed cell death | 7.E-06 | 2.E-03 |
| GO:0008219 | cell death | 5.E-05 | 1.E-02 |
| GO:0007167 | enzyme linked receptor protein signaling pathway | 2.E-04 | 2.E-02 |
| GO:0007169 | transmembrane receptor protein tyrosine kinase signaling pathway | 2.E-04 | 2.E-02 |
| GO:0006796 | phosphate-containing compound metabolic process | 2.E-04 | 2.E-02 |
| GO:0006793 | phosphorus metabolic process | 2.E-04 | 2.E-02 |

**Supplemental Table S4 Characterization of gene-edited *Arabidopsis* and *N. tabacum* lines**

| line name | species | gene-editing target | chimeric edition | complementation with *Actinidia FrBy* | pollen germination[1] | gynoecium development[2] |
|---|---|---|---|---|---|---|
| Atnull #5 | *Arabidopsis* | *AT1G30800* | | - | - | +++ |
| Atnull #2 | *Arabidopsis* | *AT1G30800* | X | - | + | +++ |
| AtCont 1 | *Arabidopsis* | NA | | - | +++ | +++ |
| AtCont 2 | *Arabidopsis* | NA | | - | +++ | +++ |
| Ntnull #1 | *N. tabacum* | *FAS1* | X | - | +++ | +++ |
| Ntnull #2 | *N. tabacum* | *FAS1* | X | - | +++ | +++ |
| Ntnull #3 | *N. tabacum* | *FAS1* | | - | - | +++ |
| Ntnull #4 | *N. tabacum* | *FAS1* | | - | ± | +++ |
| Ntnull #5 | *N. tabacum* | *FAS1* | X | - | +++ | +++ |
| NtCont 1 | *N. tabacum* | NA | | - | +++ | +++ |
| NtCont 2 | *N. tabacum* | NA | | - | +++ | +++ |
| Ntnull #3-FrBy #1 | *N. tabacum* | *FAS1* | | X | ++ | +++ |
| Ntnull #3-FrBy #2 | *N. tabacum* | *FAS1* | | X | ++ | +++ |
| Ntnull #3-FrBy #3 | *N. tabacum* | *FAS1* | | X | + | +++ |

[1] Based on pollen germination ratio and pollen tube vigor.

[2] Based on size and seed production.

**Supplemental Table S5 Summary of the draft genome assembly in cv. Soyu by 10X Genomics data**

| Input data | Our data | Ideal values according to Supernova |
|---|---|---|
| reads (M) | 632.09 | |
| mean read length (bp) | 139.5 | 140 |
| effective read coverage (X) | 103.66 | ~42 for nominal 56X cov. |
| faction of Q30 bases in read (%) | 77.52% | 75-85 |
| median insert size (bp) | 327 | 350-400 |
| fraction of proper read pairs (%) | 89.2 | >75 |
| weighted mean mol size (kbp) | 40.49 | 50-100 |
| fraction of reads not barcoded (%) | 4.96 | |
| phased reads (nonduplicate) (%) | 52.95 | 45-50 |
| **Output data** | | |
| phaseblock N50 (kbp) | 320.93 | |
| Scaffold N50 (kbp) | 318.21 | |
| Assembly size (only scaffolds >10kb) (Mbp) | 492.33 | |
| Total assembly size (Mbp) | 720.99 | |
| Total coverage (%) | 95.1 | |

**Supplemental Table S6 List of the assembled scaffolds of cv. Soyu anchored by Y-specific contigs**

| cv. Soyu scaffolds ID | length (bp) | Nos. of Y-specific contigs anchored to Soyu scaffolds[1] | Length (bp) per anchored Y-specific contigs[2] |
|---|---|---|---|
| Contig8318 | 34527 | 9 | 3836 |
| Contig426301 | 579725 | 92 | 6301 |
| Contig7598 | 241132 | 27 | 8931 |
| Contig423383 | 53732 | 8 | 6717 |
| Contig5670 | 135918 | 14 | 9708 |
| Contig8003 | 31425 | 3 | 10475 |
| Contig422073 | 67478 | 6 | 11246 |
| Contig6851 | 58752 | 5 | 11750 |
| Contig6898 | 173508 | 10 | 17351 |
| Contig426119 | 111857 | 2 | 55929 |
| Contig6906 | 142988 | 2 | 71494 |
| Contig6077 | 219867 | 3 | 73289 |
| Contig6355 | 286844 | 3 | 95615 |
| Contig7303 | 265233 | 2 | 132617 |
| Contig7659 | 429366 | 3 | 143122 |
| Contig6712 | 338516 | 2 | 169258 |
| Contig7674 | 346861 | 2 | 173431 |
| Contig8164 | 445483 | 2 | 222742 |
| Contig7978 | 482689 | 2 | 241345 |
| Contig7591 | 703481 | 2 | 351741 |
| Contig6823 | 1378423 | 2 | 689212 |

[1] The Y-specific contigs in *A. chinensis* were derived from male-specific kmer-cataloging in Akagi et al. (2018) (15)

[2] The scaffolds where the Y-specific contigs were frequently anchored (length per anchored contigs < 20-kb), highlighted in gray ($N$ = 9), were used for further anchoring analyses with gem-indexes.

### Supplemental Table S7 List of genes predicted in the 9 scaffolds anchored by the Y-specific contigs

| Scaffold | Gene IDs | Gene name | Male specificity[1] | Functional annotation by TAIR | | | Functional annotation by nr database[2] [species] |
|---|---|---|---|---|---|---|---|
| | | | | Gene ID | Abbreviation | putative function | |
| Scaffold I | 6851:g1.t1 | | P | AT3G14470.1 | | NB-ARC domain-containing disease resistance | |
| | 6851:g2.t1 | | P | | | | |
| | 6851:g3.t1 | | P | | | | |
| | 6851:g4.t1 | | P | | | | |
| | 6851:g5.t1 | | P | | | | |
| | 6851:g6.t1 | | P | AT1G18750.1 | AGL65 | AGAMOUS-like 65 | |
| | 6851:g7.t1 | | R | | | | |
| | 6851:g8.t1 | | R | | | | |
| Scaffold II | 8318:g9.t1 | | R | | | | |
| | 8318:g10.t1 | | R | ATCG00490.1 | RBCL | ribulose-bisphosphate carboxylase-like | |
| | 8318:g11.t1 | | R | | | | |
| Scaffold III | 6898:g12.t1 | | R | | | | |
| | 6898:g13.t1 | | R | AT2G01905.1 | CYCJ18 | cyclin J18 | |
| | 6898:g14.t1 | | P | ATMG00860.1 | ORF158 | DNA/RNA polymerases superfamily | |
| | 6898:g15.t1 | | P | | | | |
| | 6898:g16.t1 | | P | | | | |
| | 6898:g17.t1 | | P | AT2G01900.1 | | DNAse I-like superfamily protein | |
| | 6898:g18.t1 | | P | | | | |
| | 6898:g19.t1 | | P | | | | |
| | 6898:g20.t1 | | P | | | | |
| | 6898:g21.t1 | | P | | | | |
| | 6898:g22.t1 | | P | | | | |
| | 6898:g23.t1 | | P | | | | |
| | 6898:g24.t1 | | R | | | | |
| | 6898:g25.t1 | | R | | | | |
| | 6898:g26.t1 | | P | AT3G24255.1 | | RNA-directed DNA polymerase | |
| Scaffold IV | 423383:g27.t1 | | R | | | | |
| | 423383:g28.t1 | | R | | | | |
| | 423383:g29.t1 | | R | ATMG00860.1 | ORF158 | DNA/RNA polymerases superfamily | |
| | 423383:g30.t1 | | R | | | | |
| | 423383:g31.t1 | | R | | | | |
| | 423383:g32.t1 | | M | | | | Platelet-derived growth factor receptor beta like [Actinidia chinensis] |
| | 423383:g33.t1 | ***Shy Girl*** | M | AT5G26594.1 | ARR24, RR24 | response regulator 24 | Shy Girl [Actinidia chinensis] |
| | 423383:g34.t1 | | M | | | | Retrovirus-related Pol polyprotein from transposon TNT 1-94 [Vitis vinifera] |
| Scaffold V | 426301:g91.t1 | | R | | | | |
| | 426301:g90.t1 | | P | | | | |
| | 426301:g89.t1 | | M | | | | Retrovirus-related Pol polyprotein from transposon 17.6 [Vitis vinifera] |
| | 426301:g88.t1 | | M | | | | Retrovirus-related Pol polyprotein from transposon 297 [Vitis vinifera] |
| | 426301:g87.t1 | | M | | | | no hits |
| | 426301:g86.t1 | | R | | | | |
| | 426301:g85.t1 | | M | | | | retrotransposon-related, gag protein [Populus trichocarpa] |
| | 426301:g84.t1 | | M | | | | Cortactin-binding protein like [Actinidia chinensis var. chinensis] |
| | 426301:g83.t1 | | R | | | | |
| | 426301:g82.t1 | ***YFT*** | M | AT1G65480.1 | FT | PEBP (phosphatidylethanolamine-binding) | FT2 [Actinidia chinensis var. chinensis] |
| | 426301:g81.t1 | | M | ATMG00860.1 | ORF158 | DNA/RNA polymerases superfamily | putative transposase, Ptta/En/Spm [Actinidia chinensis var. chinensis] |
| | 426301:g80.t1 | | M | | | | transposon-related Gag-Pol polyprotein [Vitis vinifera] |
| | 426301:g79.t1 | | M | | | | no hits |

| | | | | | | | |
|---|---|---|---|---|---|---|---|
| Scaffold V | 426301:g78.t1 | | R | ATMG00750.1 | ORF119 | GAG/POL/ENV polyprotein | |
| | 426301:g77.t1 | | R | | | | |
| | 426301:g76.t1 | | R | | | | |
| | 426301:g75.t1 | | R | AT5G28780.1 | | PIF1 helicase | |
| | 426301:g74.t1 | | M | | | | Polyprotein, putative [Solanum demissum] |
| | 426301:g73.t1 | | R | | | | |
| | 426301:g72.t1 | | M | | | | TRNA modification GTPase [Actinidia chinensis var. chinensis] |
| | 426301:g71.t1 | | M | | | | Retrovirus-related Pol polyprotein from transposon 17.6 [Vitis vinifera] |
| | 426301:g70.t1 | | M | | | | Immunoglobulin superfamily member 8 like [Actinidia chinensis var. chinensis] |
| | 426301:g69.t1 | | R | | | | |
| | 426301:g68.t1 | | R | ATMG00860.1 | ORF158 | DNA/RNA polymerases superfamily | |
| | 426301:g67.t1 | | R | AT4G23160.1 | CRK8 | cysteine-rich RLK | |
| | 426301:g66.t1 | | R | | | | |
| | 426301:g65.t1 | | R | | | | |
| | 426301:g64.t1 | | P | | | | |
| | 426301:g63.t1 | | P | | | | |
| | 426301:g62.t1 | | R | | | | |
| | 426301:g61.t1 | | R | | | | |
| | 426301:g60.t1 | | R | | | | |
| | 426301:g59.t1 | | R | | | | |
| | 426301:g58.t1 | | M | | | | no hit |
| | 426301:g57.t1 | | M | | | | Retrovirus-related Pol polyprotein from transposon 17.6 [Vitis vinifera] |
| | 426301:g56.t1 | ***Friendly Boy*** | M | AT1G30800.1 | | Fasciclin-like arabinogalactan family | FAS1 domain-containing protein [Cynara cardunculus var. scolymus] |
| | 426301:g55.t1 | | R | | | | |
| | 426301:g54.t1 | | R | | | | |
| | 426301:g53.t1 | | R | | | | |
| | 426301:g52.t1 | | R | AT1G43260.1 | | hAT transposon superfamily protein | |
| | 426301:g51.t1 | | M | | | | no hit |
| | 426301:g50.t1 | | R | | | | |
| | 426301:g49.t1 | | R | | | | |
| | 426301:g48.t1 | | R | | | | |
| | 426301:g47.t1 | | R | | | | |
| | 426301:g46.t1 | | M | | | | Transposon Ty3-G Gag-Pol polyprotein [Cajanus cajan] |
| | 426301:g45.t1 | | M | | | | Stress response protein [Actinidia chinensis var. chinensis] |
| | 426301:g44.t1 | | M | | | | Gag-Pol polyprotein [Vitis vinifera] |
| | 426301:g43.t1 | | R | AT3G01410.2 | | Polynucleotidyl transferase | |
| | 426301:g42.t1 | | R | | | | |
| | 426301:g41.t1 | | R | | | | |
| | 426301:g40.t1 | | P | | | | |
| | 426301:g39.t1 | | P | | | | |
| | 426301:g38.t1 | | P | | | | |
| | 426301:g37.t1 | | P | | | | |
| | 426301:g36.t1 | | R | | | | |
| | 426301:g35.t1 | | R | ATMG00860.1 | ORF158 | DNA/RNA polymerases superfamily | |
| Scaffold VI | 7598:g92.t1 | | P | | | | |
| | 7598:g93.t1 | | R | | | | |
| | 7598:g94.t1 | | P | | | | |
| | 7598:g95.t1 | | R | | | | |
| | 7598:g96.t1 | | P | AT1G48720.1 | | unknown protein; Has 229 | |
| | 7598:g97.t1 | | P | | | | |
| | 7598:g98.t1 | | P | | | | |

| | | | | | | | | |
|---|---|---|---|---|---|---|---|---|
| Scaffold Ⅵ | 7598:g99.t1 | M | | | | | | Mutator-like transposase; 53847-56139 [Arabidopsis thaliana] |
| | 7598:g100.t1 | M | | | | | | Retrovirus-related Pol polyprotein from transposon TNT 1-94 [Vitis vinifera] |
| | 7598:g101.t1 | R | AT1G59960.1 | | NAD(P)-linked oxidoreductase superfamily | | | |
| | 7598:g102.t1 | M | | | | | | Serine/arginine repetitive matrix protein [Actinidia chinensis var. chinensis] |
| | 7598:g103.t1 | M | ATMG00860 | | DNA/RNA polymerase superfamily | | | reverse transcriptase-like protein, partial [Escherichia coli] |
| | 7598:g104.t1 | R | | | | | | |
| | 7598:g105.t1 | R | | | | | | |
| | 7598:g106.t1 | R | ATMG00860.1 | ORF158 | DNA/RNA polymerases superfamily | | | |
| | 7598:g107.t1 | R | | | | | | |
| | 7598:g108.t1 | R | | | | | | |
| | 7598:g109.t1 | M | | | | | | Capsular polysaccharide phosphotransferase [Actinidia chinensis var. chinensis] |
| | 7598:g110.t1 | R | | | | | | |
| | 7598:g111.t1 | R | | | | | | |
| | 7598:g112.t1 | R | AT1G59950.1 | | NAD(P)-linked oxidoreductase superfamily | | | |
| | 7598:g113.t1 | P | AT1G59960.1 | | NAD(P)-linked oxidoreductase superfamily | | | |
| | 7598:g114.t1 | P | | | | | | |
| | 7598:g115.t1 | P | | | | | | |
| | 7598:g116.t1 | P | | | | | | |
| | 7598:g117.t1 | P | | | | | | |
| | g118 | P | | | | | | |
| | 7598:g119.t1 | P | | | | | | |
| | 7598:g120.t1 | P | | | | | | |
| | 7598:g121.t1 | R | AT4G23160.1 | CRK8 | cysteine-rich RLK | | | |
| | 7598:g122.t1 | R | | | | | | |
| | 7598:g123.t1 | P | AT1G59960.1 | | NAD(P)-linked oxidoreductase superfamily | | | |
| Scaffold Ⅶ | 422073:g124.t1 | R | | | | | | |
| | 422073:g125.t1 | M | | | | | | Retrovirus-related Pol polyprotein from transposon 17.6 [Vitis vinifera] |
| | 422073:g126.t1 | R | | | | | | |
| | 422073:g127.t1 | R | | | | | | |
| | 422073:g128.t1 | R | ATMG00860.1 | ORF158 | DNA/RNA polymerases superfamily... | 103 | 1e-25 | |
| | 422073:g129.t1 | R | | | | | | |
| | 422073:g130.t1 | R | | | | | | |
| Scaffold Ⅷ | 5670:g131.t1 | P | | | | | | |
| | 5670:g132.t1 | P | | | | | | |
| | 5670:g133.t1 | P | | | | | | |
| | 5670:g134.t1 | P | | | | | | |
| | 5670:g135.t1 | R | | | | | | |
| | 5670:g136.t1 | P | | | | | | |
| | 5670:g137.t1 | P | | | | | | |
| | 5670:g138.t1 | R | | | | | | |
| | 5670:g139.t1 | P | AT5G01670.1 | | NAD(P)-linked oxidoreductase superfamily | | | |
| | 5670:g140.t1 | P | AT2G37770.2 | | NAD(P)-linked oxidoreductase superfamily | | | |
| | 5670:g141.t1 | P | AT2G37770.1 | | NAD(P)-linked oxidoreductase superfamily | | | |
| | 5670:g142.t1 | P | ATMG00860.1 | | DNA/RNA polymerases superfamily | | | |
| Scaffold ∅ | 8003:g143.t1 | R | | | | | | |
| | 8003:g144.t1 | R | AT4G23160.1 | CRK8 | cysteine-rich RLK | | | |
| | 8003:g145.t1 | M | | | | | | Retrovirus-related Pol polyprotein from transposon 17.6 [Vitis vinifera] |

[1] M: male-specific, P: hypothetical PAR, R: repetitive regions. Male-specific genes were highlighted in gray

[2] Male specific genes were annotated also with the nr database. The highest hit, excluding “hypothetical” or “uncharacterized” proteins, were given ($< 1e^{-10}$)

**Supplemental Table S8 Expression patterns of the genes predicted in the 9 scaffolds anchored by the Y-specific contigs.**

| Scaffold | Gene IDs | Gene name | Male specificity[1] | Expression levels (RPKM) | | | | | |
|---|---|---|---|---|---|---|---|---|---|
| | | | | Female deficient anther | Female whole flower | Female carpel | Male anther | Male whole flower | Male deficient carpel |
| Scaffold I | 6851:g1.t1 | | P | 0.0 | 0.0 | 0.1 | 0.0 | 0.0 | 0.0 |
| | 6851:g2.t1 | | P | 0.3 | 0.1 | 0.5 | 0.0 | 0.0 | 0.0 |
| | 6851:g3.t1 | | P | 0.1 | 0.2 | 0.1 | 0.0 | 0.0 | 0.0 |
| | 6851:g4.t1 | | P | 0.3 | 0.0 | 0.2 | 0.0 | 0.0 | 0.0 |
| | 6851:g5.t1 | | P | 1.4 | 0.6 | 1.7 | 2.3 | 0.0 | 0.0 |
| | 6851:g6.t1 | | P | 1.2 | 1.1 | 1.0 | 1.0 | 0.0 | 1.7 |
| | 6851:g7.t1 | | R | 3.7 | 3.5 | 4.7 | 6.2 | 1.5 | 5.6 |
| | 6851:g8.t1 | | R | 24.5 | 23.6 | 31.3 | 44.1 | 13.9 | 31.8 |
| Scaffold II | 8318:g9.t1 | | R | 11.3 | 9.9 | 15.3 | 19.5 | 4.4 | 16.6 |
| | 8318:g10.t1 | | R | 6.6 | 22.2 | 5.8 | 5.7 | 26.9 | 4.1 |
| | 8318:g11.t1 | | R | 1.7 | 2.4 | 1.6 | 3.5 | 1.4 | 2.2 |
| Scaffold III | 6898:g12.t1 | | R | 2.5 | 0.8 | 3.1 | 4.2 | 0.0 | 2.8 |
| | 6898:g13.t1 | | R | 2.0 | 1.3 | 1.3 | 1.2 | 1.5 | 1.4 |
| | 6898:g14.t1 | | P | 0.9 | 0.6 | 0.9 | 1.4 | 0.0 | 0.0 |
| | 6898:g15.t1 | | P | 0.6 | 0.4 | 0.6 | 0.0 | 0.0 | 0.0 |
| | 6898:g16.t1 | | P | 1.0 | 0.5 | 1.1 | 1.2 | 0.0 | 0.0 |
| | 6898:g17.t1 | | P | 0.0 | 0.2 | 0.2 | 0.0 | 0.0 | 0.0 |
| | 6898:g18.t1 | | P | 0.4 | 0.1 | 0.4 | 0.0 | 0.0 | 0.0 |
| | 6898:g19.t1 | | P | 0.1 | 0.1 | 0.2 | 0.0 | 0.0 | 0.0 |
| | 6898:g20.t1 | | P | 0.2 | 0.1 | 0.1 | 0.0 | 0.0 | 0.0 |
| | 6898:g21.t1 | | P | 0.0 | 0.1 | 0.0 | 0.0 | 0.0 | 0.0 |
| | 6898:g22.t1 | | P | 0.1 | 0.0 | 0.0 | 0.0 | 0.0 | 0.0 |
| | 6898:g23.t1 | | P | 0.2 | 0.2 | 0.1 | 0.0 | 0.0 | 0.0 |
| | 6898:g24.t1 | | R | 3.6 | 3.4 | 4.9 | 6.5 | 1.9 | 5.0 |
| | 6898:g25.t1 | | R | 4.5 | 6.0 | 7.3 | 10.7 | 3.1 | 7.5 |
| | 6898:g26.t1 | | P | 0.3 | 0.3 | 0.3 | 0.0 | 0.0 | 0.0 |
| Scaffold IV | 423383:g27.t1 | | R | 28.1 | 33.0 | 37.3 | 54.3 | 24.0 | 41.6 |
| | 423383:g28.t1 | | R | 9.2 | 8.6 | 15.9 | 21.8 | 4.8 | 17.0 |
| | 423383:g29.t1 | | R | 1.1 | 0.5 | 1.2 | 1.7 | 0.0 | 1.1 |
| | 423383:g30.t1 | | R | 0.7 | 0.6 | 1.2 | 2.0 | 0.0 | 1.2 |
| | 423383:g31.t1 | | R | 0.5 | 0.9 | 1.1 | 1.9 | 0.0 | 0.0 |
| | 423383:g32.t1 | | M | 0.0 | 0.0 | 0.0 | 0.0 | 0.0 | 0.0 |
| | 423383:g33.t1 | ***Shy Girl*** | **M** | **0.0** | **0.0** | **0.0** | **0.0** | **1.5** | **7.8** |
| | 423383:g34.t1 | | M | 0.1 | 0.0 | 0.1 | 0.0 | 0.0 | 0.0 |
| Scaffold V | 426301:g91.t1 | | R | 2.2 | 3.2 | 3.0 | 4.1 | 1.2 | 3.1 |
| | 426301:g90.t1 | | P | 0.0 | 0.0 | 0.0 | 0.0 | 0.0 | 0.0 |
| | 426301:g89.t1 | | M | 0.0 | 0.0 | 0.0 | 0.0 | 0.0 | 0.0 |
| | 426301:g88.t1 | | M | 0.1 | 0.1 | 0.1 | 0.0 | 0.0 | 0.0 |
| | 426301:g87.t1 | | M | 0.0 | 0.0 | 0.0 | 0.0 | 0.0 | 0.0 |
| | 426301:g86.t1 | | R | 3.2 | 7.2 | 0.3 | 2.4 | 5.0 | 1.2 |
| | 426301:g85.t1 | | M | 0.1 | 0.1 | 0.1 | 0.0 | 0.0 | 0.0 |
| | 426301:g84.t1 | | M | 0.0 | 0.0 | 0.0 | 0.0 | 0.0 | 0.0 |
| | 426301:g83.t1 | | R | 0.9 | 0.5 | 0.7 | 1.6 | 1.3 | 3.5 |
| | 426301:g82.t1 | ***YFT*** | M | 0.2 | 0.2 | 0.5 | 0.0 | 0.0 | 0.0 |
| | 426301:g81.t1 | | M | 0.3 | 0.4 | 0.5 | 0.0 | 0.0 | 0.0 |
| | 426301:g80.t1 | | M | 0.2 | 0.1 | 0.1 | 0.0 | 0.0 | 0.0 |
| | 426301:g79.t1 | | M | 0.2 | 0.2 | 1.2 | 0.0 | 0.0 | 0.0 |
| | 426301:g78.t1 | | R | 7.0 | 8.0 | 9.8 | 14.3 | 5.6 | 10.2 |
| | 426301:g77.t1 | | R | 0.5 | 0.7 | 0.7 | 1.1 | 0.0 | 1.1 |
| | 426301:g76.t1 | | R | 1.1 | 1.2 | 1.4 | 2.8 | 0.0 | 1.8 |
| | 426301:g75.t1 | | R | 1.9 | 1.6 | 2.3 | 3.6 | 1.1 | 1.6 |
| | 426301:g74.t1 | | M | 0.0 | 0.2 | 0.0 | 0.0 | 0.0 | 0.0 |
| | 426301:g73.t1 | | R | 0.2 | 0.1 | 0.0 | 0.0 | 0.0 | 0.0 |
| | 426301:g72.t1 | | M | 0.1 | 0.0 | 0.0 | 0.0 | 0.0 | 0.0 |
| | 426301:g71.t1 | | M | 0.1 | 0.1 | 0.4 | 0.0 | 0.0 | 0.0 |
| | 426301:g70.t1 | | M | 0.1 | 0.2 | 0.2 | 0.0 | 0.0 | 0.0 |
| | 426301:g69.t1 | | R | 7.7 | 10.4 | 12.3 | 18.1 | 6.5 | 12.9 |
| | 426301:g68.t1 | | R | 0.5 | 0.4 | 0.7 | 1.3 | 0.0 | 0.0 |
| | 426301:g67.t1 | | R | 8.8 | 10.4 | 12.3 | 16.3 | 7.4 | 11.9 |
| | 426301:g66.t1 | | R | 2.2 | 2.8 | 3.7 | 5.6 | 1.3 | 3.7 |
| | 426301:g65.t1 | | R | 3.5 | 4.0 | 4.1 | 5.9 | 1.7 | 4.0 |
| | 426301:g64.t1 | | P | 1.5 | 1.1 | 0.7 | 0.0 | 0.0 | 0.0 |
| | 426301:g63.t1 | | P | 0.1 | 0.1 | 0.1 | 0.0 | 0.0 | 0.0 |
| | 426301:g62.t1 | | R | 32.2 | 28.9 | 42.3 | 62.3 | 15.6 | 42.4 |
| | 426301:g61.t1 | | R | 28.1 | 22.9 | 44.1 | 70.0 | 13.5 | 50.8 |
| | 426301:g60.t1 | | R | 1.1 | 2.1 | 1.6 | 3.0 | 1.3 | 2.9 |
| | 426301:g59.t1 | | R | 0.4 | 0.3 | 0.8 | 1.4 | 0.0 | 0.0 |
| | 426301:g58.t1 | | M | 0.4 | 0.3 | 0.2 | 0.0 | 0.0 | 0.0 |
| | 426301:g57.t1 | | M | 0.3 | 0.1 | 0.1 | 0.0 | 0.0 | 0.0 |
| | 426301:g56.t1 | ***Friendly Boy*** | **M** | **0.0** | **0.0** | **0.0** | **2.1** | **0.0** | **0.8** |
| | 426301:g55.t1 | | R | 4.8 | 5.7 | 6.4 | 10.8 | 2.9 | 5.7 |

| | | | | | | | |
|---|---|---|---|---|---|---|---|
| | 426301:g54.t1 | R | 0.4 | 0.2 | 0.2 | 1.1 | 0.0 | 0.0 |
| | 426301:g53.t1 | R | 1.3 | 1.2 | 1.8 | 0.0 | 0.0 | 2.1 |
| | 426301:g52.t1 | R | 0.7 | 0.1 | 0.3 | 1.4 | 0.0 | 0.0 |
| | 426301:g51.t1 | M | 0.1 | 0.2 | 0.4 | 0.0 | 0.0 | 0.0 |
| | 426301:g50.t1 | R | 0.5 | 0.5 | 0.9 | 1.5 | 0.0 | 0.0 |
| | 426301:g49.t1 | R | 2.6 | 1.6 | 2.2 | 3.7 | 1.2 | 2.9 |
| | 426301:g48.t1 | R | 7.1 | 4.8 | 11.2 | 15.4 | 2.2 | 9.3 |
| | 426301:g47.t1 | R | 9.6 | 7.5 | 14.7 | 20.7 | 4.0 | 18.4 |
| | 426301:g46.t1 | M | 0.1 | 0.0 | 0.0 | 0.0 | 0.0 | 0.0 |
| Scaffold V | 426301:g45.t1 | M | 0.0 | 0.0 | 0.1 | 0.0 | 0.0 | 0.0 |
| | 426301:g44.t1 | M | 0.1 | 0.1 | 0.1 | 0.0 | 0.0 | 0.0 |
| | 426301:g43.t1 | R | 3.0 | 3.1 | 4.6 | 5.0 | 0.0 | 4.1 |
| | 426301:g42.t1 | R | 1.9 | 1.6 | 2.3 | 2.6 | 0.0 | 1.8 |
| | 426301:g41.t1 | R | 1.7 | 2.0 | 2.5 | 3.2 | 0.0 | 1.6 |
| | 426301:g40.t1 | P | 0.1 | 0.0 | 0.0 | 0.0 | 0.0 | 0.0 |
| | 426301:g39.t1 | P | 0.2 | 0.3 | 0.4 | 0.0 | 0.0 | 0.0 |
| | 426301:g38.t1 | P | 0.0 | 0.1 | 0.3 | 0.0 | 0.0 | 0.0 |
| | 426301:g37.t1 | P | 0.2 | 0.2 | 0.3 | 0.0 | 0.0 | 0.0 |
| | 426301:g36.t1 | R | 0.2 | 0.2 | 0.1 | 0.0 | 0.0 | 0.0 |
| | 426301:g35.t1 | R | 0.9 | 0.8 | 1.2 | 1.9 | 0.0 | 1.1 |
| | 7598:g92.t1 | P | 0.0 | 0.2 | 0.1 | 0.0 | 0.0 | 0.0 |
| | 7598:g93.t1 | R | 4.9 | 5.3 | 5.3 | 8.4 | 3.1 | 5.0 |
| | 7598:g94.t1 | P | 0.1 | 0.4 | 0.1 | 0.0 | 0.0 | 0.0 |
| | 7598:g95.t1 | R | 6.5 | 6.4 | 7.6 | 12.1 | 4.1 | 8.4 |
| | 7598:g96.t1 | P | 0.1 | 0.1 | 0.1 | 0.0 | 0.0 | 0.0 |
| | 7598:g97.t1 | P | 0.2 | 0.0 | 0.0 | 0.0 | 0.0 | 0.0 |
| | 7598:g98.t1 | P | 1.4 | 1.6 | 2.0 | 2.7 | 0.0 | 2.1 |
| | 7598:g99.t1 | M | 0.1 | 0.0 | 0.0 | 0.0 | 0.0 | 0.0 |
| | 7598:g100.t1 | M | 0.0 | 0.0 | 0.0 | 0.0 | 0.0 | 0.0 |
| | 7598:g101.t1 | R | 0.1 | 0.8 | 0.1 | 0.0 | 0.0 | 0.0 |
| | 7598:g102.t1 | M | 0.0 | 0.0 | 0.0 | 0.0 | 0.0 | 0.0 |
| | 7598:g103.t1 | M | 0.1 | 0.0 | 0.0 | 0.0 | 0.0 | 0.0 |
| | 7598:g104.t1 | R | 9.5 | 9.0 | 12.5 | 18.1 | 4.8 | 11.8 |
| | 7598:g105.t1 | R | 27.7 | 33.5 | 36.7 | 53.1 | 20.7 | 40.2 |
| | 7598:g106.t1 | R | 13.0 | 18.6 | 19.2 | 30.3 | 11.3 | 23.1 |
| Scaffold VI | 7598:g107.t1 | R | 13.9 | 17.5 | 20.8 | 29.8 | 9.6 | 21.1 |
| | 7598:g108.t1 | R | 5.4 | 6.8 | 6.6 | 9.4 | 4.4 | 6.9 |
| | 7598:g109.t1 | M | 0.0 | 0.0 | 0.0 | 0.0 | 0.0 | 0.0 |
| | 7598:g110.t1 | R | 3.2 | 2.7 | 3.7 | 2.7 | 1.1 | 3.3 |
| | 7598:g111.t1 | R | 1.4 | 1.4 | 2.8 | 2.2 | 0.0 | 4.0 |
| | 7598:g112.t1 | R | 5.3 | 4.8 | 7.2 | 9.4 | 2.2 | 7.4 |
| | 7598:g113.t1 | P | 0.7 | 0.0 | 0.5 | 0.0 | 0.0 | 0.0 |
| | 7598:g114.t1 | P | 0.2 | 0.2 | 0.3 | 0.0 | 0.0 | 0.0 |
| | 7598:g115.t1 | P | 0.1 | 0.1 | 0.1 | 0.0 | 0.0 | 0.0 |
| | 7598:g116.t1 | P | 0.0 | 0.1 | 0.0 | 0.0 | 0.0 | 0.0 |
| | 7598:g117.t1 | P | 0.1 | 0.0 | 0.4 | 0.0 | 0.0 | 0.0 |
| | g118 | P | 0.0 | 0.0 | 0.0 | 0.0 | 0.0 | 0.0 |
| | 7598:g119.t1 | P | 0.2 | 0.0 | 0.2 | 0.0 | 0.0 | 0.0 |
| | 7598:g120.t1 | P | 0.2 | 0.1 | 0.1 | 0.0 | 0.0 | 0.0 |
| | 7598:g121.t1 | R | 0.5 | 0.4 | 0.7 | 1.1 | 0.0 | 0.0 |
| | 7598:g122.t1 | R | 0.8 | 0.3 | 1.1 | 1.3 | 0.0 | 0.0 |
| | 7598:g123.t1 | P | 0.4 | 1.6 | 0.2 | 0.0 | 1.8 | 0.0 |
| | 422073:g124.t1 | R | 12.1 | 10.0 | 13.8 | 18.1 | 7.4 | 14.6 |
| | 422073:g125.t1 | M | 0.2 | 0.1 | 0.2 | 0.0 | 0.0 | 0.0 |
| | 422073:g126.t1 | R | 6.9 | 5.7 | 11.0 | 17.5 | 3.2 | 13.2 |
| Scaffold VII | 422073:g127.t1 | R | 4.2 | 3.6 | 5.9 | 8.3 | 2.3 | 6.6 |
| | 422073:g128.t1 | R | 4.3 | 4.5 | 6.8 | 10.0 | 2.3 | 6.9 |
| | 422073:g129.t1 | R | 1.4 | 1.4 | 1.5 | 2.3 | 1.0 | 1.7 |
| | 422073:g130.t1 | R | 1.1 | 1.2 | 1.6 | 2.8 | 0.0 | 1.3 |
| | 5670:g131.t1 | P | 0.0 | 0.1 | 0.3 | 0.0 | 0.0 | 0.0 |
| | 5670:g132.t1 | P | 1.8 | 2.3 | 2.1 | 1.2 | 1.3 | 1.1 |
| | 5670:g133.t1 | P | 0.5 | 0.0 | 0.0 | 0.0 | 0.0 | 0.0 |
| | 5670:g134.t1 | P | 0.0 | 0.0 | 0.0 | 0.0 | 0.0 | 0.0 |
| | 5670:g135.t1 | R | 1.0 | 0.9 | 1.5 | 1.9 | 0.0 | 1.1 |
| Scaffold VIII | 5670:g136.t1 | P | 0.9 | 0.8 | 0.6 | 1.9 | 0.0 | 0.0 |
| | 5670:g137.t1 | P | 1.6 | 0.8 | 1.6 | 2.2 | 0.0 | 0.0 |
| | 5670:g138.t1 | R | 15.7 | 17.2 | 24.1 | 36.3 | 10.5 | 24.5 |
| | 5670:g139.t1 | P | 3.1 | 6.3 | 2.4 | 1.2 | 9.4 | 4.1 |
| | 5670:g140.t1 | P | 2.9 | 4.3 | 4.1 | 4.7 | 3.8 | 4.5 |
| | 5670:g141.t1 | P | 11.0 | 14.9 | 8.5 | 5.0 | 21.4 | 10.7 |
| | 5670:g142.t1 | P | 0.3 | 0.2 | 0.3 | 0.0 | 0.0 | 0.0 |
| | 8003:g143.t1 | R | 6.8 | 6.4 | 9.5 | 13.2 | 3.5 | 9.9 |
| Scaffold ∅ | 8003:g144.t1 | R | 0.7 | 1.0 | 1.1 | 1.3 | 0.0 | 1.5 |
| | 8003:g145.t1 | M | 0.0 | 0.0 | 0.0 | 0.0 | 0.0 | 0.0 |

[1] M: male-specific, P: hypothetical PAR, R: repetitive regions. Male-specific genes were highlighted in gray

#### Supplemental Table S9 Pedigree of the KH line and the segregating population derived from the KH line

| selection | sex[1] | segregation test[2] | F parent | M parent | Cross | Progeny type | Genotype | Note |
|---|---|---|---|---|---|---|---|---|
| **Initial parents** | | | | | | | | |
| Hayward | F | | - | - | | | XX | |
| Matua | M | | - | - | | | XY | |
| M114 | IM | | - | - | | | XY | putative epimutant of Matua |
| **progenies derived from M114** | | | | | | | | |
| KFM | IM | | Hayward | M114 | F x IM | $F_1$ | XY | |
| KH | H | X | Hayward | M114 | F x IM | $F_1$ | $XY^h$ | |
| o12_05 | F | X | Hayward | KH | F x H | $BC_1F_1$ | XX | |
| o12_06 | F | X | Hayward | KH | F x H | $BC_1F_1$ | XX | |
| o12_01 | H | X | Hayward | KH | F x H | $BC_1F_1$ | $XY^h$ | |
| o12_02 | H | X | Hayward | KH | F x H | $BC_1F_1$ | $XY^h$ | |
| o12_03 | H | X | Hayward | KH | F x H | $BC_1F_1$ | $XY^h$ | |
| o14_02 | F | X | Hayward | o12_01 | F x H | $BC_2F_1$ | XX | |
| o14_03 | H | X | Hayward | o12_01 | F x H | $BC_2F_1$ | $XY^h$ | |
| o02_01 | H | X | KH | KFM | H x IM | $F_2$ | $XY^h$ | |
| o02_02 | H | X | KH | KFM | H x IM | $F_2$ | $XY^h$ | |
| o02_03 | H | X | KH | KFM | H x IM | $F_2$ | $XY^h$ | |
| o26_01 | H | X | Qinmei | o02_03 | F x H | $F_1$ | $XY^h$ | |
| o26_03 | H | X | Qinmei | o02_03 | F x H | $F_1$ | $XY^h$ | |
| s02_02 | H | X | KH | KH | H selfed | $S_1F_1$ | $XY^h$ | |

[1] F: female, M: male, IM: inconstant male, H: hermaphrodite

[2] used in assessment of co-segregation with a Y-marker (Supplemental Figure S12)

**Supplemental Table S10 Phenotypic characterization of p*FrBy-FrBy* induced kiwifruit.**

| line name | *pFrBy-FrBy* construct | fruit size[1] | seed (100-120 DPA)[2] | cross pollination[3] | pollen germination |
|---|---|---|---|---|---|
| 18-E7 | X | +++ | X | N/A | N/A |
| 18-E8 | X | +++ | X | N/A | X |
| 19-E1 | X | +++ | N/A | X | N/A |
| 19-E2 | X | +++ | X | X | N/A |
| 20-E6 | X | +++ | X | X | N/A |
| 21-E12 | X | +++ | X | N/A | X |
| 21-E13 | X | +++ | X | X | X |
| 19-E3 | X | ++ | N/A | N/A | N/A |
| 19-E4 | X | ++ | N/A | N/A | N/A |
| 21-E9 | X | ++ | N/A | N/A | X |
| 21-E11 | X | + | - | N/A | N/A |
| 20-E5 | X | + | - | N/A | N/A |
| cv. Hort16A (♀) | - | + | - | - | - |
| cv. Bruce (♂) | - | - | - | X | X |

[1]Self-pollination

[2]Black seed observed

[3]Development of large fruit bearing fertile seed in cross pollination with rapid flowering cv. Hort16A

**Supplemental Table S11: List of plant materials.**

| spp/line | accession name | gender | ploidy | application[1] | location |
|---|---|---|---|---|---|
| KE population (*Actinidia rufa x A. chinensis*) | Ke2 | female | 2X | MY | Kagawa Univ. Japan |
| | Ke4 | female | 2X | MY | Kagawa Univ. Japan |
| | Ke5 | female | 2X | MY | Kagawa Univ. Japan |
| | Ke14 | female | 2X | AT, MY | Kagawa Univ. Japan |
| | Ke15 | female | 2X | AT, MY | Kagawa Univ. Japan |
| | Ke16 | female | 2X | MY | Kagawa Univ. Japan |
| | Ke19 | female | 2X | MY | Kagawa Univ. Japan |
| | Ke20 | female | 2X | MY | Kagawa Univ. Japan |
| | Ke24 | female | 2X | AT, MY | Kagawa Univ. Japan |
| | Ke25 | female | 2X | MY | Kagawa Univ. Japan |
| | Ke29 | female | 2X | MY | Kagawa Univ. Japan |
| | Ke31 | female | 2X | AT, MY | Kagawa Univ. Japan |
| | Ke32 | female | 2X | AT, MY | Kagawa Univ. Japan |
| | Ke33 | female | 2X | MY | Kagawa Univ. Japan |
| | Ke35 | female | 2X | MY | Kagawa Univ. Japan |
| | Ke36 | female | 2X | MY | Kagawa Univ. Japan |
| | Ke42 | female | 2X | MY | Kagawa Univ. Japan |
| | Ke46 | female | 2X | MY | Kagawa Univ. Japan |
| | Ke48 | female | 2X | MY | Kagawa Univ. Japan |
| | Ke51 | female | 2X | MY | Kagawa Univ. Japan |
| | Ke3 | male | 2X | AT, MY | Kagawa Univ. Japan |
| | Ke6 | male | 2X | MY | Kagawa Univ. Japan |
| | Ke7 | male | 2X | MY | Kagawa Univ. Japan |
| | Ke9 | male | 2X | MY | Kagawa Univ. Japan |
| | Ke10 | male | 2X | MY | Kagawa Univ. Japan |
| | Ke11 | male | 2X | MY | Kagawa Univ. Japan |
| | Ke12 | male | 2X | MY | Kagawa Univ. Japan |
| | Ke13 | male | 2X | MY | Kagawa Univ. Japan |
| | Ke17 | male | 2X | MY | Kagawa Univ. Japan |
| | Ke21 | male | 2X | MY | Kagawa Univ. Japan |
| | Ke22 | male | 2X | AT, MY | Kagawa Univ. Japan |
| | Ke23 | male | 2X | AT, MY | Kagawa Univ. Japan |
| | Ke26 | male | 2X | MY | Kagawa Univ. Japan |
| | Ke27 | male | 2X | AT, MY | Kagawa Univ. Japan |
| | Ke28 | male | 2X | AT, MY | Kagawa Univ. Japan |
| | Ke30 | male | 2X | MY | Kagawa Univ. Japan |
| | Ke34 | male | 2X | MY | Kagawa Univ. Japan |
| | Ke37 | male | 2X | MY | Kagawa Univ. Japan |
| | Ke39 | male | 2X | MY | Kagawa Univ. Japan |
| | Ke45 | male | 2X | MY | Kagawa Univ. Japan |
| | Ke52 | male | 2X | MY | Kagawa Univ. Japan |
| | Ke53 | male | 2X | MY | Kagawa Univ. Japan |
| *A. chinensis* | FCM1 | male | 2X | MY | Kagawa Univ. Japan |
| | Soyu | male | 2X | GA, MY | Kagawa Univ. Japan |
| *A. rufa* | Fuchu | female | 2X | MY | Kagawa Univ. Japan |
| *A. deliciosa* | Hayward | female | 6X | HT, LCM, ISH | Kyoto Univ. Japan |
| | KH[2] | hermaphrodite | 6X | HT | Plant & Food Research, Nea Zealand |
| | Tomuri | male | 6X | LCM, ISH | Kyoto Univ. Japan |
| | Matsua | male | 6X | HT | Kagawa Univ. Japan |

[1]AT: anther transcriptome, GA: genome assembly, HT: hermaphrodite testing, ISH: *in situ* RNA hybridization, MY: mapping to Y chromosome

[2]Information of the segregating population related KH was given in Table S9

**Supplemental Table S12: List of primers used in this study.**

| Primer name | Sequences (5-3) | Target | Note |
|---|---|---|---|
| **For gene-editing** | | | |
| AT1G30800_ge3_F | ATTGACGCCGTCGAGTGCGTAACC | gRNA for *AT1G30800* in *Arabidopsis* | |
| AT1G30800_ge3_R | AAACGGTTACGCACTCGACGGCGT | | |
| M_oligo3_F | ATTGTCAAATCATCAAACACC | gRNA for *FAS1* in *Nicotiana tabacum* | |
| M_oligo3_R | AAACGGTGTTTGATGATTTGA | | |
| pPVL02_AtU6_infBamF | AACTCCATAAGGATCCCGACGTCGCATGCTCCCG | | To make pPLV2-GE cassett |
| pPVL02_Hsp_infBamR | ATCGGGGATCGGATCCACTAGTGATATCACCACTTTG | | |
| kiwi-YFAS-prom-F-LIC2 | TTCTAGTTGGAATGGGTTACATAGTGGTAGGCAATTGTTAATGGT | *FrBy* under the native promoter | To make pPLV2-pFrBy-FrBy |
| kiwi-YFAS-R-wStp-LIC2 | TCCTTATGGAGTTGGGTTTCATTAACAAACCCAAACCCTAAAATAAAC | | |
| AT1G30800_F | ATGGCCACCTCAAGCCAT | *AT1G30800* genotyping | |
| AT1G30800_R | TCAGAACAGAGCCAGAGTCG | | |
| NtFAS_check_F | CCCAACACCACCACATTT | *FAS1* in *N. tabacum* genotyping | |
| NtFAS_check_R | TGGAAGGTAAGAGAGTTGGG | | |
| **For PCR** | | | |
| FrBy-F1 | ATGGCAAAGTGGTTCTCTCTCCAT | full length of *FrBy* | designed from *A. chinensis* |
| FrBy-R1 | TTAACAAACCCAAACCCTAAAATAAAC | | |
| FrBy-F2 | ATGCTGCCCTCTCTCACTTT | partial *FrBy* | For *FrBy* RNA probe |
| FrBy-R2 | GGATGGGTGATAGGAGAGGC | | |
| FrBy-RTPCR-F1 | AACCAACACTCGCCTTCCCA | partial *FrBy* | |
| FrBy-RTPCR-R1 | GGATGGGTGATAGGAGAGGCATC | | |
| SyGI-spe-ORF-F | TGAAAGTAACAAAAATGG**T**CCG | partial *SyGI* | Underlined "T" in bold is originally "C". Applicable only in *A. chinensis* and *A. rufa* |
| SyGI-commonRR-spR | TCAATATTGGTACTTGATGTTGAGTTGG | | |
| Achn-act1-F | ATGGCCGATGCTGAGGATATTCAG | *Actinidia actin, Achn107351* | |
| Achn-act1-R | ACACCTGTGTGCCTGGGTCGA | | |
